# Structure of the Inmazeb cocktail and resistance to escape against Ebola virus

**DOI:** 10.1101/2022.10.11.511805

**Authors:** Vamseedhar Rayaprolu, Ben Fulton, Ashique Rafique, Emilia Arturo, Dewight Williams, Chitra Hariharan, Heather Callaway, Amar Parvate, Sharon L. Schendel, Diptiben Parekh, Sean Hui, Kelly Shaffer, Kristen Pascal, Elzbieta Wloga, Stephanie Giordano, Richard Copin, Matthew Franklin, RuthMabel Boytz, Callie Donahue, Robert Davey, Alina Baum, Christos A. Kyratsous, Erica Ollmann Saphire

**Affiliations:** La Jolla Institute for Immunology, La Jolla, CA 92037; Regeneron Pharmaceuticals, Tarrytown, NY 10591; Eyring Materials Center, Arizona State University, Tempe, AZ 85281; Dept. of Medicine, University of California, San Diego, CA 92037; Department of Microbiology, Boston University of Medicine and NEIDL, Boston University, Boston, MA 02118; Pacific Northwest Center for CryoEM, Portland, OR 97225; Environmental Molecular Sciences Laboratory, Pacific Northwest National Laboratory, Richland, WA 99354; Washington University School of Medicine, St. Louis, Missouri, MO 63110; Division of Biology and Biological Engineering, California Institute of Technology, Pasadena, CA 91125; American Association of Immunologists, Rockville, MD 20852

## Abstract

Monoclonal antibodies can provide important pre- or post-exposure protection against disease for those not yet vaccinated or in individuals that fail to mount a protective immune response after vaccination. A key concern in use of monotherapy monoclonal antibody products lies in the high risk of mutagenic escape. Inmazeb (REGN-EB3), a three-antibody cocktail against Ebola virus, demonstrated efficacy in lessening disease course and improving survival in a randomized, controlled trial. Here we present the cryoEM structure at 3.1 Å of the Ebola virus glycoprotein, determined without symmetry averaging, in a simultaneous complex with eight Fab fragments of antibodies in the Inmazeb cocktail. This structure allows modeling of previously disordered portions of the glycan cap, maps the non-overlapping epitopes of Inmazeb, and illuminates the basis for complementary activities, as well as residues that are critical for resistance to escape by each component of this cocktail and other clinically relevant antibodies. We also provide direct evidence that, unlike monotherapy treatments, including those targeting conserved epitopes, the Inmazeb protects against the rapid emergence of EBOV escape mutants and supports the benefit of the combination approach.

## Introduction

Nearly 40 outbreaks of Ebola Virus Disease (EVD) have occurred, including sustained outbreaks in 2014-2016 of over 28,000 cases and 11,000 deaths, another in 2018 involving 3,300 cases and 2,200 deaths, and a re-emergence in June 2020 that may be linked to recrudescent virus from a survivor of an earlier outbreak (Pratt, 2020). Although an effective vaccine against Ebola virus (EBOV) is available, there are challenges associated with widespread vaccination and there is risk of breakthrough cases in vaccinated individuals (Mulangu et al., 2019). Given the exceptionally high mortality rate, monoclonal antibody (mAb)-based therapeutics remain important for treating patients infected with EBOV. While several mAb therapies are in development to treat EBOV infections, REGN-EB3 (Inmazeb) is the first FDA-approved therapeutic for treatment of EBOV infection, followed by mAb114 (Ansuvimab) the second (<u>link</u>).

Therapeutic antibodies against EBOV target the viral glycoprotein (GP). EBOV GP is the only viral protein expressed on the virion surface and is required for host cell attachment, endosomal entry and membrane fusion. In the producer cell, the GP precursor is cleaved by host furin to yield GP1 and GP2 subunits that remain linked by a disulfide bond (Sanchez et al., 1998). Three GP1-GP2 heterodimers associate and are displayed as trimers on the virus surface (Lee et al., 2008). The GP1 subunit drives attachment to target cells and includes the receptor-binding site (RBS), a glycan cap that shields the receptor-binding site, and a mucin-like domain (MLD). The GP2 subunit includes a transmembrane domain (TM) that anchors GP in the viral membrane, a stalk, and an internal fusion loop (IFL) that promotes fusion of the virus with target cell membranes. After macropinocytosis-driven internalization of virions into target cells, endosomal cathepsins process GP, remove the glycan cap and MLD to yield a cleaved version of GP, termed GP_CL,_ which can bind the intracellular receptor Niemann-Pick C1 (NPC1) (Wang et al., 2016). Following receptor binding, GP2 rearranges into a six-helix bundle that drives membrane fusion (Gregory et al., 2011). In addition to membrane-bound GP, infected cells also produce secreted GP (sGP) that carries the N-terminal 295 amino acids of GP, but lacks both the GP2 and MLD (Sanchez et al., 1998). sGP is the primary product of the GP gene and may elicit production of non-neutralizing antibodies that could dampen the effectiveness of the immune response, but its function remains unclear (Ito et al., 2001)

Antibodies against GP have been shown to target sites across the entire molecule. Epitopes are found in the membrane-proximal stalk domain, the GP2 fusion loop, the base which includes regions of both GP1 and GP2, and in GP1, the receptor-binding head, glycan cap and MLD. Multiple studies, including a parallel analysis by an international consortium (Saphire et al., 2018a), have determined that a combination of antibody functions correlates with in vivo protection, including both neutralization (measured by blocking of infection in cell culture), and induction of multiple immune ‘effector’ functions by the antibody Fc. Some antibodies only have one antiviral function, whereas others, such as those that bind the GP1 head domain, can achieve both neutralization and Fc-mediated activities (Saphire et al., 2018a)(Gunn et al., 2018). The combination of neutralization and effector functions are also achieved, for example, in the ZMapp and rEBOV-520 plus rEBOV-548 cocktails (Gilchuk et al., 2020; Qiu et al., 2014).

The REGN-EB3 (Inmazeb) cocktail contains three fully human mAbs (Pascal et al., 2018) that are each directed against a unique, non-overlapping epitope on EBOV GP. The three antibodies in the REGN-EB3 cocktail were selected based on their ability to simultaneously bind EBOV GP and on their complementary combination of functional properties. REGN3479 (maftivimab), targets the fusion loop and is potently neutralizing (Pascal et al., 2018). REGN3471 (odesivimab), targets the GP1 head and sGP. Although REGN3471 is poorly neutralizing, it mediates effector function and is partially protective in a guinea pig model of EBOV infection. REGN3470 (atoltivimab), targets the glycan cap, is partially neutralizing, and also mediates Fc effector functions that can promote killing of EBOV-infected cells. During selection and development of these antibodies, it was hypothesized that a three-antibody cocktail may also reduce the potential for selection of antibody-resistance by requiring the simultaneous selection of escape mutations in the GP to each component (Pascal et al., 2018)

The therapeutic efficacy of REGN-EB 3 cocktail was evaluated in the PALM trial (Pamoja Tulinde Maisha-“Together We Save Lives”) a randomized, controlled trial conducted in the Democratic Republic of the Congo (DRC) during the 2018 EBOV outbreak. This trial compared the efficacy of the triple mAb cocktail ZMapp (control group) with the antiviral agent Remdesivir, the single antibody mAb114, and the REGN-EB3 triple mAb cocktail (Mulangu et al., 2019). The study endpoint was survival at 28 days post-treatment. Patients treated with mAb114 or REGN-EB3 were substantially less likely to die than those treated with ZMapp (percentage surviving: 35.1% and 33.5%, respectively, vs. 51.3%). In fact, the significantly higher efficacy of REGN-EB3 prompted the early termination of the clinical trial after reviewing 499 of the 700 patients enrolled. These results provided the first clinically-evaluated, specific treatment options for EBOV infection. Based on the PALM study results, REGN-EB3 was granted FDA approval as the first antiviral mAb cocktail for treatment of EVD, the first mAb product for treatment of a viral infection, and only the second mAb product against a viral antigen (De Clercq and Li, 2016).

The lack of high-resolution structure for the REGN-EB3 cocktail in complex with EBOV GP complicates understanding of the precise contacts that mediate its functions and its complementary activities. Mapping the epitope footprints of these three antibodies can facilitate monitoring of clinical isolates for GP that carry mutations that are resistant to REGN-EB3. Indeed, the ability of EBOV GP to acquire mutations and develop drug-induced resistance to approved mAb therapeutics has not been previously evaluated. The relevance of in vitro viral escape studies with SARS-CoV-2 have now been confirmed in the clinical setting, with demonstration that monotherapy treatment can select drug-resistant viruses in treated individuals (Copin et al. 2021; Rockett et al. 2021; GSK)

Here we report a 3.1Å resolution cryoEM structure of the complete REGN-EB3 cocktail simultaneously bound to the EBOV GP trimer. This structure maps the non-overlapping epitopes of REGN-EB3, and illuminates the basis for complementary activities, as well as residues that are critical for resistance to escape by each component of this cocktail and other clinically relevant antibodies. We also provide direct evidence that, unlike monotherapy treatments, including those targeting conserved epitopes, the REGN-EB3 combination protects against the rapid emergence of EBOV escape mutants and supports the benefit of the combination approach, as has been extensively demonstrated for SARS-CoV-2 (Baum et al., 2020; Copin et al., 2021).

## Results

To define the structural basis of binding, neutralization and protection by the REGN-EB3 cocktail of antibodies, we first carried out cryo-EM analysis of Fabs of the REGN-EB3 cocktail in complex with recombinant, fully glycosylated EBOV GP that lacks the MLD (Mayinga strain, Figure 1A). In initial 2D class averages, complexes were fully occupied with nine Fab fragments (i.e., three copies of each of the three different antibodies). In 3D classes, some particles had nine Fab fragments bound, whereas others had eight Fab fragments bound (three each of REGN3471 and REGN3479 plus two copies of REGN3470). Homogeneous and non-uniform refinements using cryoSPARC (Punjani et al., 2017) yielded a cryo-EM reconstruction of the complex resolved to ∼3.1Å. Models of the GP and the Fab Fv only were docked into the density and refined (Figures 1B-C, S1-2, and Table S1-2). We determined this structure using C1 symmetry so that individual differences among the monomers can be discerned. All previous cryoEM structures of EBOV GP at this resolution were determined using C3 symmetry in which the three monomers in the trimer were averaged around the three-fold axis. There is typically a trade-off between resolution and examination of asymmetry: resolution is improved by averaging monomers together, but biologically relevant individual asymmetric elements are lost during the computational process. For this structure, application of C3 symmetry and averaging resulted in a loss of signal for all three Fv regions of REGN3470 to the extent that precluded unambiguous model building. In contrast, the C1 symmetry maps revealed clear density for one copy of REGN3470 and moderately strong density for a second copy, which allowed us to build two copies of this antibody into the final structure.

**Figure 1:**
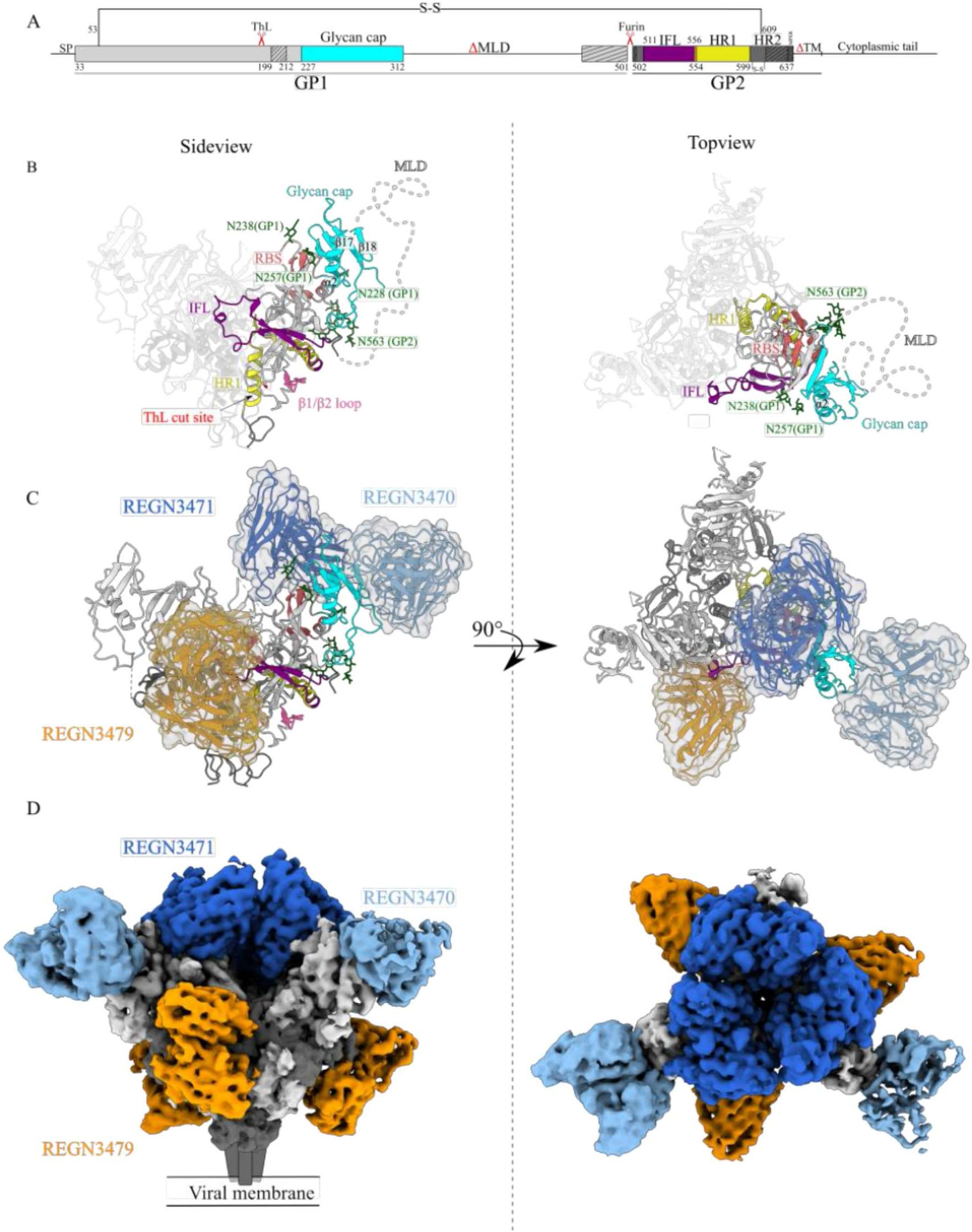
A) Schematic representation of EBOV GP ectodomain. The mucin-like domain (MLD) and transmembrane (TM) domains are deleted. Thermolysin (ThL) and furin cleavage sites are indicated. A disulfide bond links the N-terminal portion of GP1 (light grey) to GP2 (dark grey). The glycan cap, internal fusion loop (IFL) and heptad repeats 1 (HR1) are colored cyan, purple and yellow, respectively. Hash-marked regions in the schematic correspond to disordered regions in the structure B) Cartoon representation of side and top views of the EBOV GP trimer in the complex. The dotted line indicates the approximate region of the MLD. The β1-β2 loop (pink), receptor binding site (RBS; brick red), ThL cut site (red), glycan cap (cyan), mucin-like domain (MLD; white), internal fusion loop (IFL; purple), heptad repeat 1 (HR1; yellow) and glycans (green) are shown. C) Cartoon representation of EBOV GP trimer bound to a single Fv each of the three different antibodies illustrated for clarity (side view and top view). Cryo-EM maps and cartoon representations of REGN3470, REGN3471 and REGN3479. D) The 3.1Å cryoEM map of the EBOV-GP bound to eight Fabs of the REGN-EB3 cocktail. GP1 and GP2 are light and dark grey, respectively. In both (C) and (D) REGN3470, REGN3471 and REGN3479 are colored light blue, dark blue and orange, respectively.

### REGN3479 Fab recognizes a quaternary epitope

The REGN3479 Fab binds a quaternary epitope that bridges the fusion loop, which forms a paddle-shaped region of hydrophobicity (West et al., 2018)), of one GP monomer, monomer A, to a GP1/GP2-containing site in an adjacent monomer, monomer B (Figure 2A). Within this footprint, REGN3479 binds residues 527-530, 535 and 536 of the fusion loop paddle of monomer A, as well as a conformational site involving three sections of monomer B and a glycan. This conformation site comprises GP1 residues Pro34, Leu43, and Val45 (β1-β2 strands); GP2 N-terminal residues Ile504, Val 505, Ala507; GP2 HR1_A_ residues Gly 560, Glu564, and Gln567; and the glycan attached to Asn563 (Figure 2B-E). Leu 529 at the tip of the fusion loop was previously shown to bend inward to form a hydrophobic bridge with Ile544.

**Figure 2:**
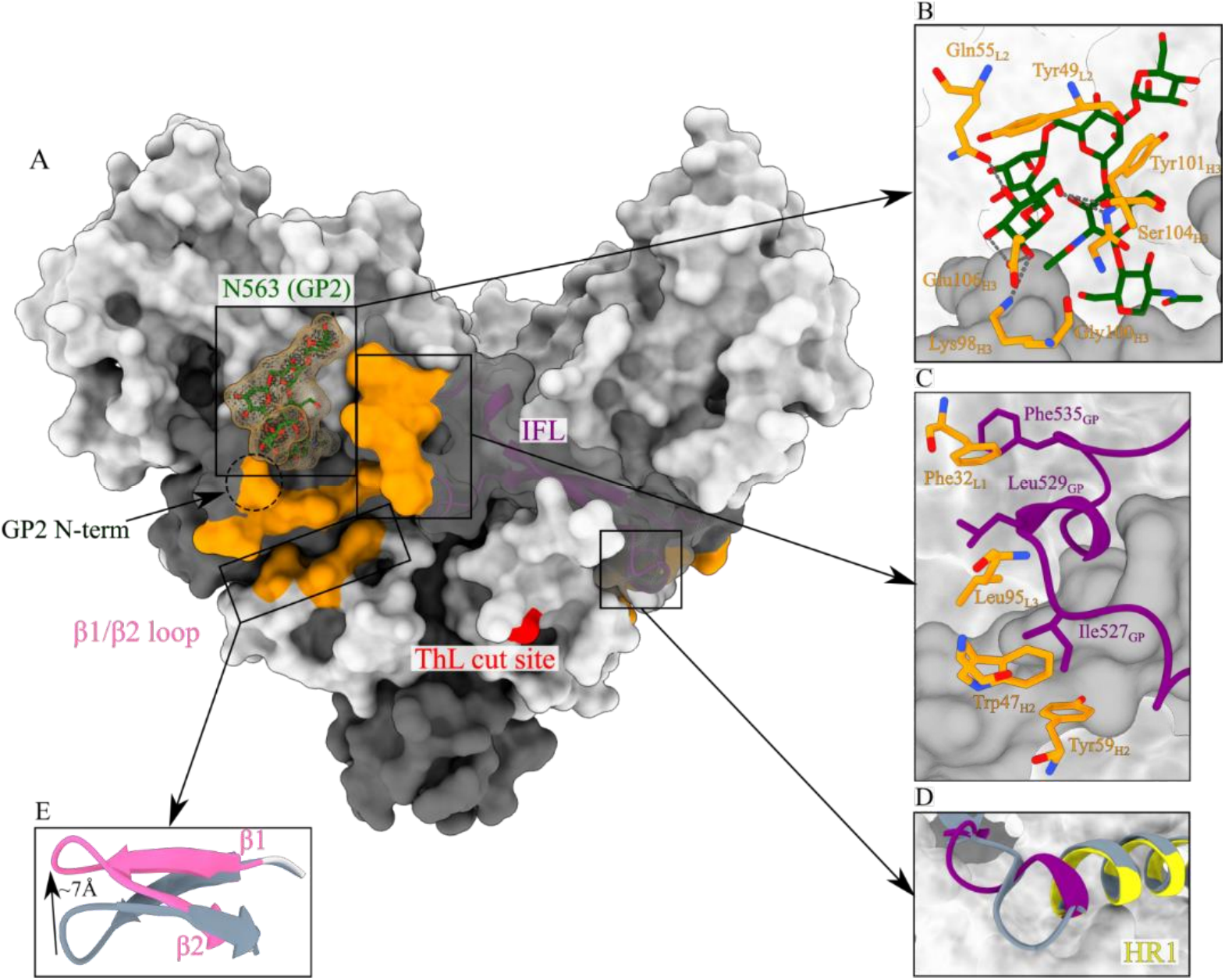
REGN3479 targets a quaternary epitope on the GP trimer. A) Surface representation of the GP trimer with the REGN3479 epitope footprint colored in orange, Glycan moieties are in green and the thermolysin cleavage site is in red. B) REGN3479 residues (orange) interact extensively with the Asn563 glycan (green). C) REGN3479 interactions (orange) with the fusion loop (purple). LC residue Phe32 wedges into the fusion loop tip. D) Overlay of GP in the unbound (gray, PDB ID:5JQ3) and REGN3479-bound form (purple) with the HR1 shown as a gray (unbound) and yellow (bound), respectively, ribbon diagram. Antibody binding introduces an additional helical turn in the HR1 N-terminus (purple). E) The β1/β2 loop shifts by 7Å in the unbound (gray) to bound (pink) shows a ∼7Å shift mediated by REGN3479 binding.

Phe535 interacts with this bridge to form a hydrophobic patch that is required for formation of a fusion-competent fist-like structure (Gregory et al., 2014). Phe32 in the REGN3479 light chain (LC) disrupts formation of this hydrophobic patch formation by wedging between Leu529 and Phe535 of monomer 1 in the GP trimer (Figure 2C). REGN3479 binding introduces other conformational adjustments on GP including a ∼180° rotation of Ile527 in the fusion loop paddle into a hydrophobic pocket created by the heavy chain (HC) and LC residues Trp47 and Tyr59, respectively. REGN 3479 binding also induces translation of 277-280 of the β17 strand 6Å inwards towards the base of GP helix α2. Furthermore, residues 549-553 fusion loop stem C-terminus translate upon REGN3479 binding and residues 552-553 lift to contribute to a helical extension of HRlA (Figure 2D).

A primary interaction of REGN3479 Fab is with the conserved GP-linked glycan on Asn563 on the monomer B. Tyr49 in REGN3479 LC forms CH-pi stacking interactions with two α-mannose moieties, whereas HC residue TyrlOl stacks with the βmannose. The main-chain atoms of LC residues Tyr49 and Gln55 and HC residues GlylO0 and TyrlOl, as well as the side-chain atoms of Lys98, Serl04 and Glul06 in the HC sandwich the GP glycan in a network of hydrophobic interactions, hydrogen bonds and salt bridges (Figure 2B, Table S3). This glycan has been shown to reduce GP processing and incorporation (Wang et al., 2017). Mutations in this glycan position have been shown to disrupt crucial GP conformations required for infection (Lennemann et al., 2015). High levels of GP on the viral surface reduce titers and therefore are undesirable (Mohan et al., 2015). Shielding of Gln563 by REGN3479 may enhance surface levels of GP on the virus and thereby reduce viral titers.

### REGN3471 binds both the head and glycan cap of GP

The epitope footprint of REGN3471 Fab includes both the receptor-binding head (60%) and glycan cap (40%) of GP1 (Figure 1C). GP residues 111-118 that are critical for binding to the cellular NPC1 receptor, nestle between the HC and LC of REGN3471, whereas GP residues 143, 144 and 146 interact with all three CDRs of the HC (Figure 3A). The extended CDR_L1_ of REGN3471 reaches ∼10Å into the receptor-binding site to form hydrogen bonds with GP Glu112. Meanwhile, the CDR_L2_ interacts with the glycan linked to Asn238 of GP as well as Thr270 and Lys 272. This interaction shifts Thr270 by ∼9Å towards the β18 strand (Figure 3B-C, Table S3). Alanine scanning of GP previously demonstrated that both Thr270 and Lys272 are critical residues involved in binding of antibodies that target the GP head (Davidson et al., 2015). The partial protection (∼33%) offered by REGN3471 alone is most likely due to the inhibition of receptor binding, as well as Fc effector functions.

**Figure 3:**
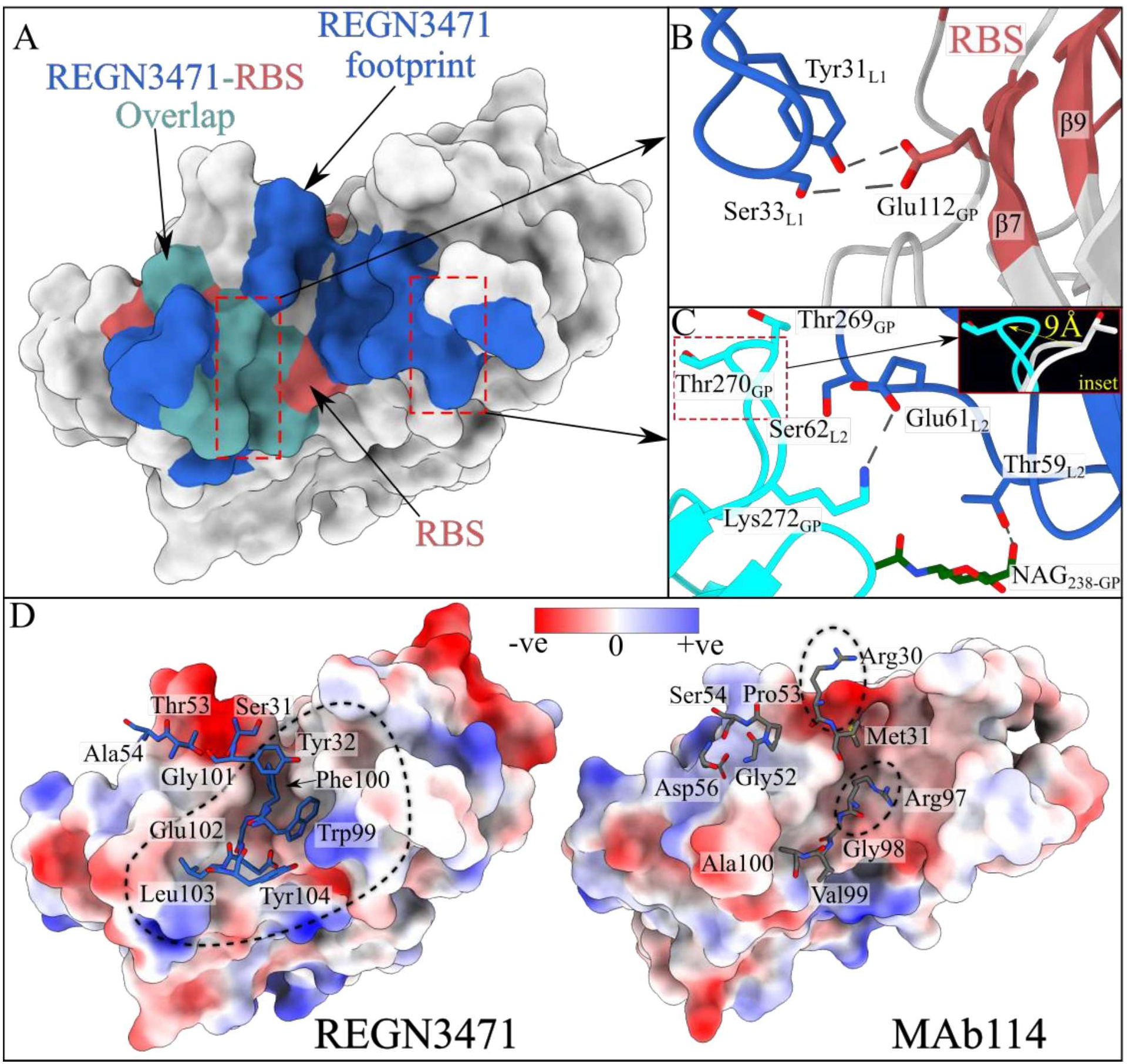
Interaction of REGN3471 with EBOV GP receptor-binding site (RBS). A) Top view of REGN3471 footprint shown in blue, receptor-binding site (RBS) in brick red. Residues that overlap between the REGN3471 footprint and RBS shown in teal. B) REGN3471 CDRLl forms hydrogen bonds with residues in the hydrophilic, receptor binding crest of GP. C) Salt-bridge interaction between Lys 272 of GP and Glu61 of REGN3471 enhances antibody binding affinity. The inset shows the overlay of GP in the unbound (gray, PDB ID:5JQ3) and bound (cyan) form. Steric clashes with Ser62 of the Fab displace the GP Thr270 by ∼9Å. These steric clashes and Thr270 loop movement are not seen with mAb114 binding. D) Electrostatic maps of GP bound to REGN3471 (left) or mAb114 (right). Antibody residues unique to REGN3471 or mAb114 are labeled. Several hydrophobic residues in REGN3471 occupy the hydrophilic crest of GP. Although resolution of the mAb114 complex is low, stronger and more favorable hydrophilic residues of mAb114 relative to REGN3471 interact with the GP crest to form high-affinity salt bridges and also participate in other interactions.

### REGN3470 stabilizes the β71IH8 loop in the glycan cap

REGN3470 binds to the top of the glycan cap of GP1 (Figure 1C), at a site involving the antiparallel β17 and β18 strands, which are connected by a 27-residue descending loop (residues 279-306) (Figure 2A). Notably, this structure of the GP-REGN3470 complex allows the first visualization of the entire β17/18 loop (Figure 4A), which was disordered in all previous structures of EBOV GP. In complex with the REGN3470 Fab, the β17/18 loop of GP is fully ordered, and the main chain direction as well as the identity and position of the side chains are clear. Interestingly, this clarity occurs only for the two GP monomers that are in complex with the REGN3470 Fab. In the remaining, unbound GP monomer, residues 262-270, 280-282 and 293-311 are disordered, similar to previous structures.

**Figure 4:**
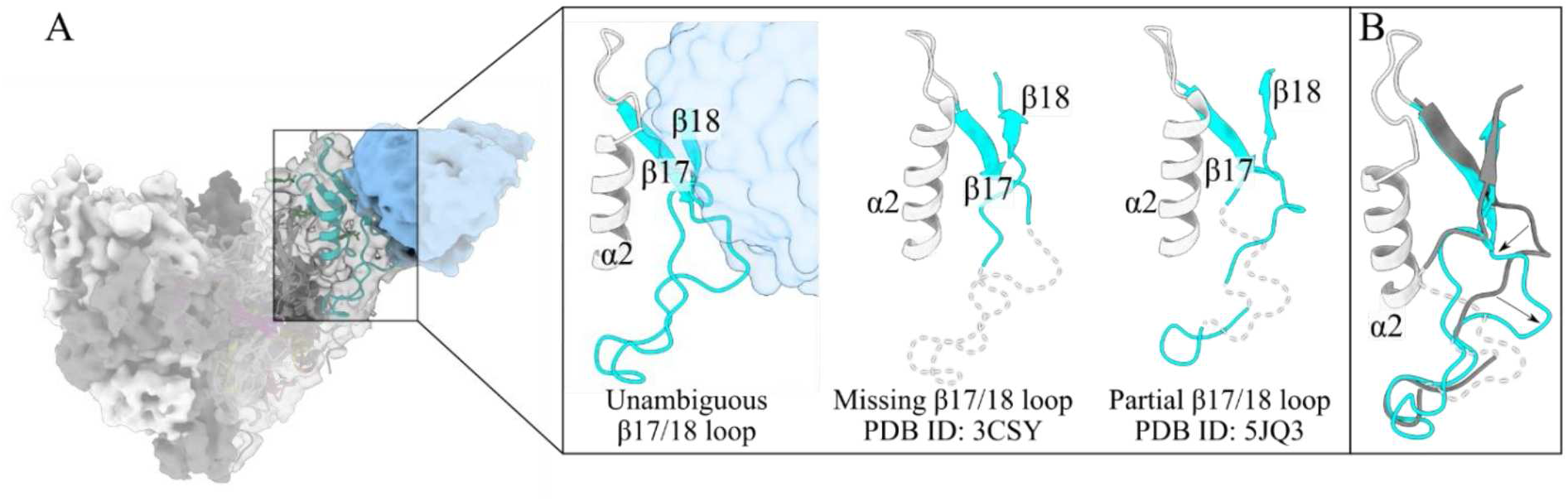
A) CryoEM map ofREGN3470 bound to EBOV GP. Inset compares the β17/18 loop in REGN3470 bound to GP and two previously published structures (PDB ID:3CSY and 5JQ3). Unambiguous density of this loop is observed in the REGN3470 complex. B) Comparison of unbound GP (PDB ID: 5JQ3, gray) with GP bound to REGN3470 shows movement of the loop upon REGN3470 binding.

The β17 and β18 strands of the GP glycan cap are bound by 19 residues of REGN3470, 10 of which interact with Asn278, Glu280 and Asp282 (Table S3). The increased order of the loop connecting the β17 and β18 strands in the REGN-EB3 complex likely arises from binding of bulky and polar antibody residues in the antibody HC including Asn31, Tyr32, His53, and Asn99. This binding consequentially drives GP residues 277-280 of the β17 strand ∼6Å inwards towards the base of GP helix a2. The ordering of the loop creates a new intra-GP backbone interaction between Ile285 and Arg247 that causes the intervening GP residues 281-285 to buckle outward, bringing the entire loop encompassing residues 285-312 into order (Figure 4B). Elsewhere in the glycan cap, LC residues Ser30 and Tyr32 interact with GP residue Ser263 to unravel half a helix turn from the C terminus of α2 and change the position of the loop between the α2 helix and the β17 strand.

Notably, GP residues bound by REGN3470 lie entirely within the glycan cap, which is removed following cleavage in the endosome by cathepsins B and L (Brecher et al., 2012; Chandran et al., 2005), or other endosomal proteases (Marzi et al., 2012). Removal of the REGN3470 epitope during viral entry could explain why REGN3470 is a less potent neutralizer but does elicit immune effector functions (Pascal et al., 2018) that require binding to intact GP displayed on cell surfaces.

### Glycan cap removal abolishes REGN3470 and REGN3471 GP binding

Following macropinocytosis, EBOV-GP is cleaved by cathepsins at the lower pH in the endosome. This cleavage event removes epitopes contained in the glycan cap and the MLD to produce cleaved GP (GPcL) in which the receptor-binding site (RBS) is exposed to allow binding to the NPCl receptor. To mimic cathepsin B cleavage in vitro, we treated GP with thermolysin (ThL) at a molar ratio of 60:1 GP:ThL (Misasi et al., 2012) and then used surface plasmon resonance (SPR) to assess the binding affinity and kinetics of the three antibodies of REGN-EB3 immobilized on the chip surface for GP and GPcL. REGN3479 had high affinity for GP as evidenced by a Ko of 1.64nM increased by over IO-fold for GPcL (0.14nM). REGN3471 had a four-fold reduction in affinity for GPcL relative to GP (4.4nM vs. 15.8nM). In addition to the reduced affinity, the binding half-life for REGN3471 was reduced by 4-fold. REGN3470 showed no binding to the GPcL, as expected (Figure 5A and Figure S3).

**Figure 5:**
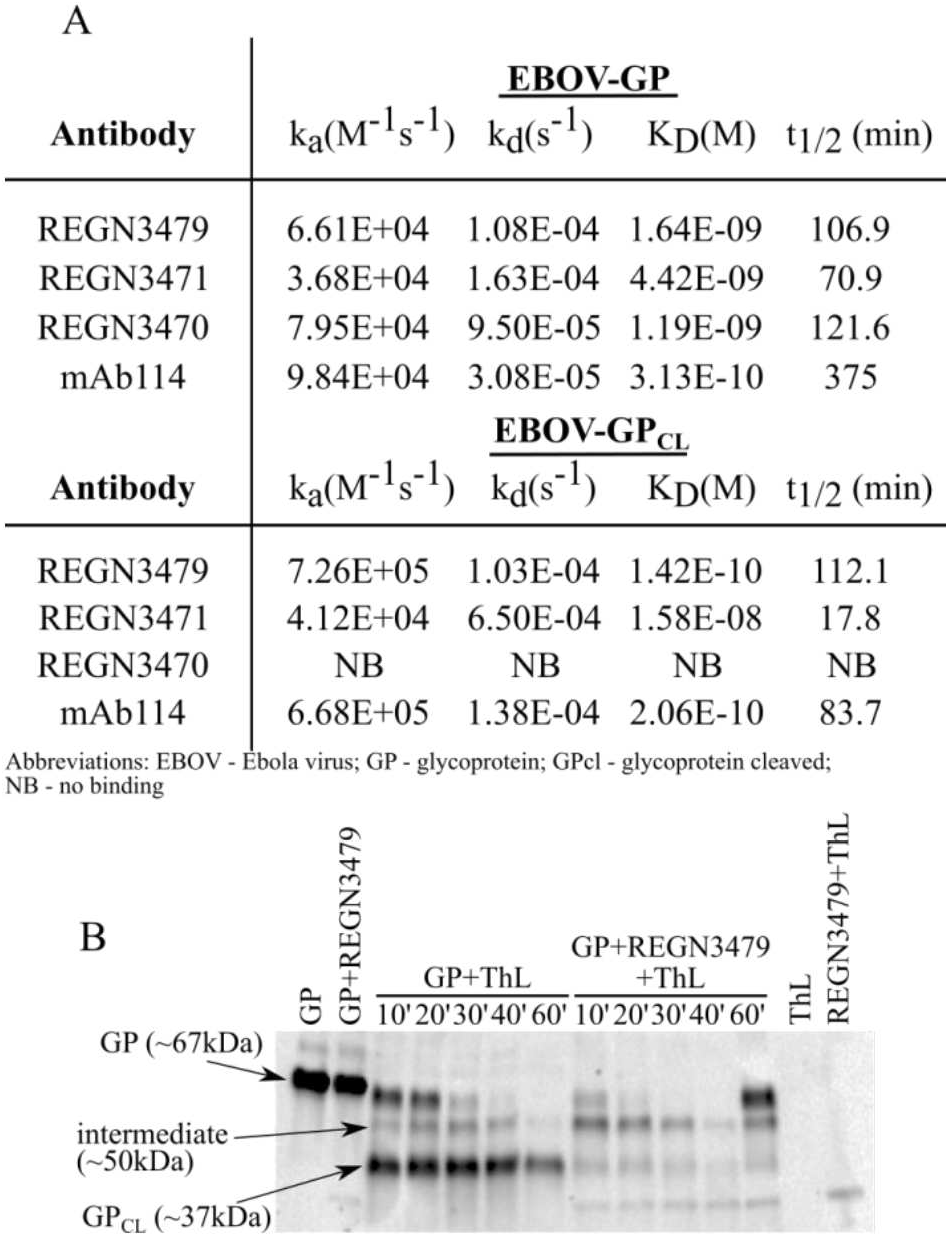
A) Ligand binding properties of REGN-EB3 cocktail antibodies. (A) Summary of equilibrium dissociation constants (Ko) for the interaction of surface-captured anti-EBOV antibody with recombinant EBOV GP or GPcL trimer protein respectively. ka, association rate constant; ktl, dissociation rate constant; Ko, equilibrium dissociation constant; b12, dissociative half-life. B) Time-dependent thermolysin cleavage of GP in the absence and presence of REGN3479.

We also treated pre-formed individual GP-Fab complexes with ThL and monitored the cleavage event over time using western blotting under non-reducing conditions. Digestion of unbound GP with ThL over time resulted in the disappearance of the major intact GP band (∼67kDa) and the appearance of the major digestion product, GPcL (−37kDa), as well as an intermediate (−50kDa) band. Interestingly, ThL digestion of REGN3479 complexed with GP showed a clear reduction in GP_CL_ band intensity and a small increase in intensity of the larger intermediate (∼50kDa). The marked reduction in GP_CL_ band intensity suggests that the presence of REGN3479 may block the ThL cleavage site (Figure 5B). No discernible difference in cleavage patterns was seen for GP complexed with either REGN3470 or REGN3471 compared to unbound GP (data not shown).

We further evaluated the formation of a stable GP_CL_-antibody complex using size-exclusion chromatography (SEC). GP_CL_ was incubated with each of the Fabs in the REGN-EB3 cocktail individually and the elution profile of the complex monitored using SEC (Figure S4). GP cleavage did not affect complex formation with REGN3479 as evidenced by the presence of two peaks in the SEC trace, one corresponding to the complex and the other to free Fab. REGN3470, for which the epitope lies exclusively in the glycan cap, does not bind cap-deleted GP, and thus showed two peaks corresponding to only GP_CL_ and free Fab, respectively. No peak for the complex was observed. REGN3471, for which the glycan cap comprises ∼40% of the epitope, exhibited some binding of GP_CL_, viewed as three peaks, corresponding to complex, GP_CL_ and free Fab, respectively (Figure S4).

### REGN-EB3 prevents rapid viral escape

We used a chimeric VSV virus expressing EBOV GP (VSV-EBOV-GP) to characterize the likelihood of escape mutations in GP following exposure to individual mAbs, and both competing and non-competing antibody combinations, targeting EBOV GP. As a first step, we tested the neutralization potency of these antibodies against VSV-EBOV-GP (Figure 6A and B). Consistent with previous reports, REGN3479 and mAb114 were potently neutralizing in our assay, with IC50s of 25.9 ng/ml and 54.7 ng/ml, respectively (Misasi et al., 2016). REGN3470 also demonstrated high neutralization potency, although neutralization never reached 100%, leaving a small population of infectious virus un-neutralized even at high antibody concentrations (Figure 6C), as previously observed (Pascal et al., 2018). REGN3471 was poorly neutralizing. The non competing REGN-EB3 and REGN3479+mAb114 combinations also potently neutralized VSV-EBOV-GP.

**Figure 6:**
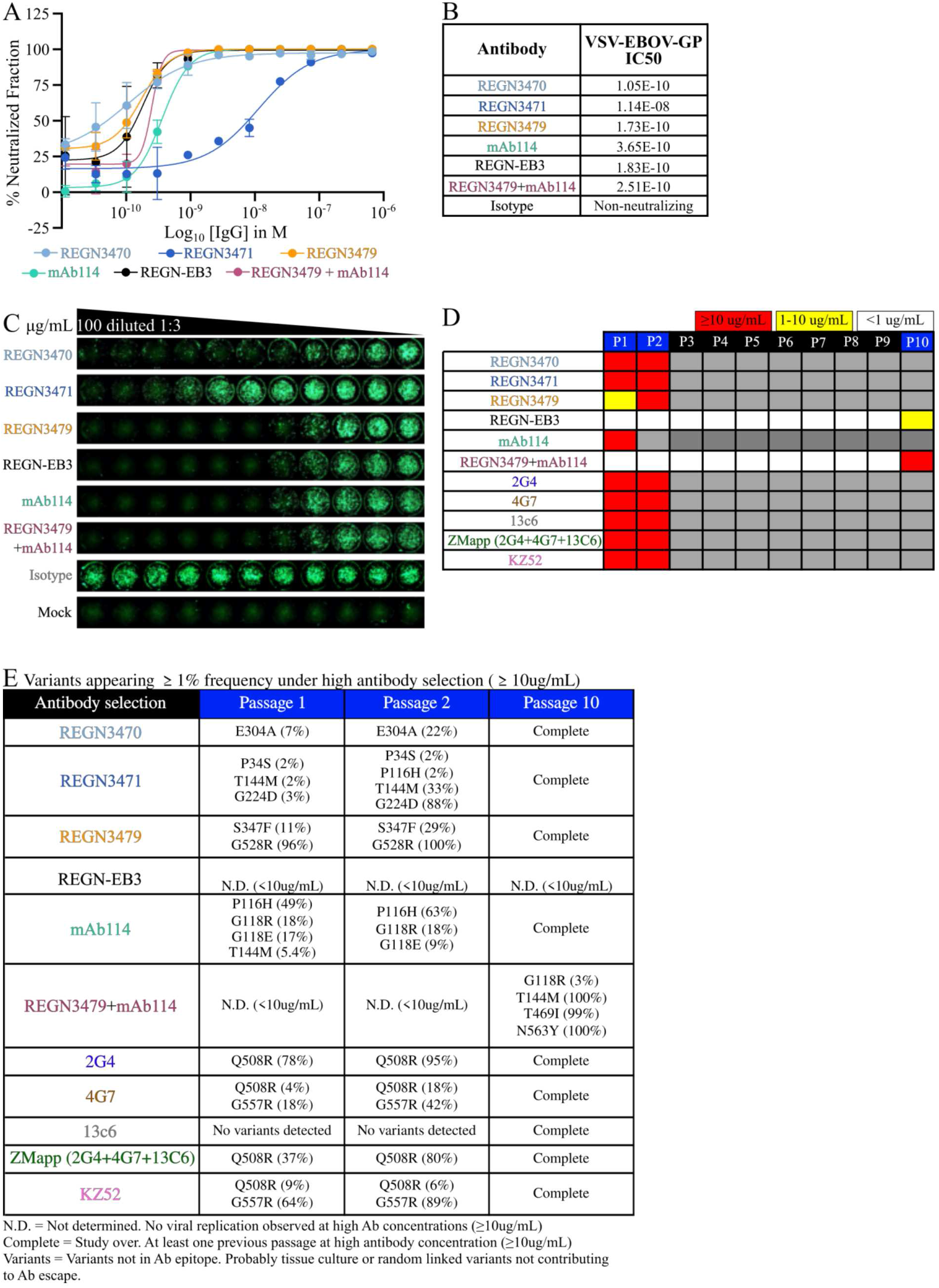
Neutralization of fully replicative VSV-EBOV-GP by the REGN-EB3 combination, individual REGN-EB3 components, mAb114, and a two mAb combination (REGN3479 + mAb114). A. Neutralization curves, B. IC50 values (in M), and C. Images of representative plates. D. Selection of escape variants to individual mAbs and combinations. To identify escape variants, VSV-EBOV-GP was propagated under antibody pressure to select for resistant virus. For each passage the maximal antibody concentration under which virus replication was observed is indicated: 2:H0ug1mL (red), H-10µg/mL (orange), <1ug/mL (white). E. Variants identified through RNAseq under greater than 2:H0µg1mL of antibody selection. Values in parentheses are the relative frequency of reads encoding a given variant among all reads in that position. N.D.: Not determined as no viral replication was observed at high (2:H0µg1mL) mAb concentration. Complete: at least one previous passage at high antibody concentration (2:H0ug1mL).

To determine the likelihood of mAb-induced viral escape and identify the mutations in GP associated with loss of therapeutic neutralizing activity, escape studies with VSV-EBOV-G were conducted by passaging virus in the presence of either individual mAbs or mAbs in combination. Virus resistance characterized by cytopathic cell death was observed within one or two passages for all mAbs used as a monotherapy. (Figure 6D). Sequencing analysis revealed that single mutations in the GP protein could result in complete viral resistance to all individual neutralizing mAbs (Figure 6E). Notably, escape mutations were selected within or adjacent to the contact residues of REGN3470, REGN3471, and REGN3479 (Figure 6E). Mutations that were previously shown to impact antibody neutralization were also selected for other antibodies against EBOV GP including KZ52 (G557R), 2G4 (G508R), and 4G7 (G557R) mAbs (Audet et al., 2014; Davidson et al., 2015; Mbala-Kingebeni et al., 2019; Miller et al., 2016; Qiu et al., 2012; Saphire et al., 2018b)

To understand how epitope conservation correlates with the development of antibody-induced resistance, we assessed the escape kinetics of mAb114 and REGN3479. mAb114 binds to a highly conserved region in the receptor-binding domain (RBD) required for viral entry (Misasi et al., 2016), while REGN3479 binds the conserved fusion loop and can neutralize multiple ebolaviruses, including EBOV Mayinga, EBOV Mali, Bundibugyo virus, and Sudan virus (Figure S5). Both mAb114 and REGN3479 rapidly selected resistant viruses within a single passage, similar to all other individual antibodies (Figure 6D and 6E). Furthermore, viruses resistant to mAb114 could be plaque purified and expanded without reversion of the escape mutations or obvious fitness defects (data not shown).

Next, we assessed the evolution of resistance to antibody cocktails containing components targeting overlapping (ZMapp) or distinct epitopes (REGN-EB3 and REGN3479+mAb114). Similar to the monotherapies, selection of a single variant (G508R) led to complete escape from the triple ZMapp combination containing two neutralizing antibodies that bind overlapping epitopes (2G4 and 4G7; both of which are affected by G508R) and one non-neutralizing antibody (13C6). G508R was also identified in an EBOV-infected cynomolgus macaque that succumbed to the virus after ZMapp therapy, confirming that similar mutations are selected in recombinant VSV-EBOV-GP virus in vitro as with authentic EBOV in vivo (Qiu et al., 2012). However, combinations of three (REGN-EB3) or two (REGN3479 + mAb114) antibodies binding distinct epitopes increased resistance to viral escape. Ten passages and multiple mutations were required to escape the potently neutralizing two mAb combination (REGN3479 + mAb114), and complete resistance was not observed with REGN-EB3 following 10 passages (Figure 6E).

To validate the mutants detected through sequencing analysis, we generated lentivirus-based pseudoparticles (EBOVpp) bearing individual escape mutations within the 2014 Zaire EBOV GP sequence. As expected, neutralization potency of every individual neutralizing mAb was impacted by mutations detected during selection (Table 1, Figure S6). Two identified mutants (P34S and T469I) did not impact antibody potency. These were likely tissue culture adaptations or were genetically linked to other variants that affect antibody neutralization selected during passaging. Despite the conservation of the mAb114 epitope, four mutants isolated under either mAb114 monotherapy or REGN3479+mAb114 cocktail selection were infectious and resulted in complete loss of mAb114 neutralization activity. A single mutation (G508R), also identified in previous studies (Qiu et al., 2012), was sufficient for complete resistance to ZMapp and the two competing individual neutralizing components (2G4 and 4G7). Importantly, no single individual escape mutation impacted neutralization of the non-competing three mAb (REGN-EB3) or two mAb (mAb114 + REGN3479) combinations that target distinct and non-overlapping antibody epitopes.

**Table 1:** Fold-decrease in potency relative to WT pseudovirus.

|  | REGN3479 | REGN-EB3 | mAb114 | REGN3479<br>+mAb114 | 2G4 | 4G7 | ZMapp | KZ52 |
| --- | --- | --- | --- | --- | --- | --- | --- | --- |
| WT | 1 | 1 | 1 | 1 | 1 | 1 | 1 | 1 |
| P34S | 0.6 | 0.4 | 0.4 | 0.5 | 0.7 | 0.6 | 0.6 | 0.5 |
| P116H | 0.6 | 0.6 | >76.3* | 0.8 | 0.7 | 0.7 | 0.9 | 0.6 |
| G118E | 0.5 | 0.4 | >29.1* | 0.5 | 0.9 | 0.7 | 0.7 | 0.5 |
| G118R | 0.6 | 0.5 | >33.6* | 0.6 | 0.9 | 0.7 | 0.8 | 0.6 |
| T144M | 1 | 1 | >33.6* | 1.1 | 1.1 | 1.1 | 1 | 0.9 |
| G224D | 0.6 | 0.6 | 1.3 | 0.8 | 0.7 | 0.7 | 0.7 | 0.7 |
| E304A | 0.7 | 0.7 | 0.5 | 0.6 | 0.3 | 0.2 | 0.4 | 0.4 |
| T469I | 0.8 | 0.7 | 0.6 | 0.7 | 0.9 | 0.7 | 0.6 | 0.7 |
| Q508R | 2.9 | 3.1 | 1.9 | 2.9 | >88.3* | >77.3* | >73.5* | >69.5* |
| G528R | >361.5* | 1.9 | 0.9 | 4 | 0.4 | 0.3 | 0.9 | 0.8 |
| G557R | 0.9 | 0.7 | 0.5 | 1 | 0.8 | >74.5* | 1.3 | >77.1* |
| N563Y | >424.0* | 0.9 | 0.4 | 3 | 0.5 | >173.4* | >162.1* | >151.7* |

\*Variant not neutralized at the antibody concentrations tested in this assay. The change in mAb potency for this variant was calculated by dividing the maximal antibody concentration tested by the IC50 value for the WT
pseudovirus. REGN3470, REGN3471, and 13C6 poorly neutralize lentivirus pseudovirus particles and could not be tested with escape variants.

## Discussion

The recent clinical success of mAb therapies for EBOV and SARS-CoV-2 clearly demonstrate that antibody therapeutics targeting viral proteins offer a valuable option for both treatment and prophylaxis of viral infections (Mulangu et al., 2019; O’Brien et al., 2021; Weinreich et al., 2021). Although vaccination will always be a critical component of the public health response to infectious agents, it is clear that a significant portion of the population will remain vulnerable to severe outcomes due to suboptimal responses to vaccines or challenges associated with vaccine access or uptake. Mass vaccination campaigns for infectious agents like EBOV may never occur, necessitating availability of therapeutics in the event of disease outbreaks that are unpredictable in timing and location. Antibody-based therapeutics may offer additional protection from severe disease and death even to vaccinated individuals that fall into high-risk categories. Thus, antibody therapies that can be rapidly deployed can provide immediate treatment and protect recipients until vaccination, and its associated immune response mounts. Indeed, 25% of PALM trial participants who experienced some level of EBOV disease, reported that they had been vaccinated before onset of symptoms (Mulangu et al., 2019).

Protection against EBOV infection by an antibody cocktail can be achieved by combining complementary activities, including both neutralizing and Fc-recruiting functions (Gunn et al., 2018; Saphire et al., 2018a). Furthermore, by limiting the likelihood of treatment emergent-drug resistance should be a critical component of any antiviral therapy. REGN-EB3 combines individual mAbs with neutralizing (REGN3479 and REGN3470), and Fc-recruiting (REGN3470 and REGN3471) functions. Importantly, this cocktail contains antibodies with non-overlapping footprints. The combination of distinct footprints simultaneously found on GP plays a key role in limiting the likelihood of treatment-induced drug resistance.

Here we determine the cryoEM structure at 3.1Å resolution of the complete REGN3470, REGN3471, REGN3479 cocktail bound to EBOV GP. Asymmetric reconstruction revealed eight Fab fragments simultaneously bound to the GP trimer: three each of REGN3479 and REGN3471, with two of REGN3470 in the final high-resolution model.

The ordering of this ∼600kDa complex, and binding footprint of the REGN3470 antibody contained within it, allows modeling for the first time of key portions of the glycan cap including the β17-β18 strands and the 27-residue loop in between. These sections were either fully disordered or only visible as main chain tubes of indiscernible N-to-C direction in previous crystal or cryo-EM structures. Here, they have been built in entirety, with visible side chains to confirm sequence and chain directionality. We are also not aware of any published EBOV glycoprotein structure bound to antibodies that have been resolved to this resolution or better using asymmetric reconstruction of cryoEM data.

The cryo-EM structure further revealed the contents and key contacts contained within the three non-overlapping footprints of the component members of the cocktail. REGN3479 binds to a quaternary epitope bridging the fusion loop tip and paddle of one GP monomer to both GP1 and GP2 of the neighboring monomer. One key mechanism of action is by locking the GP in the pre-fusion conformation, similar to other fusion loop binding antibodies like ADI-15878. The epitope recognized by REGN3479 almost completely overlaps with that of ADI-15878 (West et al., 2018). Indeed, 10 out of the 15 GP residues that interact with REGN3479 are shared with ADI-15878. As with ADI-15878, REGN3479 recognizes the conserved N-terminal pocket while displacing the non-conserved N terminus of GP2. However, only REGN3479 contacts GP2 HR1_A_. A second difference is that REGN 3479 binds only the glycan itself, while ADI-15878 also binds the Asn side chain to which the glycan is linked.

The general footprint and the overall vertical position of REGN3471 resemble those of mAb114 (Misasi et al., 2016), another potently neutralizing therapeutic antibody. A patch of hydrophobic residues in REGN3471 sits in the hydrophilic crest of the GP (Figure 3D, left). In contrast, in mAb114, two Arginines (Arg30 and Arg97) potentially form strong salt-bridging interactions with the GP crest residues Glu231 and Glu235, respectively (Figure 3D, right). Further, REGN3471 interacts with eight residues of the glycan cap while mAb114 interacts with only four. The greater dependence on glycan cap residues for binding likely explains why mAb114 exhibits higher affinity binding than REGN3471 to EBOV-GPcl, which likely leads to its greater potency in neutralization. The similarity of footprints and heavy chains utilized by these two antibodies demonstrates that Velocimmune mice can generate nearly identical antibody responses to humans (Macdonald et al., 2014; Murphy et al., 2014).

While physical neutralization of the antigen is a key process, studies have shown that immune effector functions play an equally important role (Saphire et al., 2018a). The immune effector functions are in turn controlled by several factors including antibody binding angle and accessibility of the Fc region to the respective Fc receptors. Each of the antibodies in the REGN-EB3 cocktail have identical Fc regions and yet REGN3470 and REGN3471 show potent immune effector functions while REGN3479 has none (Pascal et al., 2018). REGN3470 binds radially outward from the GP structure, anchoring solely to the glycan cap. REGN3471 binds the head, anchoring into the receptor-binding site of EBOV GP and bridging it to the glycan cap. Here, where soluble Fab are used, we observe three Fabs binding to the three copies of the receptor-binding site in the GP trimer. In vivo where complete IgG are used, complete occupancy may be more challenging. When homology models of REGN3470, REGN3471 and REGN3479 IgGs and a crystal structure of the receptor FcγR-IIIa bound to an IgG are docked into the cryoEM map, it is evident that the Fc regions of REGN3470 and REGN3471 are easily accessible for FcγR-IIIa receptor binding while the Fc region of REGN3479 has limited accessibility (Figure S7).

A key desired characteristic of any antiviral therapy is minimization of risk of drug-induced resistance that can impact the efficacy of the therapy in the treated individual as well as seed the mutated virus into the population. With antibody therapeutics these concerns may be especially great as mAb induced mutations may also impact vaccine efficacy since these mutations may occur in epitopes recognized by endogenous polyclonal antibody responses. We have previously assessed the generation of viral resistance to SARS-CoV-2 spike antibody monotherapies and combinations and demonstrated that resistant viruses were rapidly selected with monotherapies and antibody combinations targeting overlapping epitopes, regardless of epitope conservation (Baum et al., 2020; Copin et al., 2021). In contrast, antibody combinations binding distinct epitopes reduced the risk of rapid viral escape by requiring concurrent selection of resistance mutations to each antibody.

In this study, we analyzed the ability of antibodies targeting EBOV-GP to prevent viral resistance, both as monotherapies or in different cocktail combinations. As we observed with SARS-CoV-2, all antibodies used as monotherapy, even those antibodies (mAb114 and REGN3479) targeting conserved epitopes and predicted to be resistant to mutational escape (Gaudinski et al., 2019), as well as combinations binding overlapping epitopes, can lead to rapid selection of resistant viruses. In contrast, cocktails binding non-overlapping epitopes greatly reduced the likelihood of viral escape, whether composed of two (REGN379 + mAb114) or three (REGN-EB3) mAb components, by requiring multiple mutations blocking binding of each individual antibody for resistance. Importantly, in this study we assessed viral escape to neutralization by REGN-EB3, but additional constraints on antibody resistance in vivo may be imposed by the immune effector functions of REGN3470 and REGN3471.

The combined structural and functional analysis here reveals the footprints and key contacts of a therapeutic that treats Ebola virus disease. It also demonstrates that cocktails of antibodies targeting non-overlapping epitopes are key to mitigating the risk of drug-induced viral resistance.

## Materials and Methods

### GP Expression and purification

Both mucin-containing and mucin-deleted EBOV GP were produced from stably transfected *Drosophila melanogaster* S2 cells. Briefly, Effectene (Qiagen) was used to transfect S2 cells with a modified pMT-puro vector plasmid containing the GP gene of interest. Transfected cells were selected by incubation at 27°C for 4 weeks in complete Schneider’s medium with 6µg/mL puromycin. The cells were then transitioned to Insect Xpress medium (Lonza) for large-scale expression in 2-liter Erlenmeyer flasks. Expression of secreted GP ectodomain was induced with 500 mM CuSO_4_, and the supernatant was harvested after 5 days. The GP was engineered with a double Strep-tag at the C terminus to facilitate purification using a 5 mL Strep-trap HP column (GE) and then further purified by Superdex 200 size exclusion chromatography (SEC) in 50mM Tris-buffered saline (Tris-HCl, pH 7.5, 150mM NaCl [TBS]).

### Cleavage and purification of REGN3470, REGN3471 and REGN3479 Fab fragments

To produce Fab fragments, each antibody (10mg) was digested with 200µg papain (5% w/w) for four hours at 37 °C; the reaction was quenched with 50mM iodoacetamide. The digested protein was dialyzed overnight at 4 °C against 10mM Tris pH 8.0, 10mM NaCl buffer. The Fab fraction was then purified using a MonoQ 2mL column according to manufacturer’s protocol, followed by size exclusion chromatography using a Superdex 75 increase column equilibrated with 25mM Tris, 150mM NaCl, pH 7.5. The fractionated peak was concentrated in 10kDa MWCO Amicon Ultra 15 concentrators.

### REGN3471 crystal screening, data collection and structure determination

Crystal screening of unbound Fab fragments of REGN3471 (6.6mg/mL) was performed using sparse matrix screens and an Oryx crystallization robot (Douglas instruments). Initial hits for crystals were obtained with 20% isopropyl alcohol, 20% PEG 4000, 100mM sodium citrate, pH 5.6. The crystals diffracted to 2.3Å at APS beamline 23-ID-B. The collected data were processed using XDS. Crystals were indexed in space group P2_1_2_1_2_1_ and contained two Fabs in the asymmetric unit. Molecular replacement was used to iteratively determine phases. The initial molecular replacement model was generated by submitting the REGN3471 sequence to the SWISS-MODEL server [PMID: 29788355]. Rebuilding and refinement were performed using Phenix and Coot. Molprobity and OMIT maps were used to validate model quality.

### Preparation of GP-REGN-EB3 complex

To obtain complexes of GP with Inmazeb, GP (120µg), Fab fragments of REGN3470 (135µg), REGN3471 (125µg) and REGN3479 (150µg) were incubated together at room temperature overnight in a total volume of 500µL. The resulting complex was separated from excess Fabs using a Superose 6 Increase column equilibrated with 25mM Tris, 150mM NaCl, pH 7.5. Fractions containing the complex were combined and concentrated to 3.1mg/mL using a 100kDa MWCO Amicon concentrator.

### Cryo-EM Sample preparation and data collection

C-flat™ EM (CF-2/1-4C-T, Electron Microscopy Sciences) grids were glow-discharged for 15 seconds on a Pelco Easiglow at 15mW power. The purified GP-REGN-EB3 complex (3µL, ∼0.8mg/mL) was mixed with 1µ L 0.02mM lauryl maltose neopentyl glycol (LMNG) in 1X TBS and immediately applied to glow-discharged grids at 4 °C and 100% relative humidity inside a Vitrobot Mark IV. The grids were then blotted for 8 seconds without any extra applied blot force and plunge-frozen in liquid ethane. Frozen grids were imaged on a Thermo Fisher/Scientific Titan Krios (G3) equipped with a K2 direct electron detector and a BioQuantum energy filter (Gatan) using SerialEM software. In initial efforts to freeze the GP-Fab complex onto grids, particles stuck to the grid hole edges, making data acquisition exceedingly difficult. Mixing the complex with the amphiphilic detergent Lauryl Maltose Neopentyl glycol (LMNG), immediately before freezing dramatically improved the observed particle count and enabled successful data collection. Three separate datasets were collected on separate days. A total of 6,360 movies were collected at 1.0504Å/pixel with 30 frames per movie and a dose of 33-52e^-^/Å^2^. Defocus targets cycled from - 0.8 to -2.6 microns.

### Cryo-EM map calculation, structure determination and structure refinement

All cryo-EM data processing was performed using cryoSPARC. Patch Motion correction and Patch CTF estimation were used to align and calculate the contrast transfer function (CTF), respectively. The cryoSPARC blob picking utility [PMID: 28165473] was used to pick initial particles to develop a training data set of 3,154 particles that was fed into the Topaz particle picker (PMID: 29707703], which yielded 403,095 particles. The particle set was further 2D-classified to remove irrelevant particles. The resulting particle set was used for ab-initio reconstruction. Five ab initio classes were requested and all five classes were further subjected to heterogeneous refinement using all collected particles to generate representative 3D volumes Of the five 3D volumes, two, containing 121,135 and 155,580 particles each, showed more promising features (e.g., high-resolution features, antibody occupancy, trimeric organization) than the others. One of these two 3D volumes contains two copies of REGN3470 Fab, whereas the other contained three copies, one of which is poorly ordered. The two particle sets combined yielded a final higher resolution map that contained two copies of REGN3470 and three copies each of REGN3471 and REGN3479. Hence, although the structure shows two copies, it is clear that complete occupancy is possible in solution. These two classes were combined in homogenous refinement followed by the final, non-uniform 3D refinement. All refinements were done with no symmetry explicitly applied (C1). The final refinement produced a 3.1Å map (using the Fourier shell correlation [FSC]=0.143 criterion) containing 276,715 particles. This map was further improved using DeepEMhancer, a neural network-based density modification tool. Both the raw map and density-modified map were used to guide model building.

The GP-REGN-EB3 map contains density for three copies of GP in the trimer that are simultaneously bound to three copies of REGN3471, three copies of REGN3479 and two copies of REGN3470. The GP-REGN-EB3 model was built by initially docking copies of the unbound REGN3471 Fab crystal structure, homology models of REGN3470 and REGN3479 generated using SWISS-MODEL, and a published structure of GP (PDB 5JQ3). The EM density was clear for the Fv portions of the docked Fabs with unambiguous main-chain trace and conformation of all antibodies. Several rounds of Phenix [PMID: 31588918] refinement, followed by manual rebuilding using Coot [PMID: 20383002] yielded the final structure. Interacting residues of GP with the Fabs were selected using LigPlot+ [PMID: 21919503] and were further manually validated with interatomic distances <4Å for hydrogen bonds/salt bridges or hydrophobic surfaces with interatomic distances <5Å. Structures were visualized and analyzed using UCSF Chimera [PMID: 15264254] and UCSF ChimeraX [PMID: 32881101]. Publication figures were made using Inkscape [https://inkscape.org]

### BIAcore surface plasmon resonance analysis

Binding kinetics and affinities for each antibody in the REGN-EB3 cocktail and for mAb114 were individually assessed using surface plasmon resonance (SPR) technology on a Biacore T200 instrument (Cytiva, Marlborough, MA) using a Series S CM5 sensor chip in filtered and degassed HBS-EP running buffer (10 mM HEPES, 150 mM NaCl, 3mM EDTA, 0.05% (v/v) polysorbate 20, pH 7.4). Capture sensor surfaces were prepared by covalently immobilizing a mouse anti-human Fc mAb (REGN2567) on the chip surface using the standard amine coupling chemistry, as reported previously (Johnsson et al., 1991). Following surface activation, the remaining active carboxyl groups on the CM5 chip surface were blocked by injecting 1M ethanolamine, pH8.0 for 7 minutes. A typical resonance unit (RU) signal of ∼11,000 RU was achieved after the immobilization procedure.

Analysis of REGN3479, REGN3471, REGN3470 or mAb114 binding to EBOV-GP-ΔTM, and EBOV GP lacking the MLD (EBOV-GP-Δmuc) or glycan cap (glycan cap removed-EBOV-GP-Δmuc) was carried out by capturing the antibodies over immobilized anti-human Fc surface at 37ºC. Following the capture of the antibodies, different concentrations of EBOV-GP proteins (6.25nM-200nM, two-fold serial dilution in duplicate) in the running buffer were injected for 2.5 minutes at a flow rate of 50µL/min with an 8-minute dissociation phase. At the end of each cycle, the anti-human Fc surface was regenerated using a 12-second injection of 20mM phosphoric acid.

All specific SPR binding sensorgrams were double-reference subtracted as reported previously (Myszka, 1999) and the kinetic parameters were obtained by globally fitting the double-reference subtracted data to a 1:1 binding model with mass transport limitation using Biacore T200 Evaluation software v3.1 (Cytiva). The dissociation rate constant (*k*_*d*_) was determined by fitting the change in the binding response during the dissociation phase and the association rate constant (*k*_*a*_) was determined by globally fitting analyte binding at different concentrations. The equilibrium dissociation constant (K_D_) was calculated from the ratio of *k*_*d*_ to *k*_*a*_. The dissociative half-life (t_½_) in minutes was calculated as ln2/(*k*_*d*_*60).

### Thermolysin cleavage studies

GP (2µg) was mixed with 2.25µg of each antibody separately and incubated at RT overnight before -2µg of the complex was treated with 0.5µg Thermolysin L for 1 hour. Aliquots of the reaction were taken at indicated time points and quickly quenched with 5mM EDTA final concentration. Samples from each time point were evaluated by SDS-PAGE and western blotting.

### Size-exclusion chromatography

GPcl (50 µg) was used to make a complex with equal amounts of each of the antibodies in REGN-EB3 individually. Mixtures were incubated for 4hrs at room temperature and run on S200i size exclusion column (Cytiva) equilibrated with 25mM Tris-base, 150mM NaCl pH 7.5 to monitor complex formation. Uncleaved GP (GP) and GPcl alone were also separately run on the S200i column as controls.

### VSV-EBOV-GP generation

Fully replicative VSV-EBOV-GP was generated as previously described (Lawson, Rose/Whitt, Baum). In short, full-length North Kivu EBOV GP (GenBank accession MK007329.1) was cloned in place of VSV-G in a T7 promoter-driven VSV rescue plasmid. This genomic clone and the required expression plasmids for virus rescue (VSV-N, VSV-P, VSV-L, T7 polymerase) were transfected into HEK293T cells (ATCC CRL-3216) for 48 hours. The transfected cells were then co-cultured with BHK-21 cells (ATCC CCL-10) transfected with VSV-G using the SE cell Line 4D-Nucleofector X Kit L (Lonza) and cultured until virus-mediated CPE was observed. The recovered virus was then plaque-purified two twice, expanded, sucrose-cushioned to achieve 20-fold concentration, and frozen at -80 °C. Viral stocks were then titered by plaque assay using BHK-21 cells and sequence-confirmed by RNAseq.

### Escape studies

VSV-EBOV-GP studies were performed as described for VSV-SARS-CoV-2-spike with minor modifications PMID:34161776. Briefly, 1.25 × 10^6^ pfu of VSV-EBOV-GP was incubated with serial dilutions of antibodies at room temperature for 30 minutes, and then used to infect 2.5 × 10^5^ Vero cells (ATCC: CCL-81). The cells were then monitored for virus replication (CPE) for four days. Once the majority of cells displayed CPE, the supernatants were collected and frozen at -80 °C. Total RNA was extracted using TRIzol (Life Technologies) from the cell population and subjected to RNAseq to identify variants in the EBOV-GP protein. Each subsequent passage was performed by incubating 100µ L of the supernatant under the same or higher antibody concentration until resistance to 10µg/mL antibody selection was observed in at least one passage or up to 10 passages.

### VSV-EBOV-GP neutralization assays

Neutralization assays with the fully replicative VSV-EBOV-GP were performed by incubating 2,000 pfu of virus with serial dilutions of antibody starting at a concentration of 100µg/mL. The virus/antibody mixture was incubated at room temperature for 30 minutes and used to infect 2.0 × 10^4^ Vero cells at an MOI of 0.1. At 24 hours post-infection, cells were fixed with 2% paraformaldehyde and permeabilized with 0.1% Triton-X100. VSV-EBOV-GP infected cells were immunostained with a polyclonal rabbit anti-VSV serum (Imanis Life Sciences) that recognizes internal VSV proteins and an Alexa Fluor® 488 secondary antibody. Plate scans were imaged on a Cellular Technology Limited ImmunoSpot analyzer and fluorescent focus units (ffu) were determined using a SpectraMax i3 plate reader.

### Growth of Filoviruses

Filoviruses were grown on Vero E6 cells (ATCC, VA). Viral supernatants were harvested from tissue culture media and stored at -80°C before use. Titers were determined by TCID50 (EBOV, RESTV, BDBV, SUDV) or by FFU (MARV) assays in Vero E6 cells. For TCID50, plates were monitored for CPE and fixed 5 to 7 days post infection. TCID50 values were calculated using the Reed-Muench method (Reed, L.J., and Muench, H., 1938, A SIMPLE METHOD OF ESTIMATING FIFTY PER CENT ENDPOINTS. Am J Epidemiol 27, 493–497). For FFU assay, cells were infected for approximately 48 hours, fixed in 10% formalin, and prepared for antibody staining. The cells were permeabilized in 0.1% Triton X-100, washed, and blocked in 3.5% BSA. Infected cells were then stained with a polyclonal anti-MARV VLP antibody (IBT Bioservices 04-0005) and an AF488-conjugated goat anti-rabbit secondary antibody. Plates were imaged on a Cytation 1 Multimode Plate Reader.

### Filovirus neutralization assays

Antibodies were diluted into culture medium containing 10% fetal bovine serum and incubated with virus for 1 hour at 37° C. Antibody concentrations ranged from 20 µg/ml to 0.5 pg/ml. The mixture was added to Vero E6 cells for 48 hours. Following incubation, cells were fixed in 10% neutral buffered formalin and virus was detected using smiFISH as previously described (PMID: 27599845). Briefly, cells were permeabilized with 70% ethanol and fixed with 15% formamide. Infected cells were detected with oligonucleotides targeting viral mRNAs from each filovirus and a fluorescently labeled (Cy5) detection probe (Integrated DNA Technologies). Images were acquired on a Nikon Ti2 using a 10X lens in DAPI and Cy5 channels. Images were processed using a CellProfiler pipeline to quantify nuclear staining intensity and mRNA intensity for each image. Infection was quantified as viral mRNA intensity normalized to nuclear intensity for each well, and this was further normalized to the average of infection values from wells treated with the lowest dilution of antibody.

### Pseudoparticle generation

EBOV GP pseudoparticles (EBOVpp) were generated as described [PMID: 29860496]. HEK293T cells were transfected with expression plasmids carrying the 2014 Zaire EBOV GP (KJ660346), lentivirus capsid proteins, and a lentivirus genome encoding a firefly luciferase under the control of the retrovirus long terminal repeat promoter. Supernatants were collected 48 hours post-transfection, clarified by centrifugation, and stored at -80 °C. Individual variants were cloned into 2014 Zaire EBOV GP (KJ660346) using site-directed mutagenesis and pseudotyped as described above.

### Pseudoparticle neutralization assays

Pseudoparticle neutralization assays were performed by incubating EBOV pseudoparticles with serial dilutions of the indicated antibody. After 1 hour, the VLP/antibody mixtures were added to resuspended Huh7 (JCRB Cell Bank) and incubated for 72 hours. Luciferase signal from the lentivirus reporter was detected using a BrightGlo luciferase assay kit (Promega) and a Victor X3 plate reader (PerkinElmer).

## Supporting information

PDB validation report

## Acknowledgements

We gratefully acknowledge our funding from NIAID U19 AI142790, Consortium for Immunotherapeutics against Emerging Viral Threats (E.O.S, C.W.D.) and U.S. Department of Health and Health ServicesContract No. HHSO100201700016C (Regeneron). A portion of this project has been funded in part with federal funds from the Department of Health and Human Services, Office of the Assistant Secretary for Preparedness and Response, Biomedical Advanced Research and Development Authority, under OT number HHSO100201700020C We thank all the staff of the NIH who supported this study, in particular Kaleb Sharer, Russel Byrum, Jennifer Jackson, Sarah Klim, Danny Ragland, Marisa St Claire and Lisa Hensley. The authors would like to thank Kristen Tramaglini for continuous support with this project.

## Author Contributions

**Conceptualization**, E.O.S, C.A.K, A.B.

**Methodology**, V.R, B.F, A.R, H.C, A.P, K.P, R.C, R.D

**Investigation**, V.R, H.C, C.H, B.F, A.B, A.R, E.A, K.S, S.H, D.P, E.W, S.G, R.C, K.P, R.D, R.M.B, C.D

**Formal analysis**, V.R, C.H, B.F, A.B, A.R, M.F, R.D, K.P

**Writing - original draft**, V.R, E.O.S., B.F., A.B.

**Writing - review & editing**, V.R, E.O.S, B.F, M.F, S.S, A.P, C.A.K, A.B

**Visualization**, V.R, B.F., A.R., H.C.

**Supervision**, E.O.S, C.A.K, M.F., A.B.

**Resources**, E.O.S, D.W, C.A.K, A.B, A.R,.

**Funding Acquisition**, E.O.S, C.A.K.

## Data Availability

EMDB PDB: (will provide later)

