## Supplementary information for "Structure of the Inmazeb cocktail and resistance to escape against Ebola virus"

### Supplementary material

Figure S1: Crystal structure of REGN3471. Heavy chain shown in indigo, light chain shown in blue. Complementarity determining regions (CDRs) are labeled.

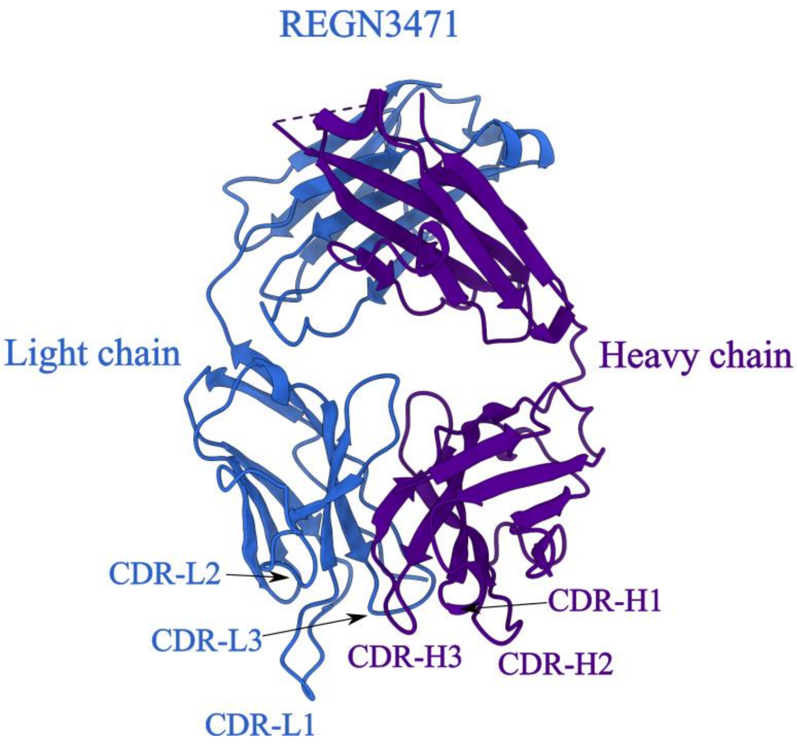

Figure S2: CryoEM single particle analysis workflow using cryoSPARC. Raw images, representative 2D classes, ab-initio and hetero refined models, FSC and final map are shown.

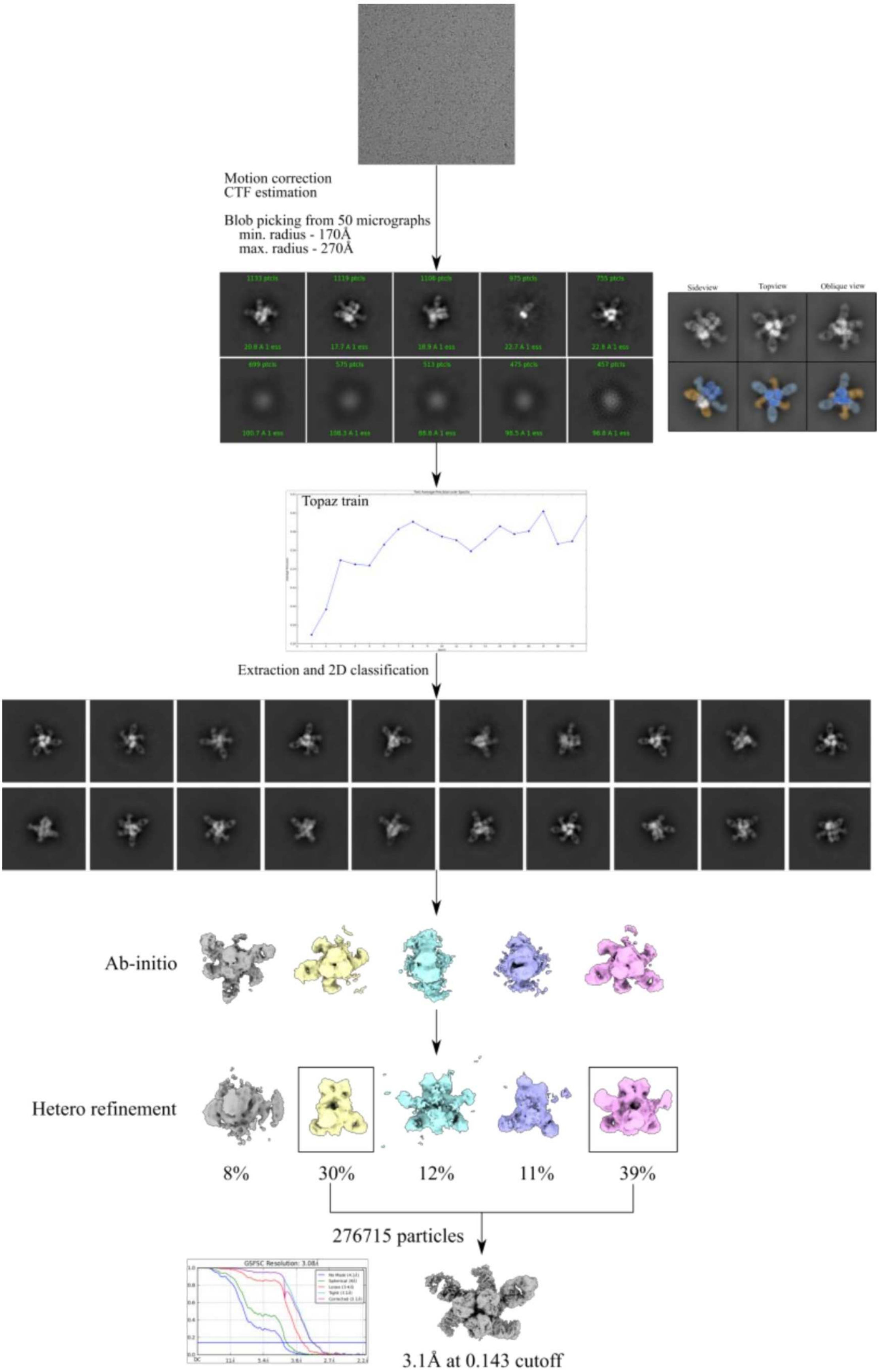


Table S1: Cryo-EM data collection parameters

| | |
|---|---|
| Protein | zGP-REGN-EB3 complex |
| EMDB | |
| Microscope | ThermoFisher FEI Titan Krios |
| Voltage (kV) | 300 |
| Magnification (nominal) | 22,500x |
| Pixel size (Å/pix) | 1.0504 |
| Frames per exposure | 30-40 |
| Defocus range (µm) | -0.8 to -2.6 |
| Micrographs collected | 6360 |
| Particles extracted/final | 546243/276715 |
| Symmetry imposed | n/a (C1) |
| Unmasked resolution at 0.143 FSC (Å) | 4.1 |
| Masked resolution at 0.143 FSC (Å) | 3.1 |

Table S2: Model refinement statistics from Molprobity

PDB ID

===============================================================

Composition (#)

| | |
|---|---|
| Chains | 28 |
| Atoms | 22621 (Hydrogens: 0) |
| Residues | Protein: 2874 Nucleotide: 0 |
| Water | 0 |
| Ligands | BMA: 3 |
| | NAG: 14 |
| | MAN: 9 |
| Bonds (RMSD) | |
| Length (Å) (# > 4σ) | 0.006 (0) |
| Angles (°) (# > 4σ) | 0.815 (10) |
| MolProbity score | 1.89 |
| Clash score | 17.84 |
| Ramachandran plot (%) | |
| Outliers | 0.07 |
| Allowed | 2.66 |
| Favored | 97.27 |
| Rama-Z (Ramachandran plot Z-score, RMSD) | |
| whole (N = 2817) | 0.46 (0.15) |
| helix (N = 190) | 1.44 (0.37) |
| sheet (N = 975) | 0.90 (0.16) |
| loop (N = 1652) | 0.16 (0.15) |
| Rotamer outliers (%) | 0.00 |
| Cβ outliers (%) | 0.00 |
| Peptide plane (%) | |
| Cis proline/general | 0.0/0.0 |
| Twisted proline/general | 0.0/0.0 |
| CaBLAM outliers (%) | 1.56 |
| ADP (B-factors) | |
| Iso/Aniso (#) | 22621/0 |
| min/max/mean | |
| Protein | 63.64/222.47/119.59 |
| Nucleotide | --- |
| Ligand | 92.95/154.11/120.36 |
| Water | --- |
| Occupancy | |
| Mean | 1.00 |
| occ = 1 (%) | 100.00 |
| 0 < occ < 1 (%) | 0.00 |
| occ > 1 (%) | 0.00 |

Data
==========================================================================

| | | |
|---|---|---|
| Box | | |
| Lengths (Å) | 138.65, 155.46, 143.90 | |
| Angles (°) | 90.00, 90.00, 90.00 | |
| Supplied Resolution (Å) | 3.1 | |
| Resolution Estimates (Å) | Masked | Unmasked |
| d FSC (half maps; 0.143) | --- | --- |
| d 99 (full/half1/half2) | 3.9/---/--- | 3.8/---/--- |
| d model | 3.5 | 3.5 |
| d FSC model (0/0.143/0.5) | 3.0/3.1/3.3 | 3.0/3.1/3.5 |
| Map min/max/mean | -0.62/1.72/-0.00 | |

Model vs. Data
==========================================================================

| | |
|---|---|
| CC (mask) | 0.82 |
| CC (box) | 0.77 |

| | |
|---|---|
| CC (peaks) | 0.58 |
| CC (volume) | 0.82 |
| Mean CC for ligands | 0.82 |

Figure S3: SPR-Sensograms. Ligand binding properties of anti-Ebola antibodies. (A) Summary of equilibrium dissociation constants ($K_D$) for the interaction of surface-captured anti-Ebola antibody with recombinant EBOV GP trimer protein respectively. $k_a$, association rate constant; $k_d$, dissociation rate constant; $K_D$, equilibrium dissociation constant; $t_{1/2}$, dissociative half-life. The association phase of

EBOV GP trimer proteins was monitored at 50μL/min for  minutes over anti-Ebola antibody captured surfaces. Sensogram of binding kinetics for each antibody in the REGN-EB3 cocktail and mAb114 to (A) EBOV GP Δmuc (uncleaved), (B) EBOV GP ΔTM, and (C) EBOV GPcl Δmuc (cleaved). EBOV GP proteins was tested in duplicate in a 2-fold dilution series concentration ranging from 6.25nM- 200nM, are shown as black lines. The data were globally fit to a 1:1 binding interaction model using T200 evaluation software 3.1. Kinetic fits from the analyses are overlaid on the binding data in red.

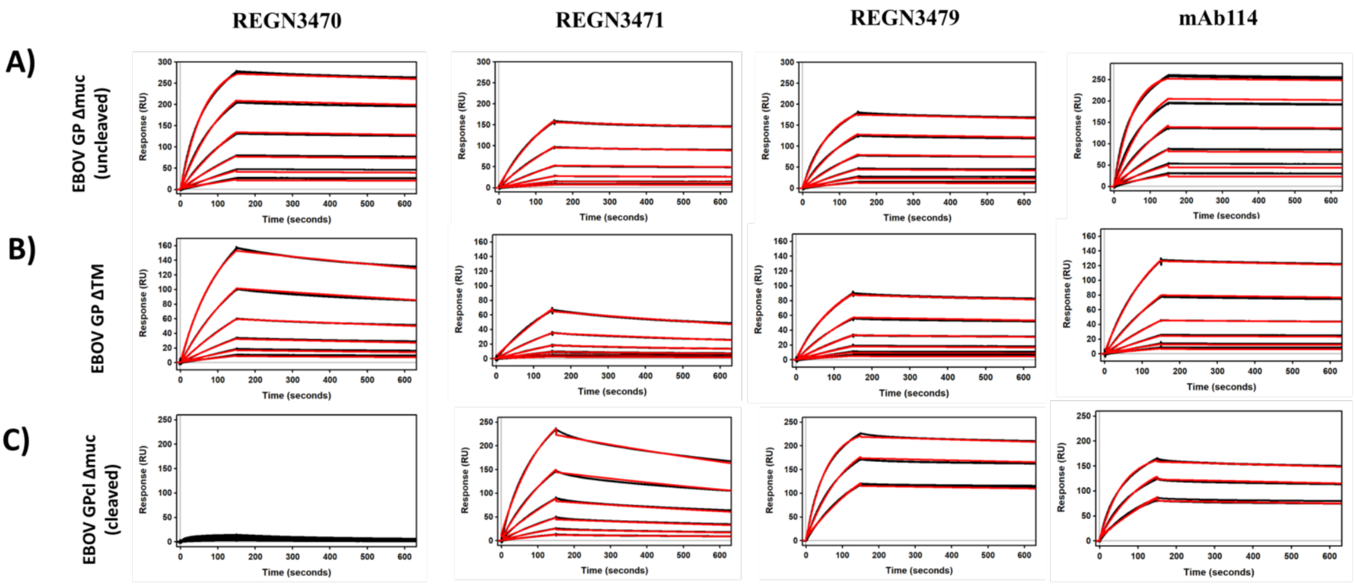


Figure S4: Size exclusion traces of unbound GP, GPcl and GPcl mixed with REGN3470, REGN3471 and REGN3479. Complex, GPcl, and Fab alone are marked with grey bounding boxes. Complex formation is seen clearly with REGN3479. REGN3471 shows a small population of a complex but majority exists as free Fab. No complex formation is seen with REGN3470.

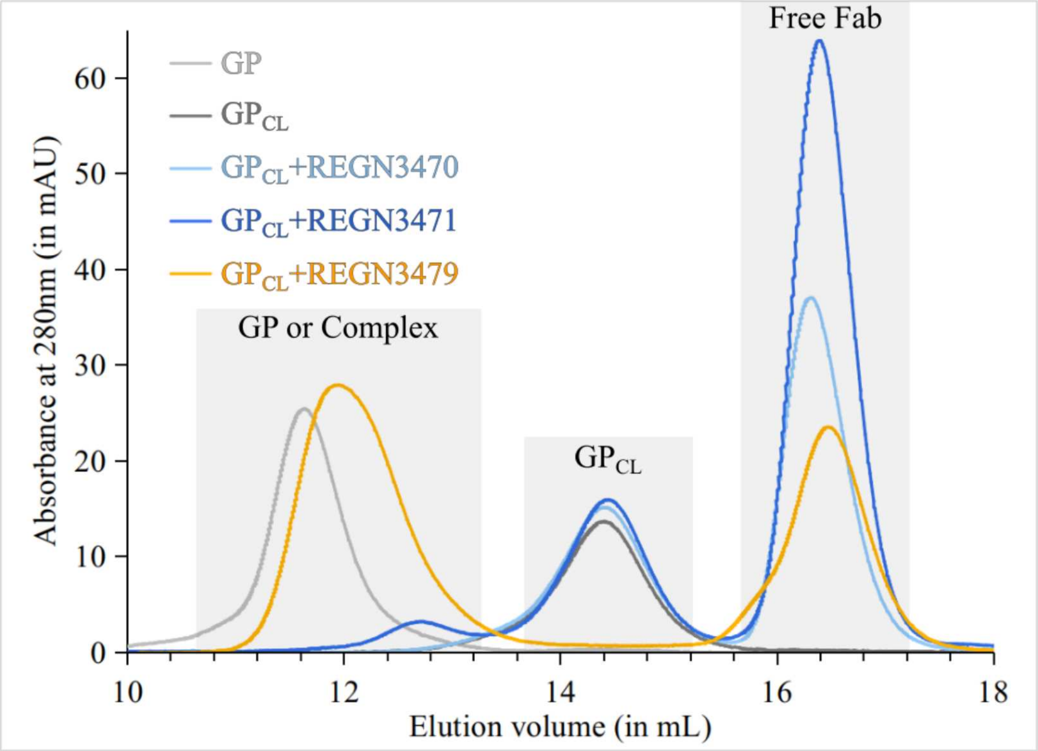


**Table S3: List of GP residues that interact with each of the antibodies in the REGN-EB3 cocktail**

| Antibody | GP Residue Total = 15 | Antibody Residue(s) Interacting with Indicated GP Residue | |
|---|---|---|---|
| | | Heavy Chain Total = 12 | Light Chain Total = 7 |
| REGN3470 | Thr259 | - | Tyr32 |
| | Ser263 | - | Ser30; Tyr32; Phe 92 |
| | Lys276 | Trp52 | - |
| | Val277 | Trp52 | - |
| | Asn278 | Asn31; Trp52; His53 | - |
| | Glu280 | Asn31; Tyr32; Gly33; Asn99; Trp100 | Thr94 |
| | Ile281 | Asn 31; Tyr32 | - |
| | Asp282 | Phe 27; Thr 28; Asn31; Tyr32 | - |
| | Arg302 | - | Tyr49 |
| | Ser303 | - | Tyr32 |
| | Glu304 | Asn101 | Thr 31; Tyr32; Ala50 |
| | Leu306 | Asn101 | - |
| | Ser307 | Trp52; Trp 100; Asn 101 | - |
| | Phe308 | - | Thr94 |
| | Thr309 | Trp 52; Asp57; Tyr59 | - |
| Antibody | GP Residue Total = 21 | Antibody Residue(s) Interacting with Indicated GP Residue | |
| | | Heavy Chain Total = 15 | Light Chain Total = 10 |
| REGN3471 | Glu112 | - | Tyr31; Ser33 |
| | Lys 114 | - | Tyr98 |
| | Lys115 | Tyr58 | - |
| | Pro116 | Asp33; His35; Asp56; Tyr58; Leu103 | - |
| | Asp117 | Trp47; Tyr58; Leu103 | Ser100; Leu102 |
| | Gly118 | Leu103 | Ser99; Ser100 |
| | Ser119 | - | Ser100 |
| | Gly143 | Gly101 | - |
| | Thr144 | Asp33; Gly101; Leu103 | - |
| | Pro146 | Ala54; Ala56 | - |
| | Arg172 | - | Tyr31 |

| | | | |
|---|---|---|---|
| | Gln221 | Trp99; Phe100; Glu102 | - |
| | Thr223 | Ser31; Thr53; Phe100; Gly101 | - |
| | Glu231 | Ser31; Phe100 | - |
| | Leu233 | Phe100 | - |
| | Leu239 | - | Ser62 |
| | Tyr241 | Trp99 | - |
| | Thr269 | - | Ser62 |
| | Gly271 | - | Ser62 |
| | Lys272 | Trp99; Tyr104 | Tyr55; Glu61 |
| | Ile274 | Tyr32; Trp99 | - |
| | (linked to Asn238) | - | Thr59 |
| **Antibody** | **GP Residue Total = 15** | **Antibody Residue(s) Interacting with Indicated GP Residue** | |
| | | **Heavy Chain Total = 21** | **Light Chain Total = 9** |
| REGN3479 | Pro34 | Ser57 | - |
| | Leu43 | Met54 | - |
| | Val45 | Met54; Gly55; Gly56 | - |
| | Ile504 | Thr28; Ser30; Ser31 | - |
| | Val505 | Ser31 | - |
| | Ala507 | Ser31; Tyr32 | - |
| | Ile527 | Trp47; Thr50; Tyr59 | Thr94; Leu95 |
| | Gly528 | - | Ser91; Tyr92; Ser93; Leu95 |
| | Leu529 | Tyr101; Pro102; His103 | Phe32; Ser91; Tyr92 |
| | Ala530 | - | Tyr92 |
| | Phe535 | - | Phe32; Tyr92 |
| | Gly536 | - | Tyr92 |
| | Gln560 | Met54 | - |
| | Glu564 | Ser52; Gly53 | - |
| | Gln567 | Tyr59; Arg99 | - |
| | NAG633 (linked to Asn563) | Tyr101 | - |

| | | | |
|---|---|---|---|
| | **BMA634 (linked to Asn 563)** | **Tyr101** | **Tyr49** |
| | **MAN637 (linked to Asn563)** | **Lys98; Arg99; Gly100; Tyr101, Ser104, Asp106** | **Leu46; Tyr49; Gln55** |

Figure S5: Neutralization of filoviruses by REGN3479. Infection was detected by smiFISH and normalized to the average of infection values from wells treated with the lowest dilution of antibody. Abbreviations: Ebola virus - EBOV, Bundibugyo virus – BUNDV, Sudan virus -SUDV, Marburg virus -MARV, Reston virus – RESTV.

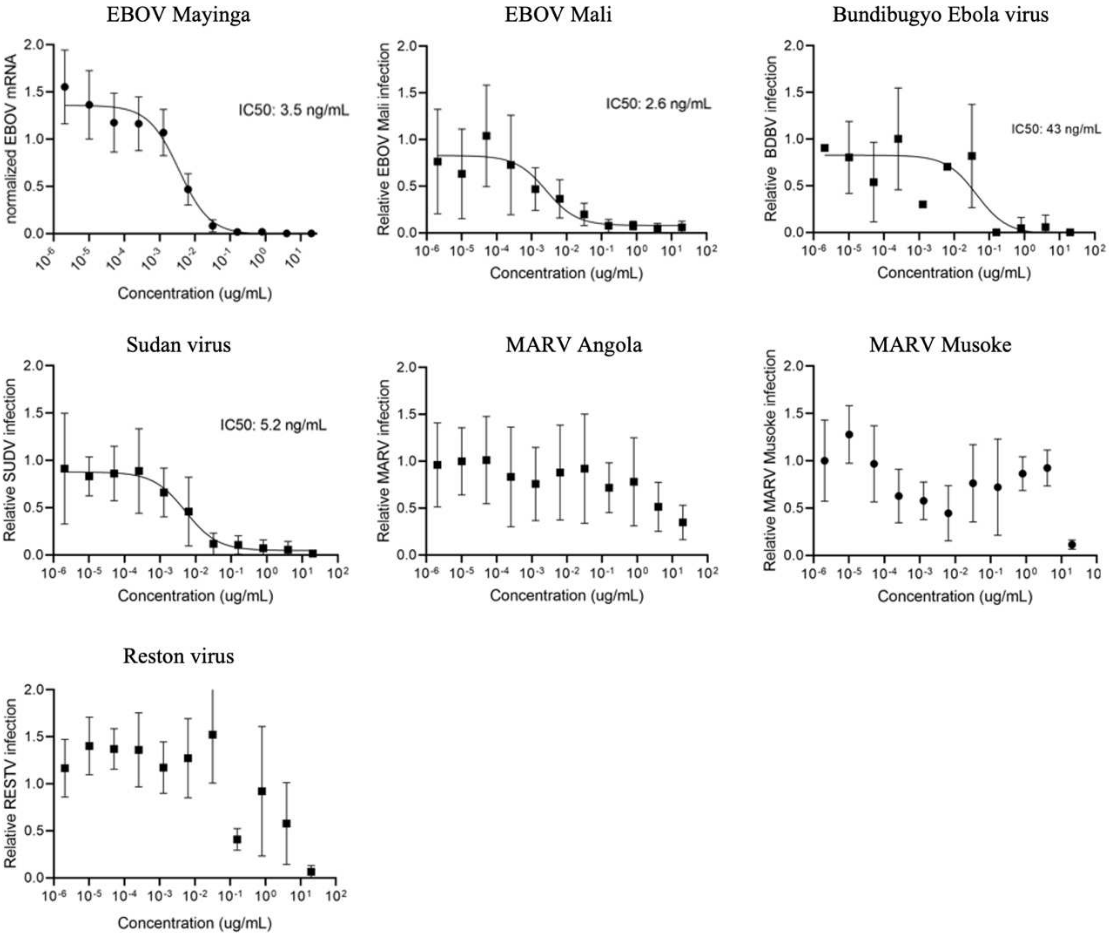
EBOV Mayinga
normalized EBOV mRNA
IC50: 3.5 ng/mL
Concentration (ug/mL)
EBOV Mali
Relative EBOV Mali infection
IC50: 2.6 ng/mL
Concentration (ug/mL)
Bundibugyo Ebola virus
Relative BDBV infection
IC50: 43 ng/mL
Concentration (ug/mL)
Sudan virus
Relative SUDV infection
IC50: 5.2 ng/mL
Concentration (ug/mL)
MARV Angola
Relative MARV infection
Concentration (ug/mL)
MARV Musoke
Relative MARV Musoke infection
Concentration (ug/mL)
Reston virus
Relative RESTV infection
Concentration (ug/mL)

Figure S6: Neutralization of lentivirus virus particles pseudotyped with EBOV GP variants detected after antibody selection.

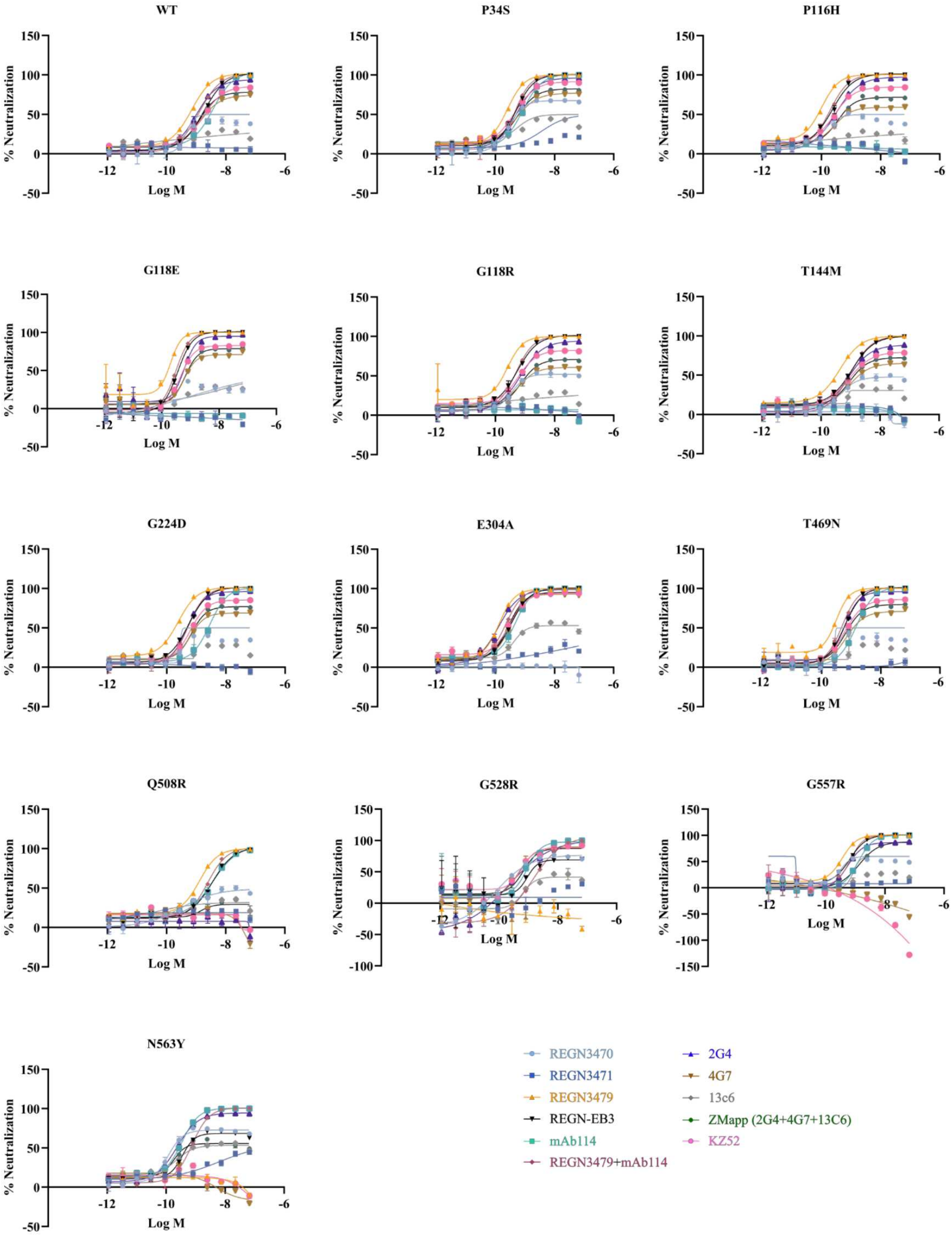

Figure S7: Homology models of all REGN-EB3 IgGs were generated using SWISS-MODEL and docked into our cryoEM structure. Structure of FcγRIIIa receptor bound to a Fab was obtained using PDB ID: 1T83 and was docked onto each of the IgG. It is evident from this representation that the binding of the FcγRIIIa receptor is feasible only with REGN3470 and REGN3471 and not with REGN3479.

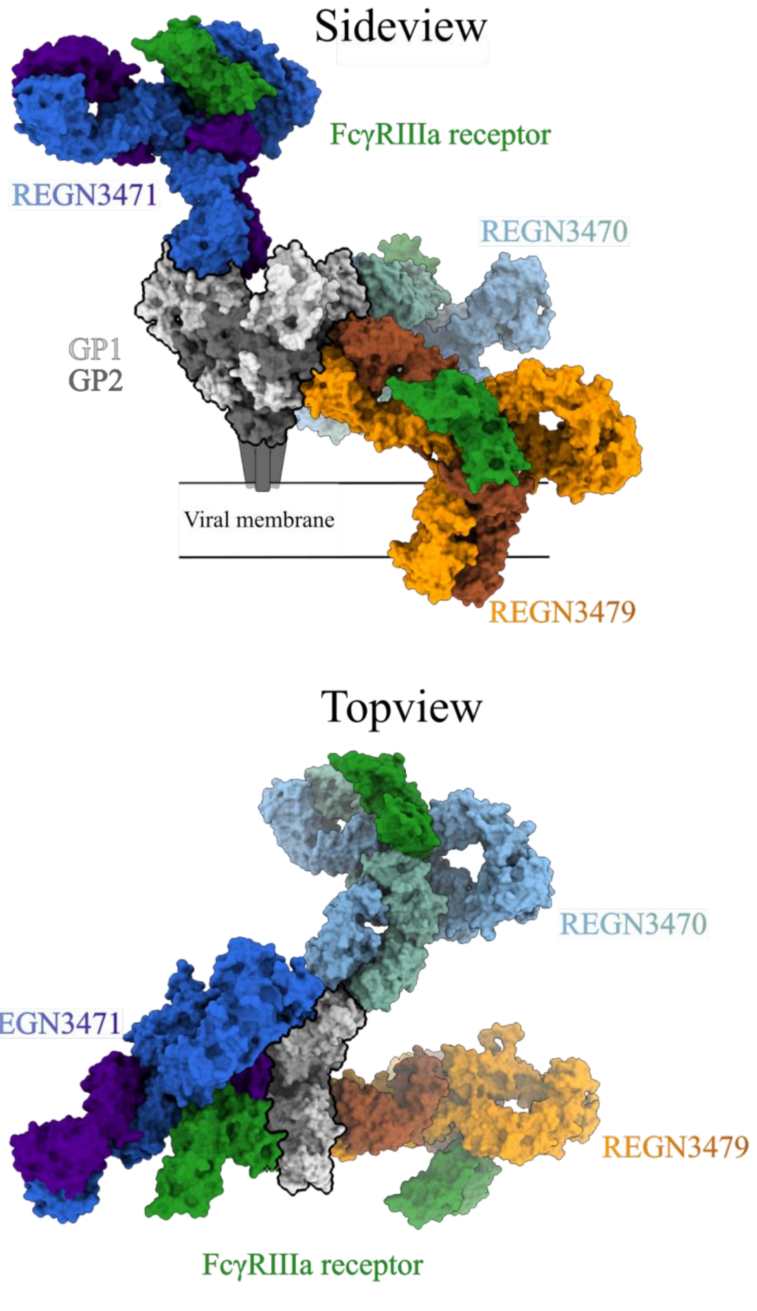
