## Supplementary material for "Structure of the Inmazeb cocktail and resistance to escape against Ebola virus": PDB validation report

Click here to access/download;PDB Validation Report;Inmazeb_Full_validation.pdf

### Full wwPDB EM Validation Report (i)

Jan 24, 2022 – 03:32 PM EST

| | | |
|---|---|---|
| PDB ID | : | 7TN9 |
| EMDB ID | : | EMD-26005 |
| Title | : | Structure of the Inmazeb cocktail and resistance to escape against Ebola virus |
| Deposited on | : | 2022-01-20 |
| Resolution | : | 3.10 Å(reported) |
| Based on initial model | : | 5JQ3 |

**This wwPDB validation report is for manuscript review**

For Manuscript Review

This is a Full wwPDB EM Validation Report.

This report is produced by the wwPDB biocuration pipeline after annotation of the structure.

We welcome your comments at **
A user guide is available at
https://www.wwpdb.org/validation/2017/EMValidationReportHelp
with specific help available everywhere you see the (i) symbol.

The following versions of software and data (see references (i)) were used in the production of this report:

| | | |
|---|---|---|
| EMDB validation analysis | : | 0.0.0.dev97 |
| Mogul | : | 1.8.5 (274361), CSD as541be (2020) |
| MolProbity | : | 4.02b-467 |
| buster-report | : | 1.1.7 (2018) |
| Percentile statistics | : | 20191225.v01 (using entries in the PDB archive December 25th 2019) |
| Ideal geometry (proteins) | : | Engh & Huber (2001) |
| Ideal geometry (DNA, RNA) | : | Parkinson et al. (1996) |
| Validation Pipeline (wwPDB-VP) | : | 2.26 |

### 1 Overall quality at a glance (i)

The following experimental techniques were used to determine the structure:
*ELECTRON MICROSCOPY*

The reported resolution of this entry is 3.10 Å.

Percentile scores (ranging between 0-100) for global validation metrics of the entry are shown in the following graphic. The table shows the number of entries on which the scores are based.

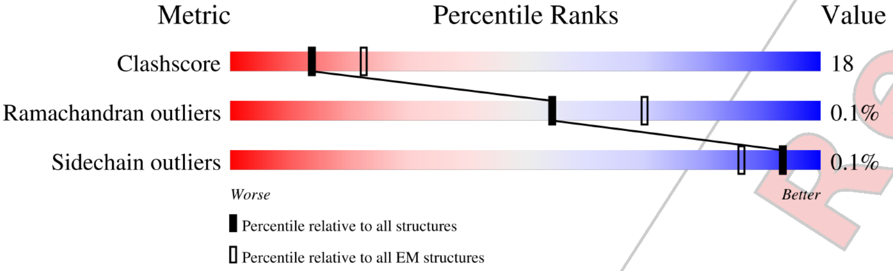


| Metric | Whole archive (#Entries) | EM structures (#Entries) |
|---|---|---|
| Clashscore | 158937 | 4297 |
| Ramachandran outliers | 154571 | 4023 |
| Sidechain outliers | 154315 | 3826 |

The table below summarises the geometric issues observed across the polymeric chains and their fit to the map. The red, orange, yellow and green segments of the bar indicate the fraction of residues that contain outliers for >=3, 2, 1 and 0 types of geometric quality criteria respectively. A grey segment represents the fraction of residues that are not modelled. The numeric value for each fraction is indicated below the corresponding segment, with a dot representing fractions <=5% The upper red bar (where present) indicates the fraction of residues that have poor fit to the EM map (all-atom inclusion < 40%). The numeric value is given above the bar.

| Mol | Chain | Length | Quality of chain |
|---|---|---|---|
| 1 | A | 217 | 33% 20% 47% |
| 1 | C | 217 | 34% 20% 47% |
| 1 | E | 217 | 38% 15% 47% |
| 2 | S | 282 | • 58% 26% 16% |
| 2 | T | 282 | 65% 30% • 5% |
| 2 | U | 282 | • 63% 32% 5% |
| 3 | N | 213 | • 28% 21% 52% |
| 3 | P | 213 | • 31% 16% 53% |

*Continued on next page...*

For Manuscript Review

*Continued from previous page...*

| Mol | Chain | Length | Quality of chain |
|---|---|---|---|
| 3 | R | 213 | 37% 12% 51% |
| 4 | B | 220 | 39% 12% 49% |
| 4 | D | 220 | 37% 14% 49% |
| 4 | F | 220 | 31% 20% 49% |
| 5 | V | 175 | 43% 19% 37% |
| 5 | W | 175 | 37% 26% 37% |
| 5 | X | 175 | 40% 21% 39% |
| 6 | M | 217 | 28% 25% 47% |
| 6 | O | 217 | 32% 21% 47% |
| 6 | Q | 217 | 33% 21% 47% |
| 7 | G | 215 | 31% 22% 47% |
| 7 | I | 215 | 16% / 28% 24% 47% |
| 8 | H | 215 | 26% 24% 50% |
| 8 | J | 215 | 25% / 21% 22% 56% |
| 9 | K | 6 | 67% 33% |
| 9 | L | 6 | 83% 17% |
| 9 | Y | 6 | 83% 17% |

For Manuscript Review

#### 2 Entry composition (i)

There are 10 unique types of molecules in this entry. The entry contains 22621 atoms, of which 0 are hydrogens and 0 are deuteriums.

In the tables below, the AltConf column contains the number of residues with at least one atom in alternate conformation and the Trace column contains the number of residues modelled with at most 2 atoms.

- Molecule 1 is a protein called REGN3471 heavy chain.

| Mol | Chain | Residues | Atoms | | | | | AltConf | Trace |
|---|---|---|---|---|---|---|---|---|---|
| 1 | A | 116 | Total 895 | C 570 | N 151 | O 170 | S 4 | 0 | 0 |
| 1 | C | 116 | Total 895 | C 570 | N 151 | O 170 | S 4 | 0 | 0 |
| 1 | E | 116 | Total 895 | C 570 | N 151 | O 170 | S 4 | 0 | 0 |

- Molecule 2 is a protein called Envelope glycoprotein.

| Mol | Chain | Residues | Atoms | | | | | AltConf | Trace |
|---|---|---|---|---|---|---|---|---|---|
| 2 | T | 269 | Total 2127 | C 1359 | N 361 | O 402 | S 5 | 0 | 0 |
| 2 | S | 237 | Total 1872 | C 1205 | N 313 | O 349 | S 5 | 0 | 0 |
| 2 | U | 269 | Total 2127 | C 1359 | N 361 | O 402 | S 5 | 0 | 0 |

- Molecule 3 is a protein called REGN3479 light chain.

| Mol | Chain | Residues | Atoms | | | | | AltConf | Trace |
|---|---|---|---|---|---|---|---|---|---|
| 3 | R | 105 | Total 796 | C 498 | N 132 | O 163 | S 3 | 0 | 0 |
| 3 | N | 103 | Total 779 | C 487 | N 130 | O 159 | S 3 | 0 | 0 |
| 3 | P | 101 | Total 760 | C 475 | N 125 | O 157 | S 3 | 0 | 0 |

- Molecule 4 is a protein called REGN3471 light chain.

| Mol | Chain | Residues | Atoms | | | | | AltConf | Trace |
|---|---|---|---|---|---|---|---|---|---|
| 4 | F | 113 | Total 873 | C 552 | N 142 | O 176 | S 3 | 0 | 0 |

*Continued on next page...*

For Manuscript Review

*Continued from previous page...*

| Mol | Chain | Residues | Atoms | | | | | AltConf | Trace |
|---|---|---|---|---|---|---|---|---|---|
| 4 | B | 113 | Total 873 | C 552 | N 142 | O 176 | S 3 | 0 | 0 |
| 4 | D | 113 | Total 873 | C 552 | N 142 | O 176 | S 3 | 0 | 0 |

- Molecule 5 is a protein called Shed GP.

| Mol | Chain | Residues | Atoms | | | | | AltConf | Trace |
|---|---|---|---|---|---|---|---|---|---|
| 5 | W | 110 | Total 862 | C 549 | N 151 | O 156 | S 6 | 0 | 0 |
| 5 | V | 110 | Total 862 | C 549 | N 151 | O 156 | S 6 | 0 | 0 |
| 5 | X | 107 | Total 839 | C 534 | N 147 | O 152 | S 6 | 0 | 0 |

There are 3 discrepancies between the modelled and reference sequences:

| Chain | Residue | Modelled | Actual | Comment | Reference |
|---|---|---|---|---|---|
| W | 544 | THR | ILE | conflict | UNP Q05320 |
| V | 544 | THR | ILE | conflict | UNP Q05320 |
| X | 544 | THR | ILE | conflict | UNP Q05320 |

- Molecule 6 is a protein called REGN3479 heavy chain.

| Mol | Chain | Residues | Atoms | | | | | AltConf | Trace |
|---|---|---|---|---|---|---|---|---|---|
| 6 | Q | 116 | Total 878 | C 549 | N 152 | O 171 | S 6 | 0 | 0 |
| 6 | M | 115 | Total 869 | C 544 | N 151 | O 168 | S 6 | 0 | 0 |
| 6 | O | 115 | Total 869 | C 544 | N 151 | O 168 | S 6 | 0 | 0 |

- Molecule 7 is a protein called REGN3470 heavy chain.

| Mol | Chain | Residues | Atoms | | | | | AltConf | Trace |
|---|---|---|---|---|---|---|---|---|---|
| 7 | G | 115 | Total 912 | C 577 | N 161 | O 169 | S 5 | 0 | 0 |
| 7 | I | 113 | Total 899 | C 569 | N 159 | O 166 | S 5 | 0 | 0 |

- Molecule 8 is a protein called REGN3470 light chain.

For Manuscript Review

| Mol | Chain | Residues | Atoms | | | | | AltConf | Trace |
|---|---|---|---|---|---|---|---|---|---|
| 8 | H | 107 | Total 810 | C 511 | N 134 | O 162 | S 3 | 0 | 0 |
| 8 | J | 95 | Total 728 | C 461 | N 119 | O 145 | S 3 | 0 | 0 |

- Molecule 9 is an oligosaccharide called alpha-D-mannopyranose-(1-6)-alpha-D-mannopyranose-(1-6)-[alpha-D-mannopyranose-(1-3)]beta-D-mannopyranose-(1-4)-2-acetamido-2-deoxy-beta-D-glucopyranose-(1-4)-2-acetamido-2-deoxy-beta-D-glucopyranose.

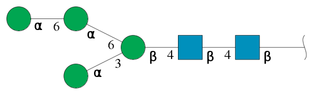

| Mol | Chain | Residues | Atoms | | | | AltConf | Trace |
|---|---|---|---|---|---|---|---|---|
| 9 | K | 6 | Total 72 | C 40 | N 2 | O 30 | 0 | 0 |
| 9 | L | 6 | Total 72 | C 40 | N 2 | O 30 | 0 | 0 |
| 9 | Y | 6 | Total 72 | C 40 | N 2 | O 30 | 0 | 0 |

- Molecule 10 is 2-acetamido-2-deoxy-beta-D-glucopyranose (three-letter code: NAG) (formula: $C_8H_{15}NO_6$) (labeled as "Ligand of Interest" by depositor).

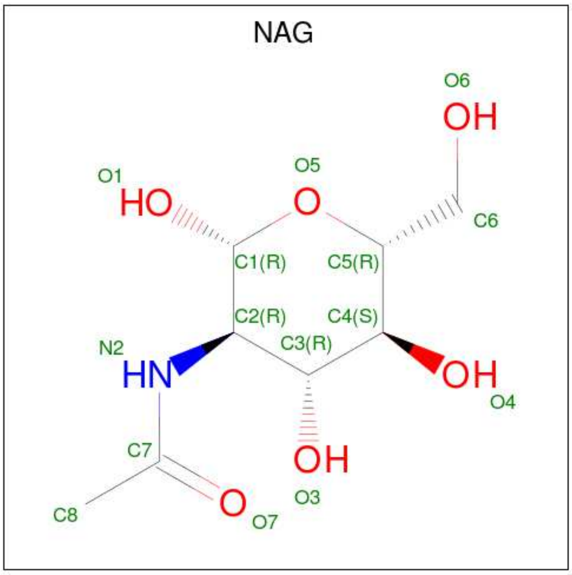


| Mol | Chain | Residues | Atoms | | | | AltConf |
|---|---|---|---|---|---|---|---|
| 10 | T | 1 | Total 42 | C 24 | N 3 | O 15 | 0 |

Continued on next page...

For Manuscript Review

*Continued from previous page...*

| Mol | Chain | Residues | Atoms | | | | AltConf |
|---|---|---|---|---|---|---|---|
| 10 | T | 1 | Total 42 | C 24 | N 3 | O 15 | 0 |
| 10 | T | 1 | Total 42 | C 24 | N 3 | O 15 | 0 |
| 10 | S | 1 | Total 28 | C 16 | N 2 | O 10 | 0 |
| 10 | S | 1 | Total 28 | C 16 | N 2 | O 10 | 0 |
| 10 | U | 1 | Total 42 | C 24 | N 3 | O 15 | 0 |
| 10 | U | 1 | Total 42 | C 24 | N 3 | O 15 | 0 |
| 10 | U | 1 | Total 42 | C 24 | N 3 | O 15 | 0 |

For Manuscript Review

### 3 Residue-property plots ⓘ

These plots are drawn for all protein, RNA, DNA and oligosaccharide chains in the entry. The first graphic for a chain summarises the proportions of the various outlier classes displayed in the second graphic. The second graphic shows the sequence view annotated by issues in geometry and atom inclusion in map density. Residues are color-coded according to the number of geometric quality criteria for which they contain at least one outlier: green = 0, yellow = 1, orange = 2 and red = 3 or more. A red diamond above a residue indicates a poor fit to the EM map for this residue (all-atom inclusion < 40%). Stretches of 2 or more consecutive residues without any outlier are shown as a green connector. Residues present in the sample, but not in the model, are shown in grey.

- Molecule 1: REGN3471 heavy chain

Chain A:

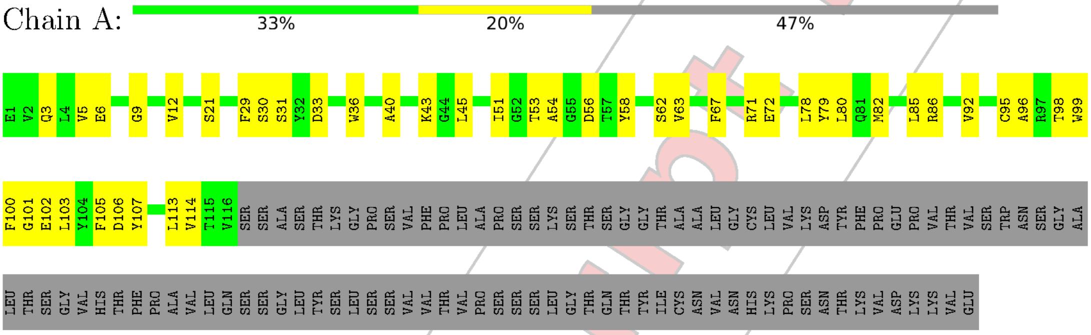


- Molecule 1: REGN3471 heavy chain

Chain C:

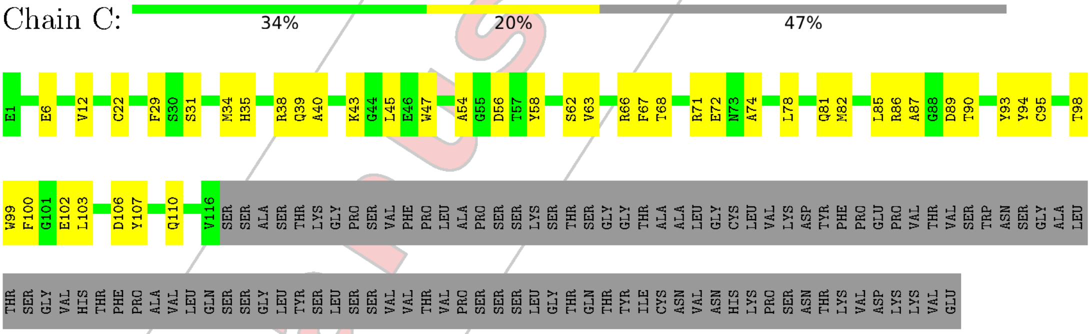


- Molecule 1: REGN3471 heavy chain

Chain E:

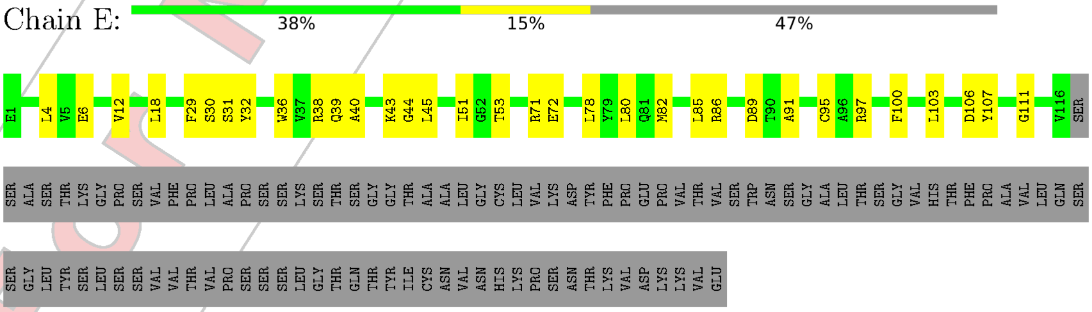


For Manuscript Review

- Molecule 2: Envelope glycoprotein

Chain T:

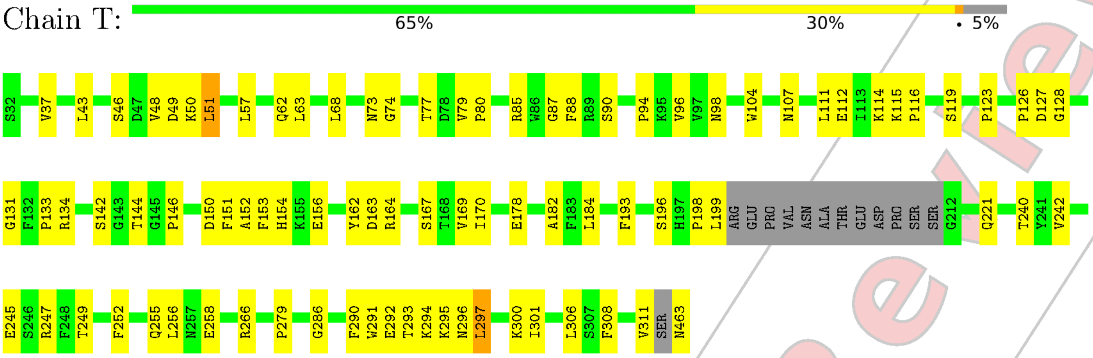


- Molecule 2: Envelope glycoprotein

Chain S:

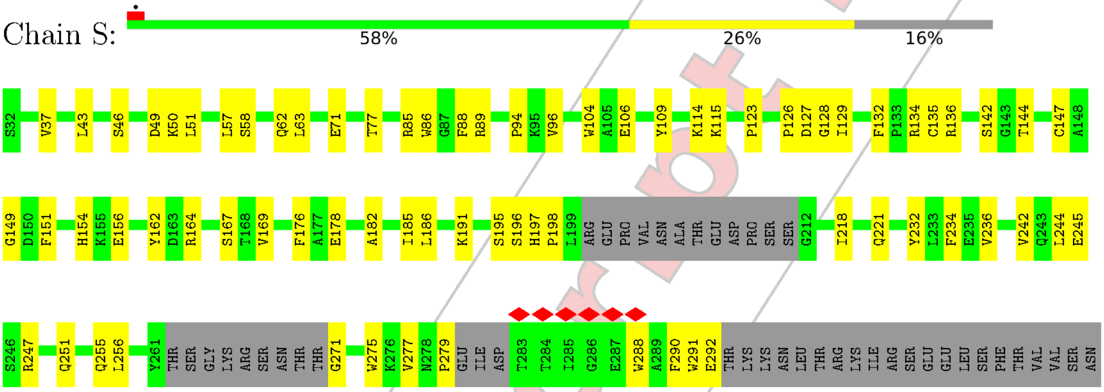


- Molecule 2: Envelope glycoprotein

Chain U:

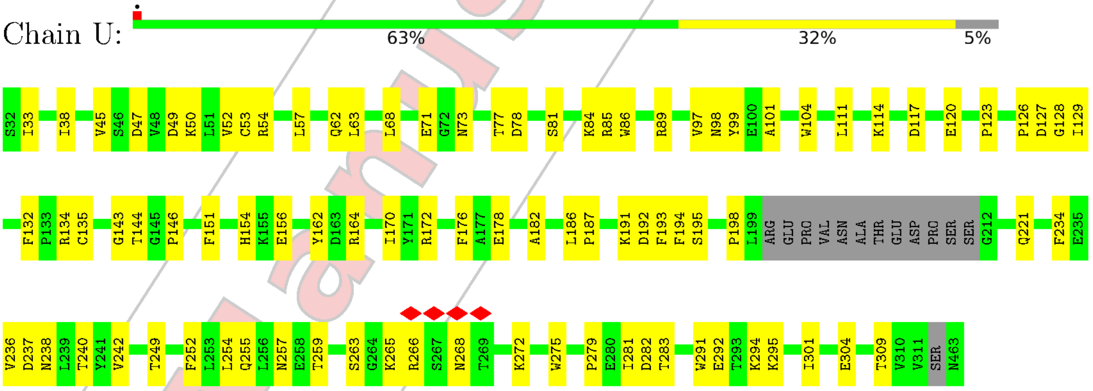


- Molecule 3: REGN3479 light chain

Chain R:

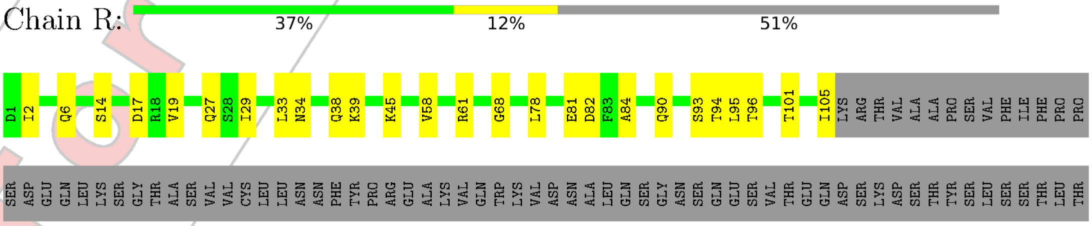


For Manuscript Review

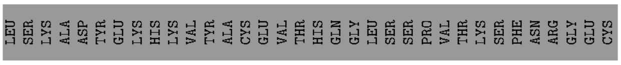

• Molecule 3: REGN3479 light chain

Chain N:

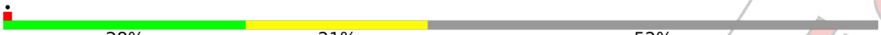


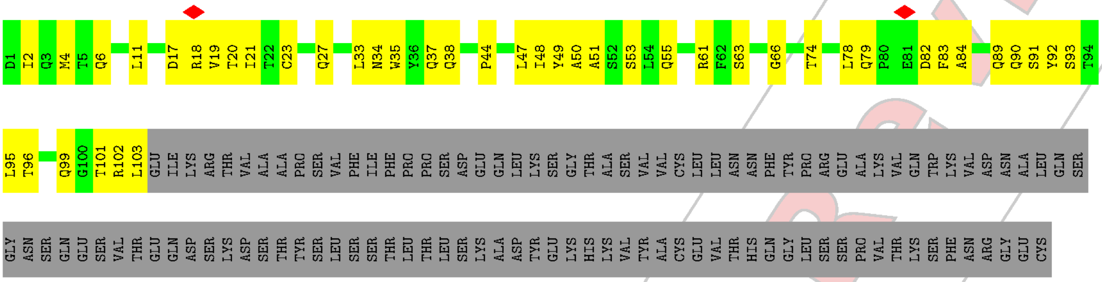

• Molecule 3: REGN3479 light chain

Chain P:

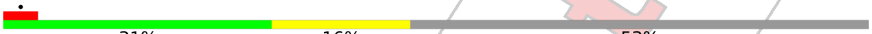


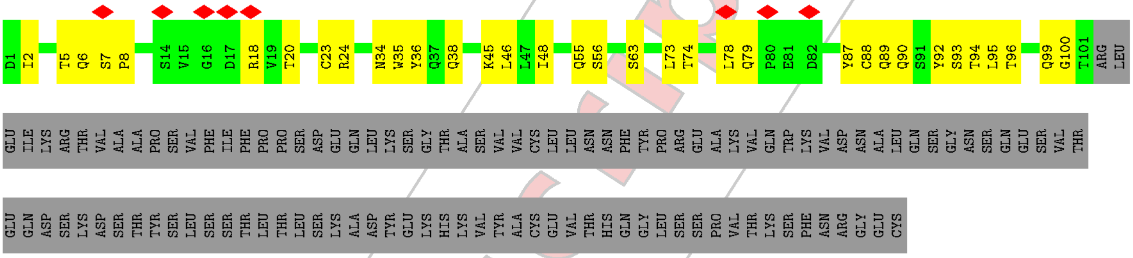

• Molecule 4: REGN3471 light chain

Chain F:

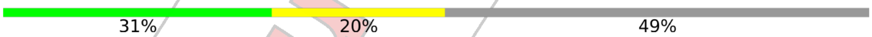


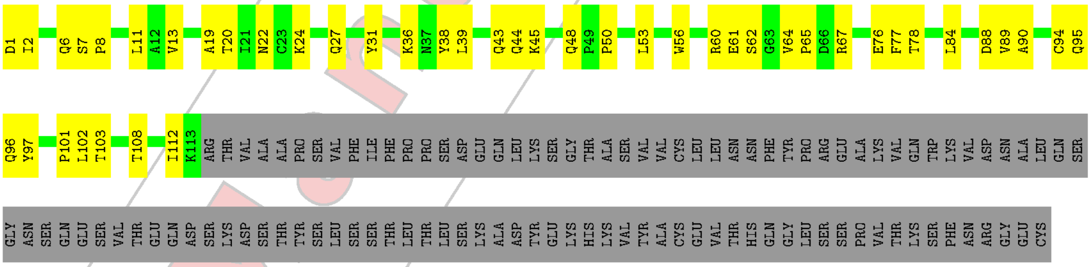

• Molecule 4: REGN3471 light chain

Chain B:

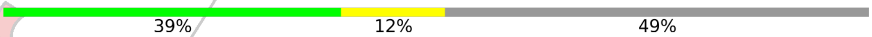


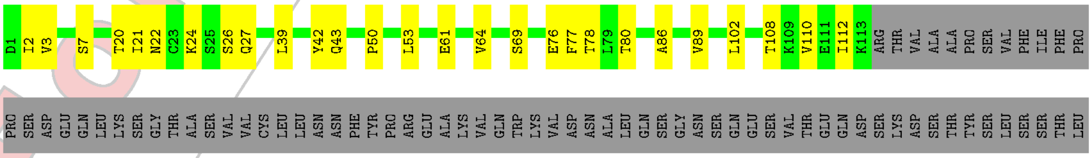

For Manuscript Review

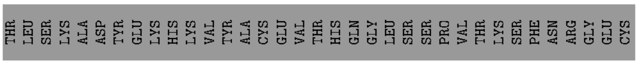


- Molecule 4: REGN3471 light chain

Chain D:

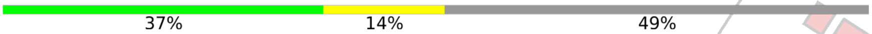


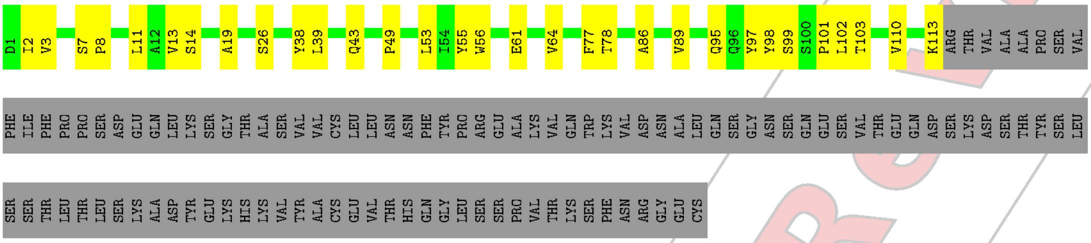


- Molecule 5: Shed GP

Chain W:

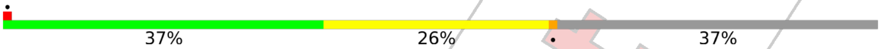


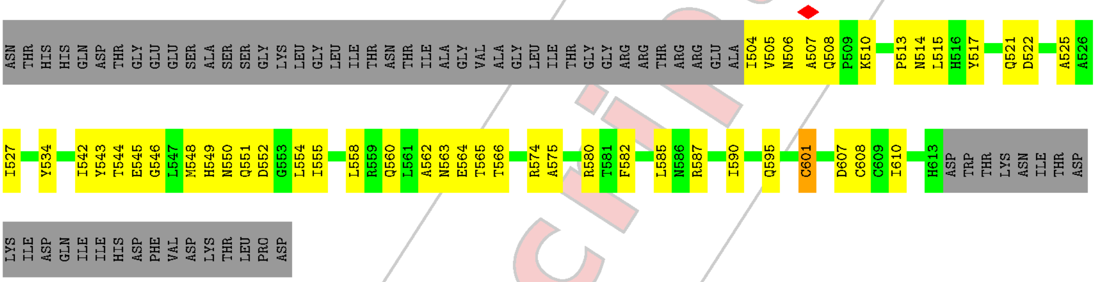


- Molecule 5: Shed GP

Chain V:

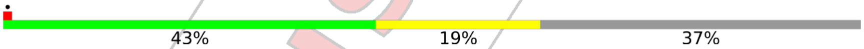


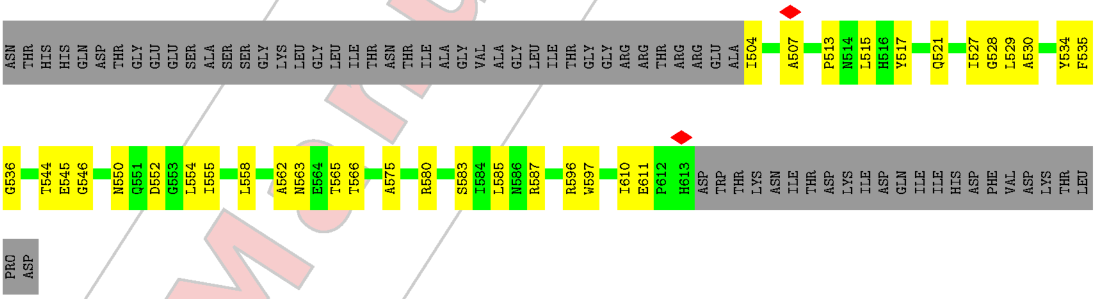


- Molecule 5: Shed GP

Chain X:

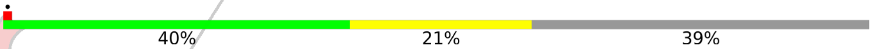


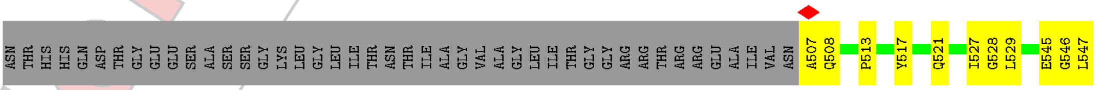


For Manuscript Review

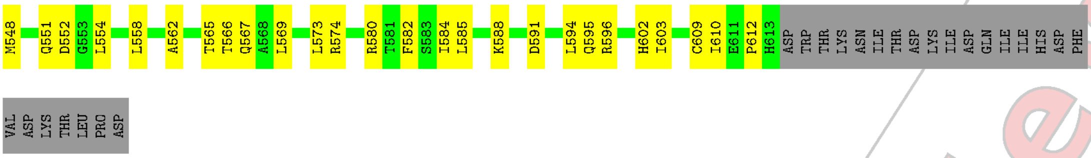


- Molecule 6: REGN3479 heavy chain

Chain Q:

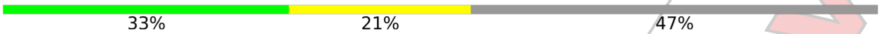


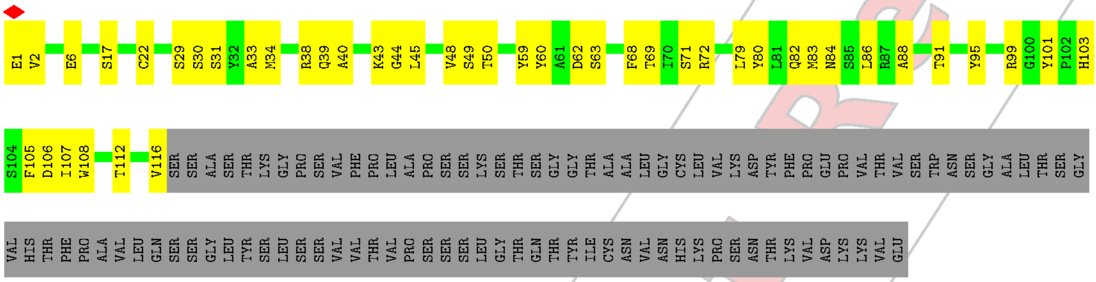


- Molecule 6: REGN3479 heavy chain

Chain M:

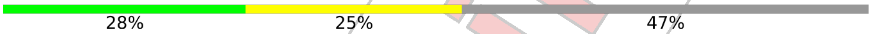


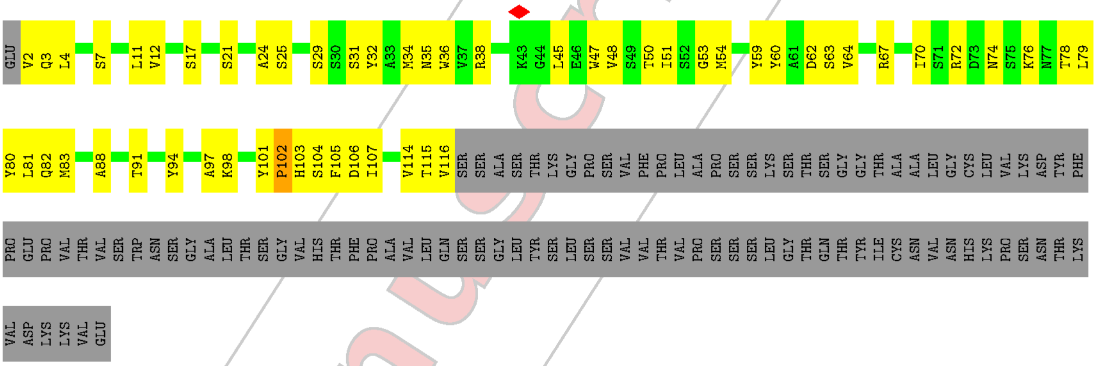


- Molecule 6: REGN3479 heavy chain

Chain O:

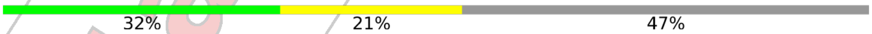


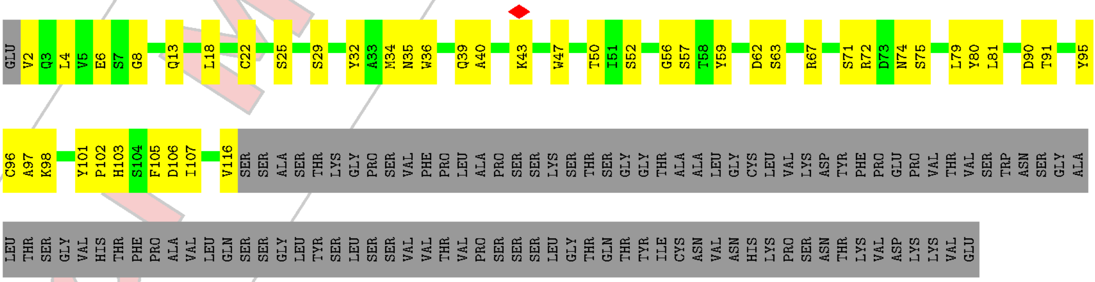


- Molecule 7: REGN3470 heavy chain

For Manuscript Review

Chain G: 31% 22% 47%

Q1 V2 Q3 L4 V5 E6 G9 Q13 R16 S17 L18 R19 F27 M34 H35 W36 V37 R38 Q39 L45 E46 W47 V50 D62 S63 V64 K65 G66 R67 F68 D73 K76 Y80 L81 Q82 M83 N84 S85 L86 R87 A88 E89 D90 T91 A92 V93 Y94 A97

R98 N99 W100 N101 L102 F103 D104 Y105 T110 L111 V112 T113 V114 S115 SER ALA SER THR LYS GLY PRO SER VAL PHE PRO LEU ALA PRO SER SER LYS SER THR SER GLY GLY THR ALA ALA LEU GLY CYS LEU VAL LYS ASP TYR PHE PRO GLU PRO VAL THR VAL SER TRP ASN SER GLY

ALA LEU THR SER GLY VAL HIS THR PHE PRO ALA VAL LEU GLN SER SER GLY LEU TYR SER LEU SER SER VAL VAL THR VAL PRO SER SER SER LEU GLY THR GLN THR TYR ILE CYS ASN VAL ASN HIS LYS PRO SER ASN THR LYS VAL ASP LYS LYS VAL GLU

- Molecule 7: REGN3470 heavy chain

Chain I: 16% 28% 24% 47%

Q1 V5 E6 S7 G8 G9 G10 V11 V12 Q13 P14 G15 R16 S17 L18 R19 C22 F27 H35 W36 V37 R38 Q39 A40 P41 G42 M43 G44 L45 E46 I51 W52 H53 D54 G55 S56 D57 Y60 A61 D62 S63 V64 K65 G66 R67 F68 T69 I70 S71 R72 D73 N74 S75 K76

N77 T78 L79 Y80 L81 Q82 M83 N84 S85 L86 R87 A88 E89 D90 T91 A92 V93 Y94 Y95 C96 A97 R98 N99 W100 N101 L102 F103 D104 Y105 W106 G107 Q108 G109 T110 L111 V112 T113 VAL SER SER ALA SER THR LYS GLY PRO SER VAL PHE PRO LEU ALA PRO SER SER LYS SER THR SER GLY

GLY THR ALA ALA LEU GLY CYS LEU VAL LYS ASP TYR PHE PRO GLU PRO VAL THR VAL SER TRP ASN SER GLY ALA LEU THR SER GLY VAL HIS THR PHE PRO ALA VAL LEU GLN SER SER GLY LEU TYR SER LEU SER SER VAL VAL THR VAL PRO SER SER SER LEU GLY THR GLN THR

TYR ILE CYS ASN VAL ASN HIS LYS PRO SER ASN THR LYS VAL ASP LYS LYS VAL GLU

- Molecule 8: REGN3470 light chain

Chain H: 26% 24% 50%

ASP I2 Q3 M4 T5 Q6 S7 P8 L11 S12 A13 S14 D17 R18 I19 T20 I21 T22 C23 R24 I29 S30 T31 Y32 L33 H34 W35 Y36 Q37 Q38 L47 I48 Q55 V58 R61 F62 S63 G66 T69 D70 F71 T72 L73 T74 S77 L78 E81 A84

C88 S91 F92 S93 T94 P95 P96 I97 Q101 G102 T103 K104 L105 K108 ARG THR VAL ALA ALA PRO SER VAL PHE ILE PHE PRO PRO SER ASP GLU GLN LEU LYS SER GLY THR ALA SER VAL VAL CYS LEU LEU ASN ASN PHE TYR PRO ARG GLU ALA LYS VAL GLN TRP LYS

VAL ASP ASN ALA LEU GLN SER GLY ASN SER GLN GLU SER VAL THR GLU GLN ASP SER LYS ASP SER THR TYR SER LEU SER SER THR LEU THR LEU SER LYS ALA ASP TYR GLU LYS HIS LYS VAL TYR ALA CYS GLU VAL THR HIS GLN GLY LEU SER SER PRO VAL THR LYS SER PHE

ASN ARG GLY GLU CYS

- Molecule 8: REGN3470 light chain

Chain J: 25% 21% 22% • 56%

ASP I2 Q3 M4 T5 Q6 S7 P8 S9 S10 LEU SER ALA SER VAL GLY ASP ARG ILE THR ILE T22 C23 R24 A25 S26 Q27 S28 I29 Y32 L33 H34 W35 Y36 Q37 Q38 K39 P40 G41 K42 A43 P44 L47 I48 Y49 A50 A51 S52 T53 L54 Q55 S56 G57 V58 P59 S60 R61 F62

For Manuscript Review

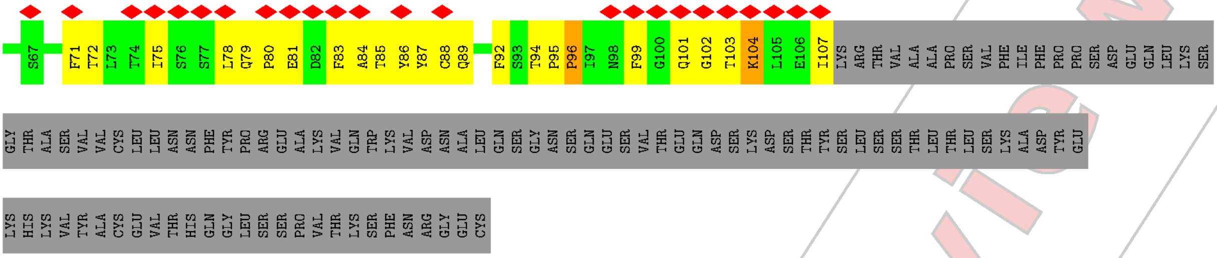


- Molecule 9: alpha-D-mannopyranose-(1-6)-alpha-D-mannopyranose-(1-6)-[alpha-D-mannopyranose-(1-3)]beta-D-mannopyranose-(1-4)-2-acetamido-2-deoxy-beta-D-glucopyranose-(1-4)-2-acetamido-2-deoxy-beta-D-glucopyranose

Chain K:

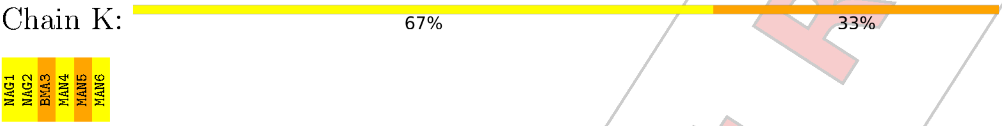


- Molecule 9: alpha-D-mannopyranose-(1-6)-alpha-D-mannopyranose-(1-6)-[alpha-D-mannopyranose-(1-3)]beta-D-mannopyranose-(1-4)-2-acetamido-2-deoxy-beta-D-glucopyranose-(1-4)-2-acetamido-2-deoxy-beta-D-glucopyranose

Chain L:

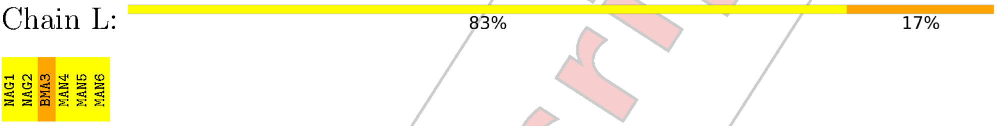


- Molecule 9: alpha-D-mannopyranose-(1-6)-alpha-D-mannopyranose-(1-6)-[alpha-D-mannopyranose-(1-3)]beta-D-mannopyranose-(1-4)-2-acetamido-2-deoxy-beta-D-glucopyranose-(1-4)-2-acetamido-2-deoxy-beta-D-glucopyranose

Chain Y:

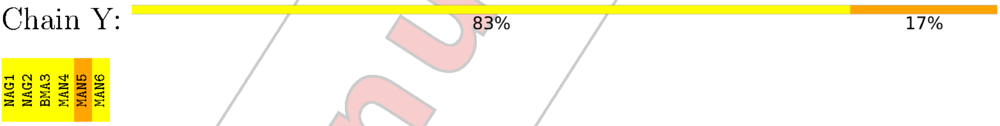


For Manuscript Review

### 4 Experimental information (i)

| Property | Value | Source |
|---|---|---|
| EM reconstruction method | SINGLE PARTICLE | Depositor |
| Imposed symmetry | POINT, C1 | Depositor |
| Number of particles used | 276715 | Depositor |
| Resolution determination method | FSC 0.143 CUT-OFF | Depositor |
| CTF correction method | PHASE FLIPPING AND AMPLITUDE CORRECTION | Depositor |
| Microscope | FEI TITAN KRIOS | Depositor |
| Voltage (kV) | 300 | Depositor |
| Electron dose ($e^-/\text{Å}^2$) | 52.4 | Depositor |
| Minimum defocus (nm) | 800 | Depositor |
| Maximum defocus (nm) | 2600 | Depositor |
| Magnification | Not provided | |
| Image detector | GATAN K2 SUMMIT (4k x 4k) | Depositor |
| Maximum map value | 1.718 | Depositor |
| Minimum map value | -0.619 | Depositor |
| Average map value | -0.000 | Depositor |
| Map value standard deviation | 0.030 | Depositor |
| Recommended contour level | 0.25 | Depositor |
| Map size (Å) | 537.8048, 537.8048, 537.8048 | wwPDB |
| Map dimensions | 512, 512, 512 | wwPDB |
| Map angles (°) | 90.0, 90.0, 90.0 | wwPDB |
| Pixel spacing (Å) | 1.0504, 1.0504, 1.0504 | Depositor |

For Manuscript Review

### 5 Model quality (i)

#### 5.1 Standard geometry (i)

Bond lengths and bond angles in the following residue types are not validated in this section: BMA, NAG, MAN

The Z score for a bond length (or angle) is the number of standard deviations the observed value is removed from the expected value. A bond length (or angle) with $|Z| > 5$ is considered an outlier worth inspection. RMSZ is the root-mean-square of all Z scores of the bond lengths (or angles).

| Mol | Chain | Bond lengths | | Bond angles | |
|---|---|---|---|---|---|
| | | RMSZ | $\#\|Z\| > 5$ | RMSZ | $\#\|Z\| > 5$ |
| 1 | A | 0.44 | 0/918 | 0.52 | 0/1245 |
| 1 | C | 0.41 | 0/918 | 0.52 | 0/1245 |
| 1 | E | 0.43 | 0/918 | 0.51 | 0/1245 |
| 2 | S | 0.38 | 0/1921 | 0.49 | 0/2609 |
| 2 | T | 0.40 | 0/2179 | 0.54 | 2/2958 (0.1%) |
| 2 | U | 0.40 | 1/2178 (0.0%) | 0.52 | 1/2955 (0.0%) |
| 3 | N | 0.29 | 0/795 | 0.53 | 0/1078 |
| 3 | P | 0.31 | 0/776 | 0.52 | 0/1053 |
| 3 | R | 0.34 | 0/812 | 0.55 | 0/1101 |
| 4 | B | 0.40 | 0/893 | 0.54 | 0/1214 |
| 4 | D | 0.41 | 0/893 | 0.53 | 0/1214 |
| 4 | F | 0.42 | 0/893 | 0.57 | 0/1214 |
| 5 | V | 0.37 | 0/884 | 0.56 | 0/1205 |
| 5 | W | 0.42 | 0/884 | 0.56 | 0/1205 |
| 5 | X | 0.40 | 0/861 | 0.55 | 0/1173 |
| 6 | M | 0.32 | 0/888 | 0.53 | 0/1201 |
| 6 | O | 0.32 | 0/888 | 0.50 | 0/1201 |
| 6 | Q | 0.33 | 0/897 | 0.51 | 0/1213 |
| 7 | G | 0.33 | 0/936 | 0.50 | 0/1271 |
| 7 | I | 0.33 | 0/923 | 0.59 | 1/1253 (0.1%) |
| 8 | H | 0.31 | 0/829 | 0.56 | 0/1126 |
| 8 | J | 0.29 | 0/746 | 0.52 | 0/1012 |
| All | All | 0.37 | 1/22830 (0.0%) | 0.53 | 4/30991 (0.0%) |

All (1) bond length outliers are listed below:

| Mol | Chain | Res | Type | Atoms | Z | Observed(Å) | Ideal(Å) |
|---|---|---|---|---|---|---|---|
| 2 | U | 53 | CYS | CB-SG | -5.23 | 1.73 | 1.81 |

All (4) bond angle outliers are listed below:

For Manuscript Review

| Mol | Chain | Res | Type | Atoms | Z | Observed(°) | Ideal(°) |
|---|---|---|---|---|---|---|---|
| 2 | T | 51 | LEU | CA-CB-CG | 7.78 | 133.19 | 115.30 |
| 2 | U | 53 | CYS | CA-CB-SG | 6.24 | 125.23 | 114.00 |
| 7 | I | 76 | LYS | CB-CG-CD | 5.38 | 125.59 | 111.60 |
| 2 | T | 297 | LEU | CA-CB-CG | 5.22 | 127.32 | 115.30 |

There are no chirality outliers.

There are no planarity outliers.

#### 5.2 Too-close contacts (i)

In the following table, the Non-H and H(model) columns list the number of non-hydrogen atoms and hydrogen atoms in the chain respectively. The H(added) column lists the number of hydrogen atoms added and optimized by MolProbity. The Clashes column lists the number of clashes within the asymmetric unit, whereas Symm-Clashes lists symmetry-related clashes.

| Mol | Chain | Non-H | H(model) | H(added) | Clashes | Symm-Clashes |
|---|---|---|---|---|---|---|
| 1 | A | 895 | 0 | 851 | 35 | 0 |
| 1 | C | 895 | 0 | 851 | 31 | 0 |
| 1 | E | 895 | 0 | 851 | 28 | 0 |
| 2 | S | 1872 | 0 | 1810 | 55 | 0 |
| 2 | T | 2127 | 0 | 2079 | 77 | 0 |
| 2 | U | 2127 | 0 | 2077 | 80 | 0 |
| 3 | N | 779 | 0 | 758 | 33 | 0 |
| 3 | P | 760 | 0 | 734 | 26 | 0 |
| 3 | R | 796 | 0 | 775 | 19 | 0 |
| 4 | B | 873 | 0 | 848 | 19 | 0 |
| 4 | D | 873 | 0 | 848 | 20 | 0 |
| 4 | F | 873 | 0 | 848 | 35 | 0 |
| 5 | V | 862 | 0 | 835 | 38 | 0 |
| 5 | W | 862 | 0 | 836 | 57 | 0 |
| 5 | X | 839 | 0 | 809 | 44 | 0 |
| 6 | M | 869 | 0 | 837 | 57 | 0 |
| 6 | O | 869 | 0 | 837 | 42 | 0 |
| 6 | Q | 878 | 0 | 846 | 40 | 0 |
| 7 | G | 912 | 0 | 864 | 37 | 0 |
| 7 | I | 899 | 0 | 850 | 51 | 0 |
| 8 | H | 810 | 0 | 789 | 45 | 0 |
| 8 | J | 728 | 0 | 703 | 45 | 0 |
| 9 | K | 72 | 0 | 61 | 7 | 0 |
| 9 | L | 72 | 0 | 61 | 4 | 0 |
| 9 | Y | 72 | 0 | 61 | 4 | 0 |
| 10 | S | 28 | 0 | 26 | 0 | 0 |

*Continued on next page...*

For Manuscript Review

Continued from previous page...

| Mol | Chain | Non-H | H(model) | H(added) | Clashes | Symm-Clashes |
|---|---|---|---|---|---|---|
| 10 | T | 42 | 0 | 39 | 1 | 0 |
| 10 | U | 42 | 0 | 39 | 0 | 0 |
| All | All | 22621 | 0 | 21823 | 786 | 0 |

The all-atom clashscore is defined as the number of clashes found per 1000 atoms (including hydrogen atoms). The all-atom clashscore for this structure is 18.

All (786) close contacts within the same asymmetric unit are listed below, sorted by their clash magnitude.

| Atom-1 | Atom-2 | Interatomic distance (Å) | Clash overlap (Å) |
|---|---|---|---|
| 6:O:57:SER:HB3 | 6:O:59:TYR:HE1 | 1.31 | 0.94 |
| 2:T:73:ASN:HB3 | 5:W:508:GLN:HE21 | 1.34 | 0.92 |
| 7:I:17:SER:HB3 | 7:I:83:MET:HG3 | 1.53 | 0.91 |
| 2:S:104:TRP:NE1 | 5:V:545:GLU:OE1 | 2.06 | 0.88 |
| 2:S:128:GLY:HA3 | 5:V:580:ARG:HH22 | 1.37 | 0.86 |
| 4:B:24:LYS:NZ | 4:B:76:GLU:OE1 | 2.08 | 0.85 |
| 8:H:5:THR:O | 8:H:24:ARG:N | 2.09 | 0.84 |
| 1:A:86:ARG:HD3 | 4:F:20:THR:HG21 | 1.60 | 0.83 |
| 7:I:16:ARG:HE | 7:I:17:SER:H | 1.22 | 0.83 |
| 5:W:505:VAL:HG22 | 5:W:560:GLN:HB2 | 1.62 | 0.82 |
| 2:U:237:ASP:OD1 | 2:U:238:ASN:N | 2.12 | 0.81 |
| 2:U:128:GLY:HA3 | 5:X:580:ARG:HH12 | 1.44 | 0.81 |
| 3:N:6:GLN:NE2 | 3:N:99:GLN:O | 2.11 | 0.81 |
| 8:J:36:TYR:HB2 | 8:J:87:TYR:HB2 | 1.63 | 0.81 |
| 6:O:52:SER:HB3 | 6:O:57:SER:HB2 | 1.62 | 0.80 |
| 6:Q:33:ALA:HB3 | 6:Q:99:ARG:HD2 | 1.65 | 0.79 |
| 7:I:36:TRP:HE1 | 7:I:79:LEU:HD21 | 1.45 | 0.79 |
| 7:G:13:GLN:H | 7:G:16:ARG:HH21 | 1.31 | 0.78 |
| 1:C:62:SER:N | 4:B:76:GLU:OE2 | 2.18 | 0.77 |
| 5:W:527:ILE:HD13 | 6:Q:59:TYR:HD1 | 1.50 | 0.77 |
| 5:W:574:ARG:HH21 | 5:X:521:GLN:HE22 | 1.30 | 0.77 |
| 8:H:61:ARG:NH2 | 8:H:81:GLU:OE2 | 2.18 | 0.77 |
| 1:E:40:ALA:HB3 | 1:E:43:LYS:HB3 | 1.67 | 0.76 |
| 2:T:294:LYS:HE2 | 2:T:296:ASN:HD22 | 1.51 | 0.76 |
| 3:N:63:SER:OG | 3:N:74:THR:OG1 | 2.02 | 0.76 |
| 5:V:566:THR:HG21 | 9:L:1:NAG:H82 | 1.68 | 0.76 |
| 7:G:87:ARG:NH2 | 7:G:89:GLU:OE2 | 2.19 | 0.74 |
| 1:A:82:MET:HB3 | 1:A:85:LEU:HD21 | 1.69 | 0.74 |
| 2:S:106:GLU:OE2 | 2:S:136:ARG:NH2 | 2.20 | 0.74 |
| 2:T:73:ASN:HB3 | 5:W:508:GLN:NE2 | 2.03 | 0.74 |
| 6:M:36:TRP:HE1 | 6:M:79:LEU:HD22 | 1.52 | 0.74 |

Continued on next page...

For Manuscript Review

Continued from previous page...

| Atom-1 | Atom-2 | Interatomic distance (Å) | Clash overlap (Å) |
|---|---|---|---|
| 2:T:127:ASP:OD1 | 2:T:128:GLY:N | 2.21 | 0.74 |
| 2:S:251:GLN:O | 2:S:255:GLN:NE2 | 2.21 | 0.73 |
| 2:U:85:ARG:NH1 | 2:U:178:GLU:OE2 | 2.21 | 0.73 |
| 6:M:29:SER:O | 6:M:72:ARG:NH2 | 2.21 | 0.73 |
| 1:C:6:GLU:OE2 | 1:C:95:CYS:N | 2.17 | 0.73 |
| 3:N:90:GLN:HE21 | 3:N:96:THR:H | 1.35 | 0.73 |
| 2:U:127:ASP:OD2 | 2:U:128:GLY:N | 2.22 | 0.73 |
| 2:T:51:LEU:HB3 | 5:W:595:GLN:NE2 | 2.04 | 0.73 |
| 5:W:564:GLU:OE1 | 6:M:53:GLY:N | 2.21 | 0.73 |
| 2:U:68:LEU:HD11 | 5:X:558:LEU:HG | 1.71 | 0.73 |
| 7:I:53:HIS:O | 7:I:72:ARG:NH1 | 2.22 | 0.73 |
| 1:A:45:LEU:HD21 | 4:B:50:PRO:HG2 | 1.70 | 0.72 |
| 1:A:40:ALA:HB3 | 1:A:43:LYS:HB3 | 1.71 | 0.72 |
| 7:G:39:GLN:HA | 7:G:45:LEU:HA | 1.70 | 0.72 |
| 7:I:35:HIS:HB2 | 7:I:97:ALA:HB3 | 1.70 | 0.72 |
| 8:H:18:ARG:HH12 | 8:H:74:THR:HB | 1.55 | 0.72 |
| 1:C:82:MET:HB3 | 1:C:85:LEU:HD21 | 1.72 | 0.71 |
| 7:G:19:ARG:HD2 | 7:G:80:TYR:HB3 | 1.71 | 0.71 |
| 5:X:603:ILE:HG13 | 5:X:610:ILE:HD11 | 1.72 | 0.71 |
| 1:A:3:GLN:HE22 | 1:A:5:VAL:HG22 | 1.55 | 0.70 |
| 3:P:35:TRP:HB2 | 3:P:48:ILE:HB | 1.73 | 0.70 |
| 5:W:513:PRO:O | 5:W:555:ILE:HD11 | 1.90 | 0.70 |
| 1:E:12:VAL:HG11 | 1:E:18:LEU:HB2 | 1.73 | 0.70 |
| 2:T:301:ILE:HD12 | 2:T:306:LEU:HD21 | 1.74 | 0.70 |
| 3:N:49:TYR:HB2 | 3:N:53:SER:HB2 | 1.74 | 0.70 |
| 6:O:2:VAL:N | 6:O:25:SER:O | 2.23 | 0.70 |
| 1:E:38:ARG:NH1 | 1:E:89:ASP:OD1 | 2.24 | 0.70 |
| 1:A:62:SER:N | 4:F:76:GLU:OE1 | 2.25 | 0.70 |
| 8:J:95:PRO:HD2 | 8:J:96:PRO:HD3 | 1.73 | 0.70 |
| 8:H:32:TYR:HB3 | 8:H:91:SER:HB3 | 1.73 | 0.69 |
| 6:M:91:THR:HB | 6:M:116:VAL:HG22 | 1.75 | 0.69 |
| 6:O:57:SER:HB3 | 6:O:59:TYR:CE1 | 2.22 | 0.69 |
| 5:X:566:THR:HG21 | 9:Y:1:NAG:H82 | 1.75 | 0.69 |
| 6:O:35:ASN:HD22 | 6:O:105:PHE:HE1 | 1.41 | 0.69 |
| 2:S:242:VAL:HG12 | 2:S:275:TRP:HB2 | 1.75 | 0.69 |
| 2:T:306:LEU:O | 7:G:101:ASN:ND2 | 2.26 | 0.69 |
| 2:U:81:SER:HA | 2:U:84:LYS:HE2 | 1.75 | 0.69 |
| 3:P:34:ASN:HB2 | 3:P:89:GLN:HB3 | 1.74 | 0.69 |
| 2:U:111:LEU:HD12 | 2:U:170:ILE:HG21 | 1.74 | 0.68 |
| 2:U:38:ILE:HG21 | 2:U:193:PHE:HE1 | 1.56 | 0.68 |
| 6:O:50:THR:HB | 6:O:59:TYR:HB2 | 1.75 | 0.68 |

Continued on next page...

For Manuscript Review

*Continued from previous page...*

| Atom-1 | Atom-2 | Interatomic distance (Å) | Clash overlap (Å) |
|---|---|---|---|
| 5:W:566:THR:HG21 | 9:K:1:NAG:H82 | 1.76 | 0.68 |
| 2:U:282:ASP:OD1 | 2:U:283:THR:N | 2.27 | 0.68 |
| 5:X:596:ARG:NH1 | 5:X:596:ARG:HB2 | 2.09 | 0.68 |
| 7:G:35:HIS:HB2 | 7:G:97:ALA:HB3 | 1.74 | 0.67 |
| 4:B:2:ILE:HG13 | 4:B:27:GLN:HB2 | 1.76 | 0.67 |
| 4:F:50:PRO:HG2 | 1:E:45:LEU:HD21 | 1.75 | 0.67 |
| 2:S:271:GLY:N | 4:F:62:SER:HG | 1.93 | 0.66 |
| 2:U:182:ALA:HB2 | 5:X:562:ALA:HB2 | 1.78 | 0.66 |
| 2:T:111:LEU:HD12 | 2:T:170:ILE:HG21 | 1.77 | 0.66 |
| 3:R:90:GLN:HE21 | 3:R:96:THR:H | 1.44 | 0.66 |
| 1:E:43:LYS:HG2 | 1:E:44:GLY:H | 1.58 | 0.66 |
| 2:S:86:TRP:NE1 | 2:S:178:GLU:OE2 | 2.29 | 0.66 |
| 8:J:6:GLN:HE21 | 8:J:103:THR:HG23 | 1.60 | 0.66 |
| 1:C:22:CYS:HB3 | 1:C:78:LEU:HB3 | 1.78 | 0.65 |
| 2:T:182:ALA:HB2 | 5:W:562:ALA:HB2 | 1.77 | 0.65 |
| 2:S:58:SER:OG | 2:S:62:GLN:NE2 | 2.29 | 0.65 |
| 6:Q:106:ASP:OD1 | 6:Q:107:ILE:N | 2.30 | 0.65 |
| 8:H:37:GLN:HB2 | 8:H:47:LEU:HD11 | 1.78 | 0.65 |
| 2:T:48:VAL:O | 5:W:595:GLN:NE2 | 2.30 | 0.65 |
| 2:S:288:TRP:HB3 | 2:S:292:GLU:HB2 | 1.79 | 0.65 |
| 3:P:6:GLN:O | 3:P:99:GLN:NE2 | 2.30 | 0.65 |
| 2:S:43:LEU:HD22 | 5:V:558:LEU:HD23 | 1.78 | 0.65 |
| 6:Q:62:ASP:OD1 | 6:Q:63:SER:N | 2.30 | 0.65 |
| 8:H:63:SER:O | 8:H:74:THR:OG1 | 2.10 | 0.65 |
| 7:I:16:ARG:NE | 7:I:17:SER:H | 1.94 | 0.64 |
| 8:J:6:GLN:NE2 | 8:J:101:GLN:O | 2.30 | 0.64 |
| 2:U:120:GLU:OE2 | 2:U:172:ARG:NH1 | 2.30 | 0.64 |
| 1:E:12:VAL:HG21 | 1:E:85:LEU:HD22 | 1.79 | 0.64 |
| 3:N:90:GLN:NE2 | 3:N:93:SER:O | 2.29 | 0.64 |
| 6:Q:6:GLU:HA | 6:Q:22:CYS:HA | 1.80 | 0.64 |
| 6:M:2:VAL:N | 6:M:25:SER:O | 2.31 | 0.64 |
| 6:M:36:TRP:HD1 | 6:M:70:ILE:HD11 | 1.61 | 0.64 |
| 6:O:6:GLU:OE2 | 6:O:96:CYS:N | 2.31 | 0.64 |
| 3:N:44:PRO:HG2 | 6:M:45:LEU:HD21 | 1.79 | 0.64 |
| 6:M:35:ASN:HD22 | 6:M:105:PHE:HE1 | 1.43 | 0.64 |
| 6:Q:40:ALA:HB3 | 6:Q:43:LYS:HB3 | 1.81 | 0.63 |
| 6:M:101:TYR:CD2 | 6:M:102:PRO:HD3 | 2.33 | 0.63 |
| 8:H:18:ARG:NH1 | 8:H:19:ILE:O | 2.32 | 0.63 |
| 1:A:33:ASP:OD1 | 1:A:98:THR:OG1 | 2.17 | 0.63 |
| 5:X:551:GLN:HG3 | 5:X:554:LEU:H | 1.63 | 0.63 |
| 6:M:7:SER:HG | 6:M:21:SER:HG | 1.41 | 0.63 |

*Continued on next page...*

For Manuscript Review

*Continued from previous page...*

| Atom-1 | Atom-2 | Interatomic distance (Å) | Clash overlap (Å) |
|---|---|---|---|
| 1:A:6:GLU:OE2 | 1:A:95:CYS:N | 2.25 | 0.62 |
| 2:S:291:TRP:CD1 | 5:V:513:PRO:HD2 | 2.34 | 0.62 |
| 2:U:154:HIS:NE2 | 2:U:178:GLU:OE1 | 2.29 | 0.62 |
| 3:P:90:GLN:NE2 | 3:P:93:SER:O | 2.33 | 0.62 |
| 5:V:513:PRO:O | 5:V:555:ILE:HD11 | 1.99 | 0.62 |
| 4:B:53:LEU:HA | 4:B:64:VAL:HG21 | 1.82 | 0.62 |
| 1:A:9:GLY:HA2 | 1:A:114:VAL:HG12 | 1.82 | 0.62 |
| 8:H:78:LEU:HD11 | 8:H:105:LEU:HD21 | 1.82 | 0.61 |
| 7:I:99:ASN:HB3 | 7:I:103:PHE:H | 1.65 | 0.61 |
| 3:R:95:LEU:HD21 | 6:Q:103:HIS:ND1 | 2.16 | 0.61 |
| 6:O:106:ASP:OD2 | 6:O:107:ILE:N | 2.34 | 0.61 |
| 6:Q:83:MET:HG3 | 6:Q:86:LEU:HD11 | 1.81 | 0.61 |
| 6:M:34:MET:HB2 | 6:M:79:LEU:HD12 | 1.82 | 0.61 |
| 1:C:40:ALA:HB3 | 1:C:43:LYS:HB2 | 1.81 | 0.61 |
| 7:G:99:ASN:HB3 | 7:G:102:LEU:O | 2.01 | 0.60 |
| 8:J:95:PRO:HD2 | 8:J:96:PRO:CD | 2.31 | 0.60 |
| 2:T:293:THR:O | 2:T:294:LYS:HG2 | 2.00 | 0.60 |
| 4:F:97:TYR:HB2 | 1:E:103:LEU:HB2 | 1.83 | 0.60 |
| 6:M:101:TYR:HE1 | 9:K:2:NAG:H61 | 1.66 | 0.60 |
| 8:J:49:TYR:HE1 | 8:J:55:GLN:HG3 | 1.67 | 0.60 |
| 2:T:104:TRP:NE1 | 5:W:545:GLU:OE2 | 2.35 | 0.60 |
| 6:M:35:ASN:HD21 | 6:M:103:HIS:CE1 | 2.20 | 0.60 |
| 8:H:14:SER:HB3 | 8:H:17:ASP:HB2 | 1.84 | 0.60 |
| 6:M:70:ILE:HD12 | 6:M:81:LEU:HD13 | 1.84 | 0.60 |
| 3:N:49:TYR:CD2 | 9:K:5:MAN:H61 | 2.36 | 0.60 |
| 8:H:4:MET:HG3 | 8:H:23:CYS:SG | 2.42 | 0.60 |
| 7:I:12:VAL:HG21 | 7:I:18:LEU:HD22 | 1.82 | 0.60 |
| 2:S:197:HIS:HB3 | 5:V:546:GLY:HA2 | 1.83 | 0.60 |
| 7:I:27:PHE:HB2 | 7:I:98:ARG:HH21 | 1.67 | 0.59 |
| 6:M:32:TYR:CG | 6:M:98:LYS:HD2 | 2.36 | 0.59 |
| 7:I:36:TRP:NE1 | 7:I:81:LEU:HB2 | 2.17 | 0.59 |
| 7:G:103:PHE:HZ | 8:H:97:ILE:HD12 | 1.66 | 0.59 |
| 1:C:39:GLN:HE21 | 1:C:94:TYR:HE1 | 1.50 | 0.59 |
| 2:U:291:TRP:HA | 2:U:294:LYS:HE2 | 1.82 | 0.59 |
| 6:M:36:TRP:NE1 | 6:M:79:LEU:HD22 | 2.17 | 0.59 |
| 2:U:73:ASN:HB3 | 5:X:508:GLN:NE2 | 2.17 | 0.59 |
| 2:U:198:PRO:HG2 | 5:X:548:MET:HG2 | 1.85 | 0.59 |
| 3:N:19:VAL:HG12 | 3:N:78:LEU:HD21 | 1.83 | 0.59 |
| 2:T:79:VAL:HG13 | 2:T:80:PRO:HD3 | 1.84 | 0.59 |
| 2:T:74:GLY:HA3 | 5:W:510:LYS:HB2 | 1.84 | 0.59 |
| 6:M:106:ASP:OD1 | 6:M:107:ILE:N | 2.35 | 0.58 |

*Continued on next page...*

For Manuscript Review

*Continued from previous page...*

| Atom-1 | Atom-2 | Interatomic distance (Å) | Clash overlap (Å) |
|---|---|---|---|
| 5:W:504:ILE:HG12 | 5:W:505:VAL:H | 1.67 | 0.58 |
| 5:W:580:ARG:HB3 | 5:W:582:PHE:CE2 | 2.38 | 0.58 |
| 7:I:18:LEU:O | 7:I:83:MET:HG2 | 2.02 | 0.58 |
| 3:R:61:ARG:NH1 | 3:R:82:ASP:OD1 | 2.36 | 0.58 |
| 6:Q:31:SER:HB2 | 5:V:507:ALA:H | 1.68 | 0.58 |
| 6:M:64:VAL:HG12 | 6:M:67:ARG:HH11 | 1.68 | 0.58 |
| 1:C:106:ASP:OD2 | 1:C:107:TYR:N | 2.37 | 0.58 |
| 2:U:128:GLY:H | 5:X:580:ARG:HH22 | 1.50 | 0.58 |
| 3:N:2:ILE:HG22 | 3:N:27:GLN:HB2 | 1.85 | 0.58 |
| 7:G:13:GLN:N | 7:G:16:ARG:HH21 | 2.02 | 0.58 |
| 6:M:51:ILE:HD11 | 6:M:70:ILE:HG12 | 1.85 | 0.57 |
| 1:E:72:GLU:O | 1:E:72:GLU:HG2 | 2.04 | 0.57 |
| 5:X:517:TYR:CZ | 5:X:546:GLY:HA3 | 2.38 | 0.57 |
| 3:P:95:LEU:HD11 | 6:O:103:HIS:CD2 | 2.39 | 0.57 |
| 1:E:29:PHE:O | 1:E:71:ARG:NH2 | 2.38 | 0.57 |
| 4:B:39:LEU:HD13 | 4:B:77:PHE:CD2 | 2.40 | 0.57 |
| 1:A:82:MET:HE3 | 1:A:85:LEU:HD11 | 1.87 | 0.57 |
| 3:R:14:SER:HA | 3:R:105:ILE:HD13 | 1.87 | 0.57 |
| 1:A:30:SER:O | 1:A:53:THR:OG1 | 2.13 | 0.57 |
| 6:O:72:ARG:HA | 6:O:79:LEU:HA | 1.87 | 0.57 |
| 6:M:62:ASP:OD1 | 6:M:63:SER:N | 2.38 | 0.56 |
| 2:T:255:GLN:NE2 | 2:T:301:ILE:O | 2.38 | 0.56 |
| 2:U:242:VAL:HG12 | 2:U:275:TRP:HB2 | 1.86 | 0.56 |
| 2:T:90:SER:HB2 | 2:T:150:ASP:H | 1.70 | 0.56 |
| 2:U:123:PRO:HG3 | 2:U:151:PHE:HD1 | 1.68 | 0.56 |
| 6:Q:31:SER:HB3 | 5:V:504:ILE:HG23 | 1.87 | 0.56 |
| 7:I:51:ILE:HG12 | 7:I:55:GLY:HA2 | 1.86 | 0.56 |
| 8:J:47:LEU:HA | 8:J:58:VAL:HG21 | 1.86 | 0.56 |
| 1:C:103:LEU:HB2 | 4:D:97:TYR:HB2 | 1.87 | 0.56 |
| 8:H:24:ARG:NH2 | 8:H:70:ASP:OD1 | 2.38 | 0.56 |
| 5:X:596:ARG:HB2 | 5:X:596:ARG:HH11 | 1.70 | 0.56 |
| 1:C:86:ARG:HH22 | 4:B:78:THR:HG21 | 1.69 | 0.56 |
| 2:S:115:LYS:HE2 | 2:S:147:CYS:HB2 | 1.88 | 0.56 |
| 2:S:218:ILE:HG23 | 2:S:234:PHE:CE1 | 2.41 | 0.56 |
| 6:Q:30:SER:HG | 5:V:504:ILE:N | 2.03 | 0.56 |
| 8:H:21:ILE:HG12 | 8:H:103:THR:HG21 | 1.88 | 0.56 |
| 8:H:94:THR:O | 8:H:97:ILE:HG23 | 2.06 | 0.56 |
| 1:A:72:GLU:OE2 | 1:A:79:TYR:HE2 | 1.88 | 0.56 |
| 1:A:72:GLU:O | 1:A:72:GLU:HG2 | 2.05 | 0.56 |
| 2:U:57:LEU:HD11 | 2:U:63:LEU:HG | 1.87 | 0.56 |
| 7:G:68:PHE:CD2 | 7:G:83:MET:HA | 2.41 | 0.56 |

*Continued on next page...*

For Manuscript Review

Continued from previous page...

| Atom-1 | Atom-2 | Interatomic distance (Å) | Clash overlap (Å) |
|---|---|---|---|
| 2:S:234:PHE:N | 2:S:242:VAL:O | 2.37 | 0.56 |
| 6:Q:91:THR:HB | 6:Q:116:VAL:HG12 | 1.88 | 0.56 |
| 7:G:35:HIS:HE2 | 7:G:100:TRP:HA | 1.71 | 0.56 |
| 1:C:31:SER:HB3 | 1:C:100:PHE:HE1 | 1.70 | 0.56 |
| 2:S:182:ALA:HB2 | 5:V:562:ALA:HB2 | 1.87 | 0.55 |
| 6:O:22:CYS:HB3 | 6:O:79:LEU:HD11 | 1.86 | 0.55 |
| 1:A:106:ASP:OD2 | 1:A:107:TYR:N | 2.39 | 0.55 |
| 2:U:117:ASP:O | 4:B:102:LEU:HD13 | 2.05 | 0.55 |
| 7:G:91:THR:HG23 | 7:G:113:THR:HA | 1.88 | 0.55 |
| 2:U:47:ASP:HB3 | 2:U:50:LYS:HB3 | 1.89 | 0.55 |
| 7:G:9:GLY:HA2 | 7:G:112:VAL:HG22 | 1.87 | 0.55 |
| 3:N:20:THR:HG22 | 3:N:74:THR:HG22 | 1.89 | 0.55 |
| 8:H:101:GLN:NE2 | 8:H:102:GLY:O | 2.40 | 0.55 |
| 7:I:16:ARG:HH22 | 7:I:84:ASN:HA | 1.70 | 0.55 |
| 7:I:6:GLU:HG3 | 7:I:22:CYS:HA | 1.89 | 0.55 |
| 2:S:256:LEU:HD21 | 2:S:277:VAL:HG21 | 1.88 | 0.55 |
| 3:R:17:ASP:O | 3:R:78:LEU:HD23 | 2.07 | 0.55 |
| 8:H:11:LEU:HB3 | 8:H:19:ILE:HD11 | 1.89 | 0.55 |
| 7:I:8:GLY:O | 7:I:18:LEU:HD12 | 2.07 | 0.55 |
| 1:A:31:SER:HB3 | 1:A:100:PHE:HE1 | 1.71 | 0.55 |
| 6:O:29:SER:O | 6:O:72:ARG:NH2 | 2.40 | 0.55 |
| 4:F:60:ARG:NH2 | 4:F:64:VAL:O | 2.40 | 0.54 |
| 7:I:61:ALA:HB3 | 7:I:64:VAL:HG22 | 1.89 | 0.54 |
| 4:F:45:LYS:HD3 | 4:F:90:ALA:HB2 | 1.89 | 0.54 |
| 6:Q:6:GLU:HG2 | 6:Q:112:THR:HG23 | 1.88 | 0.54 |
| 8:J:22:THR:N | 8:J:71:PHE:O | 2.40 | 0.54 |
| 7:G:86:LEU:HD12 | 7:G:114:VAL:HG22 | 1.88 | 0.54 |
| 2:U:52:VAL:HG22 | 2:U:54:ARG:H | 1.72 | 0.54 |
| 5:W:521:GLN:HG2 | 5:W:544:THR:CG2 | 2.37 | 0.54 |
| 6:M:2:VAL:HG13 | 6:M:3:GLN:HG2 | 1.89 | 0.54 |
| 6:M:35:ASN:HD21 | 6:M:103:HIS:HE1 | 1.54 | 0.54 |
| 8:H:33:LEU:HD21 | 8:H:88:CYS:HB2 | 1.90 | 0.54 |
| 8:H:95:PRO:HB2 | 8:H:96:PRO:HD3 | 1.88 | 0.54 |
| 2:S:154:HIS:CE1 | 2:S:156:GLU:HB2 | 2.43 | 0.54 |
| 2:U:164:ARG:NH2 | 5:W:575:ALA:O | 2.41 | 0.54 |
| 3:P:35:TRP:CE2 | 3:P:73:LEU:HB2 | 2.43 | 0.54 |
| 1:C:35:HIS:HD2 | 1:C:47:TRP:HE1 | 1.56 | 0.54 |
| 6:O:36:TRP:NE1 | 6:O:81:LEU:HB2 | 2.23 | 0.54 |
| 5:V:611:GLU:OE2 | 5:X:602:HIS:ND1 | 2.41 | 0.54 |
| 3:N:37:GLN:HB2 | 3:N:47:LEU:HD11 | 1.90 | 0.54 |
| 7:I:36:TRP:NE1 | 7:I:79:LEU:HD21 | 2.21 | 0.54 |

Continued on next page...

For Manuscript Review

*Continued from previous page...*

| Atom-1 | Atom-2 | Interatomic distance (Å) | Clash overlap (Å) |
|---|---|---|---|
| 8:J:33:LEU:HB3 | 8:J:51:ALA:HB2 | 1.90 | 0.54 |
| 8:J:89:GLN:OE1 | 8:J:99:PHE:HA | 2.07 | 0.53 |
| 4:D:39:LEU:HD13 | 4:D:77:PHE:CD2 | 2.44 | 0.53 |
| 3:N:51:ALA:HB1 | 3:N:66:GLY:HA3 | 1.89 | 0.53 |
| 8:J:29:ILE:HG23 | 8:J:92:PHE:HB2 | 1.90 | 0.53 |
| 2:U:120:GLU:OE2 | 2:U:172:ARG:HD3 | 2.08 | 0.53 |
| 6:Q:6:GLU:OE1 | 6:Q:6:GLU:N | 2.39 | 0.53 |
| 6:M:48:VAL:HG23 | 6:M:64:VAL:HG21 | 1.90 | 0.53 |
| 2:T:49:ASP:OD1 | 2:T:50:LYS:N | 2.41 | 0.53 |
| 2:S:162:TYR:CE2 | 2:S:176:PHE:HB3 | 2.44 | 0.53 |
| 7:G:34:MET:SD | 7:G:98:ARG:HA | 2.48 | 0.53 |
| 8:H:17:ASP:OD1 | 8:H:18:ARG:N | 2.41 | 0.53 |
| 3:P:2:ILE:HD13 | 3:P:93:SER:HB2 | 1.90 | 0.53 |
| 1:C:38:ARG:NH1 | 1:C:93:TYR:OH | 2.40 | 0.53 |
| 5:W:521:GLN:HG3 | 5:W:542:ILE:O | 2.08 | 0.53 |
| 6:M:101:TYR:HD2 | 6:M:102:PRO:HD3 | 1.73 | 0.53 |
| 7:G:64:VAL:HB | 7:G:68:PHE:HD1 | 1.74 | 0.53 |
| 5:V:521:GLN:HA | 5:V:544:THR:OG1 | 2.09 | 0.53 |
| 4:D:53:LEU:HA | 4:D:64:VAL:HG21 | 1.90 | 0.53 |
| 4:F:96:GLN:HE21 | 4:F:103:THR:H | 1.56 | 0.53 |
| 5:V:583:SER:O | 5:V:587:ARG:HG3 | 2.09 | 0.53 |
| 2:U:38:ILE:HB | 2:U:187:PRO:HD3 | 1.90 | 0.53 |
| 2:S:57:LEU:HB3 | 5:X:594:LEU:HD22 | 1.90 | 0.53 |
| 2:U:78:ASP:OD1 | 2:U:81:SER:HB3 | 2.08 | 0.53 |
| 5:W:587:ARG:HA | 5:W:590:ILE:HG12 | 1.91 | 0.53 |
| 8:J:37:GLN:HB3 | 8:J:47:LEU:HD11 | 1.90 | 0.53 |
| 1:C:68:THR:HG23 | 1:C:81:GLN:HB3 | 1.90 | 0.53 |
| 3:P:90:GLN:HE21 | 3:P:96:THR:HB | 1.74 | 0.52 |
| 1:A:101:GLY:O | 2:U:143:GLY:HA2 | 2.09 | 0.52 |
| 2:U:191:LYS:HB3 | 2:U:195:SER:HB3 | 1.92 | 0.52 |
| 6:M:12:VAL:O | 6:M:116:VAL:HA | 2.09 | 0.52 |
| 6:O:67:ARG:NH2 | 6:O:90:ASP:OD1 | 2.37 | 0.52 |
| 5:V:521:GLN:NE2 | 5:X:574:ARG:HD2 | 2.24 | 0.52 |
| 8:J:36:TYR:HE2 | 8:J:89:GLN:HB2 | 1.74 | 0.52 |
| 3:R:6:GLN:OE1 | 3:R:101:THR:HG23 | 2.09 | 0.52 |
| 7:G:104:ASP:OD1 | 7:G:105:TYR:N | 2.43 | 0.52 |
| 2:T:196:SER:O | 2:T:198:PRO:HD3 | 2.09 | 0.52 |
| 8:J:39:LYS:HD3 | 8:J:40:PRO:HD2 | 1.91 | 0.52 |
| 2:U:263:SER:HB3 | 8:J:32:TYR:CE1 | 2.45 | 0.52 |
| 6:Q:68:PHE:CE1 | 6:Q:83:MET:HB2 | 2.45 | 0.52 |
| 2:T:98:ASN:ND2 | 5:W:582:PHE:HD1 | 2.08 | 0.52 |

*Continued on next page...*

For Manuscript Review

Continued from previous page...

| Atom-1 | Atom-2 | Interatomic distance (Å) | Clash overlap (Å) |
|---|---|---|---|
| 2:T:249:THR:HG21 | 2:T:286:GLY:HA3 | 1.90 | 0.52 |
| 8:H:34:HIS:HE1 | 8:H:91:SER:HB2 | 1.74 | 0.52 |
| 1:A:29:PHE:O | 1:A:71:ARG:NH2 | 2.43 | 0.52 |
| 2:T:116:PRO:HD2 | 1:C:58:TYR:CE2 | 2.44 | 0.52 |
| 2:U:45:VAL:HG21 | 6:O:56:GLY:HA3 | 1.90 | 0.52 |
| 3:N:11:LEU:HD11 | 3:N:19:VAL:HG21 | 1.92 | 0.52 |
| 3:N:34:ASN:OD1 | 3:N:89:GLN:HB3 | 2.10 | 0.52 |
| 2:T:291:TRP:CD1 | 5:W:513:PRO:HD2 | 2.44 | 0.52 |
| 4:F:53:LEU:HA | 4:F:64:VAL:HG11 | 1.92 | 0.52 |
| 1:E:86:ARG:HH22 | 4:D:78:THR:HG21 | 1.74 | 0.52 |
| 2:S:63:LEU:HB3 | 5:V:585:LEU:HD22 | 1.92 | 0.52 |
| 4:B:20:THR:HG22 | 4:B:80:THR:HG22 | 1.92 | 0.52 |
| 2:S:104:TRP:CZ2 | 2:S:134:ARG:HD2 | 2.45 | 0.52 |
| 7:G:38:ARG:HD3 | 7:G:94:TYR:CZ | 2.44 | 0.52 |
| 3:P:63:SER:OG | 3:P:74:THR:HG23 | 2.10 | 0.52 |
| 8:J:49:TYR:HB2 | 8:J:53:THR:HG23 | 1.91 | 0.52 |
| 2:T:154:HIS:CE1 | 2:T:156:GLU:HB2 | 2.45 | 0.51 |
| 6:O:97:ALA:HB1 | 6:O:105:PHE:HB3 | 1.92 | 0.51 |
| 6:O:101:TYR:CD1 | 9:Y:2:NAG:H4 | 2.45 | 0.51 |
| 8:J:83:PHE:CG | 8:J:107:ILE:HG12 | 2.45 | 0.51 |
| 2:T:154:HIS:ND1 | 2:T:156:GLU:HB2 | 2.25 | 0.51 |
| 6:M:35:ASN:N | 6:M:97:ALA:O | 2.35 | 0.51 |
| 8:H:61:ARG:HD2 | 8:H:77:SER:O | 2.11 | 0.51 |
| 2:T:51:LEU:HB3 | 5:W:595:GLN:HE21 | 1.74 | 0.51 |
| 8:H:22:THR:HG23 | 8:H:72:THR:HG22 | 1.92 | 0.51 |
| 7:I:14:PRO:HA | 7:I:86:LEU:HD22 | 1.93 | 0.51 |
| 1:C:72:GLU:OE2 | 1:C:74:ALA:HB3 | 2.10 | 0.51 |
| 2:U:156:GLU:HG2 | 9:Y:2:NAG:H62 | 1.93 | 0.51 |
| 1:E:6:GLU:OE2 | 1:E:95:CYS:N | 2.28 | 0.51 |
| 3:R:29:ILE:N | 3:R:68:GLY:O | 2.41 | 0.51 |
| 6:M:4:LEU:HD23 | 6:M:24:ALA:HA | 1.92 | 0.51 |
| 8:J:79:GLN:HB3 | 8:J:80:PRO:HD2 | 1.93 | 0.51 |
| 2:U:98:ASN:HB2 | 5:X:582:PHE:CD1 | 2.46 | 0.51 |
| 3:P:7:SER:HB3 | 3:P:8:PRO:HD3 | 1.92 | 0.51 |
| 8:J:80:PRO:HA | 8:J:83:PHE:CD2 | 2.45 | 0.51 |
| 4:D:38:TYR:HB2 | 4:D:98:TYR:HB2 | 1.92 | 0.51 |
| 5:W:550:ASN:OD1 | 5:W:551:GLN:N | 2.44 | 0.51 |
| 8:J:85:THR:HB | 8:J:102:GLY:HA3 | 1.91 | 0.51 |
| 4:F:67:ARG:NH2 | 4:F:88:ASP:OD2 | 2.41 | 0.51 |
| 7:G:39:GLN:O | 7:G:92:ALA:HB1 | 2.11 | 0.51 |
| 7:G:98:ARG:HD2 | 7:G:98:ARG:O | 2.11 | 0.51 |

Continued on next page...

For Manuscript Review

*Continued from previous page...*

| Atom-1 | Atom-2 | Interatomic distance (Å) | Clash overlap (Å) |
|---|---|---|---|
| 1:A:36:TRP:NE1 | 1:A:80:LEU:HB2 | 2.26 | 0.51 |
| 5:V:521:GLN:HE21 | 5:X:574:ARG:HD2 | 1.74 | 0.51 |
| 2:S:196:SER:O | 2:S:198:PRO:HD3 | 2.11 | 0.51 |
| 3:P:92:TYR:O | 5:V:528:GLY:HA3 | 2.10 | 0.51 |
| 6:O:102:PRO:HA | 5:V:529:LEU:HD11 | 1.93 | 0.51 |
| 4:D:3:VAL:HG12 | 4:D:26:SER:HB3 | 1.93 | 0.51 |
| 2:S:234:PHE:HB3 | 2:S:242:VAL:HG23 | 1.93 | 0.50 |
| 7:I:64:VAL:HA | 7:I:67:ARG:NH1 | 2.26 | 0.50 |
| 1:A:58:TYR:HB3 | 2:U:117:ASP:OD1 | 2.11 | 0.50 |
| 2:S:114:LYS:O | 2:S:144:THR:HA | 2.11 | 0.50 |
| 8:J:34:HIS:N | 8:J:89:GLN:O | 2.40 | 0.50 |
| 2:S:37:VAL:HG23 | 2:S:46:SER:HB2 | 1.93 | 0.50 |
| 5:W:521:GLN:HG2 | 5:W:544:THR:HG23 | 1.93 | 0.50 |
| 8:H:55:GLN:HB3 | 8:H:58:VAL:HG13 | 1.93 | 0.50 |
| 6:O:59:TYR:HD2 | 5:V:527:ILE:CD1 | 2.24 | 0.50 |
| 8:J:37:GLN:HE21 | 8:J:84:ALA:HB3 | 1.76 | 0.50 |
| 1:E:30:SER:O | 1:E:53:THR:HG23 | 2.10 | 0.50 |
| 1:E:38:ARG:HG3 | 1:E:91:ALA:HB3 | 1.93 | 0.50 |
| 1:A:29:PHE:HZ | 1:A:78:LEU:HB2 | 1.75 | 0.50 |
| 2:S:247:ARG:HH11 | 2:S:279:PRO:HD2 | 1.76 | 0.50 |
| 3:N:61:ARG:NH2 | 3:N:82:ASP:OD2 | 2.32 | 0.50 |
| 7:G:68:PHE:HD2 | 7:G:83:MET:HA | 1.76 | 0.50 |
| 8:H:13:ALA:O | 8:H:108:LYS:N | 2.43 | 0.50 |
| 6:O:35:ASN:ND2 | 6:O:105:PHE:HE1 | 2.06 | 0.50 |
| 2:S:156:GLU:HG2 | 9:L:2:NAG:H62 | 1.94 | 0.50 |
| 3:R:81:GLU:OE2 | 3:R:81:GLU:N | 2.36 | 0.50 |
| 6:O:8:GLY:O | 6:O:18:LEU:HD21 | 2.11 | 0.50 |
| 5:W:517:TYR:CZ | 5:W:546:GLY:HA3 | 2.47 | 0.50 |
| 1:A:92:VAL:HG22 | 1:A:113:LEU:HD13 | 1.94 | 0.50 |
| 2:U:281:ILE:HG21 | 2:U:301:ILE:HD12 | 1.93 | 0.50 |
| 6:O:32:TYR:HD1 | 6:O:98:LYS:HD3 | 1.76 | 0.50 |
| 2:T:94:PRO:HB3 | 2:T:169:VAL:HG21 | 1.94 | 0.50 |
| 2:S:96:VAL:HG22 | 2:S:167:SER:HB2 | 1.93 | 0.50 |
| 2:U:104:TRP:NE1 | 5:X:545:GLU:OE1 | 2.41 | 0.50 |
| 7:I:100:TRP:HD1 | 7:I:101:ASN:OD1 | 1.95 | 0.50 |
| 2:S:232:TYR:HB2 | 2:S:244:LEU:HB2 | 1.92 | 0.49 |
| 3:N:50:ALA:HB2 | 6:M:102:PRO:HG3 | 1.94 | 0.49 |
| 3:N:95:LEU:HD21 | 6:M:103:HIS:ND1 | 2.28 | 0.49 |
| 8:J:95:PRO:CD | 8:J:96:PRO:HD3 | 2.42 | 0.49 |
| 7:G:4:LEU:HD21 | 7:G:27:PHE:HE2 | 1.78 | 0.49 |
| 8:H:30:SER:H | 8:H:92:PHE:HE2 | 1.60 | 0.49 |

*Continued on next page...*

For Manuscript Review

Continued from previous page...

| Atom-1 | Atom-2 | Interatomic distance (Å) | Clash overlap (Å) |
|---|---|---|---|
| 7:I:37:VAL:HB | 7:I:95:TYR:HB2 | 1.94 | 0.49 |
| 6:M:76:LYS:HB2 | 6:M:78:THR:HG22 | 1.94 | 0.49 |
| 7:G:67:ARG:HB3 | 7:G:85:SER:HB3 | 1.94 | 0.49 |
| 8:H:6:GLN:HA | 8:H:23:CYS:HA | 1.94 | 0.49 |
| 7:I:37:VAL:HG11 | 7:I:45:LEU:HD22 | 1.93 | 0.49 |
| 4:B:43:GLN:HB2 | 4:B:53:LEU:HD11 | 1.93 | 0.49 |
| 2:T:77:THR:OG1 | 2:T:107:ASN:ND2 | 2.45 | 0.49 |
| 2:T:311:VAL:O | 2:T:463:ASN:HB3 | 2.12 | 0.49 |
| 2:U:255:GLN:OE1 | 2:U:301:ILE:HB | 2.12 | 0.49 |
| 6:Q:79:LEU:HD23 | 6:Q:79:LEU:H | 1.77 | 0.49 |
| 4:F:38:TYR:HD1 | 4:F:97:TYR:CE2 | 2.29 | 0.49 |
| 6:M:101:TYR:CZ | 9:K:3:BMA:H3 | 2.48 | 0.49 |
| 1:C:39:GLN:HB2 | 1:C:45:LEU:HD13 | 1.94 | 0.49 |
| 2:T:258:GLU:HA | 10:T:503:NAG:H82 | 1.94 | 0.49 |
| 6:M:83:MET:HE1 | 6:M:114:VAL:HG21 | 1.95 | 0.49 |
| 3:R:38:GLN:NE2 | 6:Q:39:GLN:HE22 | 2.11 | 0.49 |
| 2:T:164:ARG:NH1 | 5:V:575:ALA:O | 2.45 | 0.49 |
| 3:R:94:THR:HG23 | 3:R:96:THR:HG23 | 1.95 | 0.49 |
| 4:F:2:ILE:HG13 | 4:F:27:GLN:HB2 | 1.93 | 0.49 |
| 4:D:43:GLN:HB2 | 4:D:53:LEU:HD11 | 1.94 | 0.49 |
| 2:T:146:PRO:HG3 | 1:C:56:ASP:OD2 | 2.13 | 0.49 |
| 2:S:88:PHE:HB3 | 2:S:149:GLY:H | 1.77 | 0.49 |
| 2:U:86:TRP:HZ3 | 2:U:111:LEU:HD21 | 1.76 | 0.49 |
| 5:W:527:ILE:HD13 | 6:Q:59:TYR:CD1 | 2.40 | 0.49 |
| 7:G:100:TRP:HE1 | 8:H:94:THR:HG21 | 1.77 | 0.49 |
| 7:I:39:GLN:O | 7:I:92:ALA:HB1 | 2.13 | 0.49 |
| 2:T:193:PHE:HE1 | 5:W:554:LEU:HD21 | 1.77 | 0.49 |
| 6:Q:105:PHE:O | 6:Q:108:TRP:NE1 | 2.45 | 0.49 |
| 8:J:78:LEU:HD23 | 8:J:83:PHE:HE1 | 1.77 | 0.49 |
| 1:E:31:SER:HB3 | 1:E:100:PHE:HE1 | 1.78 | 0.49 |
| 5:W:507:ALA:H | 6:M:31:SER:HB2 | 1.77 | 0.48 |
| 6:Q:99:ARG:NH1 | 6:Q:103:HIS:HD2 | 2.12 | 0.48 |
| 2:T:240:THR:OG1 | 2:T:266:ARG:HB2 | 2.13 | 0.48 |
| 5:W:507:ALA:HB2 | 6:M:31:SER:HB2 | 1.94 | 0.48 |
| 7:G:38:ARG:NH1 | 7:G:94:TYR:OH | 2.46 | 0.48 |
| 1:A:102:GLU:HG2 | 2:U:221:GLN:HE22 | 1.78 | 0.48 |
| 2:T:163:ASP:OD2 | 5:W:543:TYR:OH | 2.31 | 0.48 |
| 1:E:106:ASP:OD1 | 1:E:107:TYR:N | 2.46 | 0.48 |
| 2:T:68:LEU:HD11 | 5:W:558:LEU:HG | 1.96 | 0.48 |
| 2:S:197:HIS:O | 2:S:197:HIS:ND1 | 2.47 | 0.48 |
| 2:T:252:PHE:HD2 | 2:T:301:ILE:HD11 | 1.79 | 0.48 |

Continued on next page...

For Manuscript Review

For Manuscript Review

*Continued from previous page...*

| Atom-1 | Atom-2 | Interatomic distance (Å) | Clash overlap (Å) |
|---|---|---|---|
| 4:F:24:LYS:HG3 | 4:F:76:GLU:HB3 | 1.96 | 0.48 |
| 3:P:38:GLN:OE1 | 6:O:39:GLN:NE2 | 2.46 | 0.48 |
| 5:V:562:ALA:HA | 5:V:565:THR:HG22 | 1.96 | 0.48 |
| 7:I:94:TYR:O | 7:I:109:GLY:HA2 | 2.13 | 0.48 |
| 1:E:51:ILE:HG21 | 1:E:78:LEU:HD11 | 1.94 | 0.48 |
| 7:G:6:GLU:HB3 | 7:G:110:THR:HB | 1.95 | 0.48 |
| 6:O:40:ALA:HB3 | 6:O:43:LYS:HB2 | 1.95 | 0.48 |
| 7:I:68:PHE:CE1 | 7:I:83:MET:HB2 | 2.49 | 0.48 |
| 8:J:36:TYR:O | 8:J:87:TYR:N | 2.41 | 0.48 |
| 1:E:32:TYR:CD1 | 1:E:97:ARG:HD3 | 2.48 | 0.48 |
| 3:R:45:LYS:NZ | 3:R:58:VAL:HG22 | 2.29 | 0.48 |
| 7:G:36:TRP:NE1 | 7:G:81:LEU:HB2 | 2.29 | 0.48 |
| 2:T:252:PHE:CE2 | 2:T:279:PRO:HG3 | 2.49 | 0.48 |
| 3:R:93:SER:O | 3:R:94:THR:HG22 | 2.14 | 0.48 |
| 6:O:91:THR:HG22 | 6:O:116:VAL:H | 1.79 | 0.48 |
| 2:S:234:PHE:CE2 | 2:S:236:VAL:HG12 | 2.49 | 0.47 |
| 3:P:95:LEU:HD21 | 6:O:103:HIS:CD2 | 2.49 | 0.47 |
| 5:X:603:ILE:H | 5:X:603:ILE:HD12 | 1.79 | 0.47 |
| 7:I:16:ARG:HE | 7:I:17:SER:N | 2.00 | 0.47 |
| 7:I:67:ARG:HB2 | 7:I:85:SER:H | 1.80 | 0.47 |
| 4:B:89:VAL:HG23 | 4:B:110:VAL:O | 2.14 | 0.47 |
| 2:T:79:VAL:CG1 | 2:T:80:PRO:HD3 | 2.43 | 0.47 |
| 7:I:16:ARG:NH2 | 7:I:84:ASN:HA | 2.29 | 0.47 |
| 7:I:38:ARG:O | 7:I:46:GLU:HG2 | 2.14 | 0.47 |
| 6:Q:69:THR:HG23 | 6:Q:82:GLN:HB3 | 1.97 | 0.47 |
| 2:T:114:LYS:O | 2:T:144:THR:HA | 2.15 | 0.47 |
| 2:S:191:LYS:O | 2:S:195:SER:HB2 | 2.15 | 0.47 |
| 2:U:132:PHE:HD2 | 2:U:135:CYS:SG | 2.37 | 0.47 |
| 4:F:31:TYR:HB2 | 4:F:38:TYR:HE2 | 1.78 | 0.47 |
| 5:W:562:ALA:HA | 5:W:565:THR:HG22 | 1.95 | 0.47 |
| 7:G:17:SER:HA | 7:G:83:MET:O | 2.14 | 0.47 |
| 5:X:584:ILE:H | 5:X:584:ILE:HD12 | 1.80 | 0.47 |
| 4:B:61:GLU:HB3 | 4:B:64:VAL:HG23 | 1.95 | 0.47 |
| 7:I:19:ARG:HG2 | 7:I:19:ARG:HH11 | 1.79 | 0.47 |
| 2:T:242:VAL:HG11 | 2:T:256:LEU:HD21 | 1.97 | 0.47 |
| 2:U:154:HIS:CE1 | 2:U:156:GLU:HB2 | 2.49 | 0.47 |
| 2:U:282:ASP:OD1 | 2:U:283:THR:HG23 | 2.15 | 0.47 |
| 6:Q:72:ARG:HA | 6:Q:79:LEU:HA | 1.96 | 0.47 |
| 8:H:7:SER:HB2 | 8:H:8:PRO:HD3 | 1.95 | 0.47 |
| 6:O:32:TYR:CE2 | 5:X:507:ALA:HA | 2.50 | 0.47 |
| 2:T:131:GLY:O | 2:T:163:ASP:HB2 | 2.14 | 0.47 |

*Continued on next page...*

Continued from previous page...

| Atom-1 | Atom-2 | Interatomic distance (Å) | Clash overlap (Å) |
|---|---|---|---|
| 4:F:61:GLU:O | 4:F:64:VAL:HG22 | 2.14 | 0.47 |
| 6:M:98:LYS:HB3 | 6:M:107:ILE:HG22 | 1.95 | 0.47 |
| 7:I:76:LYS:NZ | 7:I:78:THR:HG21 | 2.30 | 0.47 |
| 4:B:24:LYS:HZ3 | 4:B:76:GLU:HB3 | 1.79 | 0.47 |
| 2:U:73:ASN:HB3 | 5:X:508:GLN:HE21 | 1.80 | 0.47 |
| 4:F:97:TYR:HA | 4:F:102:LEU:HD22 | 1.96 | 0.47 |
| 8:H:47:LEU:HA | 8:H:58:VAL:HG11 | 1.96 | 0.47 |
| 6:O:47:TRP:CZ3 | 5:V:527:ILE:HG13 | 2.50 | 0.47 |
| 1:C:29:PHE:O | 1:C:71:ARG:NH2 | 2.47 | 0.47 |
| 2:S:123:PRO:HG3 | 2:S:151:PHE:CD2 | 2.50 | 0.47 |
| 2:U:249:THR:HG23 | 2:U:252:PHE:H | 1.79 | 0.47 |
| 2:U:291:TRP:HD1 | 2:U:295:LYS:HZ1 | 1.63 | 0.47 |
| 4:F:43:GLN:HB2 | 4:F:53:LEU:HD11 | 1.96 | 0.47 |
| 5:W:507:ALA:CB | 6:M:31:SER:HB2 | 2.45 | 0.47 |
| 1:A:96:ALA:HB1 | 1:A:105:PHE:HB3 | 1.96 | 0.47 |
| 1:A:103:LEU:HD23 | 1:A:103:LEU:HA | 1.74 | 0.47 |
| 2:U:63:LEU:HB3 | 5:X:585:LEU:HD22 | 1.97 | 0.47 |
| 2:U:304:GLU:OE2 | 8:J:32:TYR:HA | 2.15 | 0.47 |
| 8:H:66:GLY:HA3 | 8:H:71:PHE:HA | 1.96 | 0.47 |
| 7:I:18:LEU:H | 7:I:18:LEU:HD23 | 1.78 | 0.47 |
| 6:M:60:TYR:OH | 6:M:70:ILE:N | 2.36 | 0.46 |
| 6:M:88:ALA:HA | 6:M:116:VAL:HG21 | 1.97 | 0.46 |
| 8:J:35:TRP:CZ3 | 8:J:88:CYS:HB3 | 2.51 | 0.46 |
| 7:G:62:ASP:O | 7:G:65:LYS:HG2 | 2.14 | 0.46 |
| 8:J:8:PRO:HB2 | 8:J:103:THR:HB | 1.97 | 0.46 |
| 2:T:292:GLU:HG3 | 5:W:510:LYS:NZ | 2.31 | 0.46 |
| 8:H:61:ARG:HG3 | 8:H:62:PHE:HD1 | 1.81 | 0.46 |
| 5:V:530:ALA:HA | 5:V:535:PHE:O | 2.16 | 0.46 |
| 8:J:33:LEU:HD21 | 8:J:88:CYS:HB2 | 1.97 | 0.46 |
| 5:W:563:ASN:O | 5:W:566:THR:HG22 | 2.15 | 0.46 |
| 5:W:607:ASP:OD1 | 5:W:608:CYS:N | 2.48 | 0.46 |
| 6:M:3:GLN:HB2 | 6:M:25:SER:HB2 | 1.97 | 0.46 |
| 8:H:5:THR:HB | 8:H:24:ARG:HB3 | 1.96 | 0.46 |
| 7:I:70:ILE:HD11 | 7:I:79:LEU:HD21 | 1.97 | 0.46 |
| 8:J:6:GLN:NE2 | 8:J:86:TYR:O | 2.49 | 0.46 |
| 1:E:36:TRP:HE1 | 1:E:78:LEU:CD2 | 2.29 | 0.46 |
| 2:T:51:LEU:HD12 | 2:T:51:LEU:O | 2.16 | 0.46 |
| 2:T:85:ARG:HH22 | 2:T:178:GLU:CD | 2.19 | 0.46 |
| 2:T:98:ASN:HD22 | 5:W:582:PHE:HD1 | 1.64 | 0.46 |
| 2:T:294:LYS:HG3 | 2:T:296:ASN:ND2 | 2.30 | 0.46 |
| 2:U:234:PHE:HB3 | 2:U:242:VAL:HG23 | 1.98 | 0.46 |

Continued on next page...

For Manuscript Review

Continued from previous page...

| Atom-1 | Atom-2 | Interatomic distance (Å) | Clash overlap (Å) |
|---|---|---|---|
| 3:N:34:ASN:HD22 | 6:M:104:SER:HB2 | 1.80 | 0.46 |
| 4:F:19:ALA:HB2 | 4:F:84:LEU:HD12 | 1.97 | 0.46 |
| 4:F:61:GLU:HB3 | 4:F:64:VAL:HG13 | 1.98 | 0.46 |
| 8:H:47:LEU:HD13 | 8:H:62:PHE:HD2 | 1.81 | 0.46 |
| 8:J:5:THR:OG1 | 8:J:24:ARG:NE | 2.49 | 0.46 |
| 8:J:38:GLN:HB2 | 8:J:44:PRO:HG3 | 1.98 | 0.46 |
| 4:B:7:SER:HB2 | 4:B:22:ASN:HB2 | 1.97 | 0.46 |
| 2:S:142:SER:O | 2:S:221:GLN:HA | 2.16 | 0.46 |
| 3:N:17:ASP:O | 3:N:78:LEU:HD23 | 2.15 | 0.46 |
| 5:W:514:ASN:OD1 | 5:W:549:HIS:HB2 | 2.15 | 0.46 |
| 3:P:46:LEU:HB2 | 6:O:105:PHE:O | 2.16 | 0.46 |
| 6:O:4:LEU:HD22 | 6:O:22:CYS:SG | 2.56 | 0.46 |
| 8:J:61:ARG:NH1 | 8:J:81:GLU:OE2 | 2.40 | 0.46 |
| 4:D:61:GLU:HB3 | 4:D:64:VAL:HG23 | 1.97 | 0.46 |
| 2:S:49:ASP:OD1 | 2:S:50:LYS:N | 2.49 | 0.46 |
| 2:U:252:PHE:CZ | 2:U:279:PRO:HG3 | 2.51 | 0.46 |
| 3:N:49:TYR:CE2 | 3:N:55:GLN:HA | 2.51 | 0.46 |
| 6:Q:29:SER:O | 6:Q:72:ARG:NH1 | 2.47 | 0.46 |
| 2:T:156:GLU:OE2 | 9:K:2:NAG:H62 | 2.17 | 0.45 |
| 8:J:37:GLN:CB | 8:J:47:LEU:HD11 | 2.45 | 0.45 |
| 4:B:3:VAL:HG22 | 4:B:26:SER:HB3 | 1.98 | 0.45 |
| 2:S:71:GLU:OE2 | 2:S:106:GLU:N | 2.39 | 0.45 |
| 2:S:77:THR:O | 2:S:109:TYR:OH | 2.20 | 0.45 |
| 4:F:89:VAL:O | 4:F:89:VAL:HG23 | 2.17 | 0.45 |
| 2:U:114:LYS:O | 2:U:144:THR:HA | 2.16 | 0.45 |
| 2:U:162:TYR:CE1 | 2:U:176:PHE:HB3 | 2.50 | 0.45 |
| 6:Q:71:SER:O | 6:Q:80:TYR:N | 2.42 | 0.45 |
| 8:H:29:ILE:HB | 8:H:71:PHE:HE2 | 1.81 | 0.45 |
| 1:C:87:ALA:O | 1:C:90:THR:HG22 | 2.16 | 0.45 |
| 2:U:254:LEU:O | 2:U:257:ASN:HB3 | 2.16 | 0.45 |
| 6:Q:99:ARG:HH11 | 6:Q:103:HIS:HD2 | 1.63 | 0.45 |
| 7:I:22:CYS:HB3 | 7:I:79:LEU:HD22 | 1.97 | 0.45 |
| 5:W:527:ILE:HD11 | 6:Q:103:HIS:NE2 | 2.31 | 0.45 |
| 1:A:12:VAL:HG21 | 1:A:85:LEU:HD12 | 1.99 | 0.45 |
| 5:W:505:VAL:HG12 | 5:W:505:VAL:O | 2.16 | 0.45 |
| 6:M:17:SER:HB3 | 6:M:82:GLN:NE2 | 2.32 | 0.45 |
| 6:M:51:ILE:HD13 | 6:M:79:LEU:HD11 | 1.99 | 0.45 |
| 1:C:66:ARG:NH1 | 1:C:89:ASP:OD2 | 2.41 | 0.45 |
| 1:A:63:VAL:HB | 1:A:67:PHE:CG | 2.51 | 0.45 |
| 1:A:99:TRP:CZ2 | 2:U:272:LYS:HD2 | 2.52 | 0.45 |
| 2:U:89:ARG:NH1 | 5:V:536:GLY:O | 2.49 | 0.45 |

Continued on next page...

For Manuscript Review

Continued from previous page...

| Atom-1 | Atom-2 | Interatomic distance (Å) | Clash overlap (Å) |
|---|---|---|---|
| 3:N:83:PHE:O | 3:N:103:LEU:HB2 | 2.16 | 0.45 |
| 6:M:38:ARG:HD3 | 6:M:94:TYR:CZ | 2.51 | 0.45 |
| 6:O:36:TRP:CD1 | 6:O:81:LEU:HB2 | 2.51 | 0.45 |
| 5:V:513:PRO:HA | 5:V:552:ASP:OD2 | 2.16 | 0.45 |
| 2:T:245:GLU:OE1 | 2:T:247:ARG:NH1 | 2.50 | 0.45 |
| 2:U:255:GLN:O | 2:U:259:THR:HG23 | 2.17 | 0.45 |
| 3:R:2:ILE:HD11 | 3:R:27:GLN:NE2 | 2.31 | 0.45 |
| 2:S:57:LEU:HD21 | 2:S:63:LEU:HD11 | 1.99 | 0.45 |
| 3:N:84:ALA:HA | 3:N:102:ARG:HH12 | 1.82 | 0.45 |
| 6:M:11:LEU:HD13 | 6:M:115:THR:OG1 | 2.17 | 0.45 |
| 8:H:24:ARG:NH2 | 8:H:69:THR:OG1 | 2.40 | 0.45 |
| 7:I:93:VAL:HA | 7:I:111:LEU:HA | 1.98 | 0.45 |
| 1:A:21:SER:HB3 | 1:A:79:TYR:CE1 | 2.52 | 0.45 |
| 2:T:134:ARG:NH1 | 5:W:543:TYR:HB2 | 2.31 | 0.45 |
| 2:S:51:LEU:O | 2:S:51:LEU:HD23 | 2.17 | 0.45 |
| 2:S:85:ARG:NH1 | 2:S:178:GLU:OE1 | 2.48 | 0.45 |
| 2:S:198:PRO:HB2 | 5:V:550:ASN:HD21 | 1.82 | 0.45 |
| 6:M:35:ASN:ND2 | 6:M:103:HIS:HE1 | 2.15 | 0.45 |
| 7:I:73:ASP:OD2 | 7:I:73:ASP:N | 2.50 | 0.45 |
| 2:S:123:PRO:HG3 | 2:S:151:PHE:HD2 | 1.81 | 0.44 |
| 2:U:33:ILE:HD12 | 2:U:33:ILE:H | 1.81 | 0.44 |
| 6:O:74:ASN:OD1 | 6:O:75:SER:N | 2.50 | 0.44 |
| 4:D:86:ALA:O | 4:D:89:VAL:HG12 | 2.17 | 0.44 |
| 2:T:115:LYS:HD2 | 2:T:119:SER:HB2 | 1.99 | 0.44 |
| 2:T:182:ALA:HB2 | 5:W:562:ALA:CB | 2.47 | 0.44 |
| 2:U:97:VAL:HG23 | 5:X:573:LEU:HD11 | 1.98 | 0.44 |
| 5:V:585:LEU:HD23 | 5:V:585:LEU:HA | 1.79 | 0.44 |
| 5:X:609:CYS:O | 5:X:609:CYS:SG | 2.75 | 0.44 |
| 7:I:100:TRP:HE1 | 8:J:94:THR:HB | 1.82 | 0.44 |
| 8:J:37:GLN:HE22 | 8:J:39:LYS:HB2 | 1.82 | 0.44 |
| 1:C:110:GLN:NE2 | 4:D:49:PRO:HD3 | 2.32 | 0.44 |
| 7:G:35:HIS:NE2 | 7:G:100:TRP:HA | 2.31 | 0.44 |
| 8:H:38:GLN:O | 8:H:84:ALA:HB1 | 2.18 | 0.44 |
| 3:P:6:GLN:NE2 | 3:P:88:CYS:SG | 2.89 | 0.44 |
| 3:P:78:LEU:HD12 | 3:P:79:GLN:H | 1.83 | 0.44 |
| 2:U:192:ASP:OD1 | 2:U:193:PHE:N | 2.50 | 0.44 |
| 3:N:21:ILE:HD13 | 3:N:101:THR:HG21 | 1.99 | 0.44 |
| 7:I:60:TYR:CZ | 7:I:69:THR:HA | 2.52 | 0.44 |
| 2:U:194:PHE:HA | 5:X:517:TYR:CE2 | 2.52 | 0.44 |
| 3:N:2:ILE:HD11 | 3:N:90:GLN:OE1 | 2.18 | 0.44 |
| 6:Q:50:THR:HG22 | 6:Q:59:TYR:HB2 | 1.99 | 0.44 |

Continued on next page...

For Manuscript Review

Continued from previous page...

| Atom-1 | Atom-2 | Interatomic distance (Å) | Clash overlap (Å) |
|---|---|---|---|
| 8:J:6:GLN:HE22 | 8:J:102:GLY:HA2 | 1.83 | 0.44 |
| 3:N:18:ARG:HA | 3:N:18:ARG:NE | 2.32 | 0.44 |
| 8:H:55:GLN:O | 8:H:58:VAL:HG22 | 2.17 | 0.44 |
| 6:O:62:ASP:OD1 | 6:O:63:SER:N | 2.51 | 0.44 |
| 1:E:4:LEU:HD23 | 1:E:95:CYS:SG | 2.58 | 0.44 |
| 3:R:95:LEU:HD21 | 6:Q:103:HIS:HD1 | 1.83 | 0.44 |
| 3:N:35:TRP:HB2 | 3:N:48:ILE:HG12 | 1.98 | 0.44 |
| 4:F:20:THR:HG23 | 4:F:78:THR:HG23 | 1.99 | 0.44 |
| 8:J:84:ALA:O | 8:J:104:LYS:HA | 2.18 | 0.44 |
| 4:B:86:ALA:HA | 4:B:112:ILE:HG13 | 1.99 | 0.44 |
| 1:A:98:THR:HB | 1:A:102:GLU:O | 2.17 | 0.44 |
| 2:T:87:GLY:O | 2:T:153:PHE:N | 2.33 | 0.44 |
| 6:M:59:TYR:HB3 | 5:X:527:ILE:HD12 | 1.99 | 0.44 |
| 6:M:101:TYR:HE1 | 9:K:2:NAG:C6 | 2.30 | 0.44 |
| 6:M:101:TYR:H | 9:K:5:MAN:H62 | 1.82 | 0.44 |
| 2:T:199:LEU:HD12 | 5:W:549:HIS:NE2 | 2.33 | 0.43 |
| 6:Q:39:GLN:HB2 | 6:Q:45:LEU:HD23 | 1.99 | 0.43 |
| 6:O:34:MET:HB3 | 6:O:79:LEU:HD22 | 2.00 | 0.43 |
| 1:E:31:SER:HB3 | 1:E:100:PHE:CE1 | 2.52 | 0.43 |
| 1:E:39:GLN:O | 1:E:91:ALA:HB1 | 2.18 | 0.43 |
| 4:D:7:SER:HB2 | 4:D:8:PRO:HD3 | 1.99 | 0.43 |
| 2:T:88:PHE:CE2 | 2:T:152:ALA:HB2 | 2.53 | 0.43 |
| 3:R:39:LYS:HD3 | 3:R:84:ALA:HB2 | 1.99 | 0.43 |
| 5:W:513:PRO:HA | 5:W:552:ASP:OD1 | 2.18 | 0.43 |
| 6:Q:1:GLU:HG2 | 6:Q:2:VAL:N | 2.33 | 0.43 |
| 1:C:63:VAL:HB | 1:C:67:PHE:CG | 2.53 | 0.43 |
| 1:A:33:ASP:HA | 1:A:71:ARG:HH12 | 1.83 | 0.43 |
| 4:F:39:LEU:HD13 | 4:F:77:PHE:CD2 | 2.52 | 0.43 |
| 7:G:99:ASN:OD1 | 7:G:100:TRP:N | 2.52 | 0.43 |
| 1:A:51:ILE:HD12 | 1:A:56:ASP:O | 2.18 | 0.43 |
| 3:R:45:LYS:HZ1 | 3:R:58:VAL:HG22 | 1.83 | 0.43 |
| 4:F:6:GLN:NE2 | 4:F:94:CYS:H | 2.17 | 0.43 |
| 5:W:522:ASP:OD1 | 5:W:525:ALA:HB3 | 2.19 | 0.43 |
| 3:P:93:SER:HA | 5:V:528:GLY:HA3 | 2.00 | 0.43 |
| 3:P:99:GLN:HG2 | 3:P:100:GLY:N | 2.33 | 0.43 |
| 5:V:563:ASN:O | 5:V:566:THR:HG22 | 2.18 | 0.43 |
| 7:I:36:TRP:CD2 | 7:I:81:LEU:HD22 | 2.54 | 0.43 |
| 8:J:47:LEU:O | 8:J:54:LEU:HD12 | 2.18 | 0.43 |
| 8:H:13:ALA:HB1 | 8:H:17:ASP:HB3 | 1.99 | 0.43 |
| 5:V:554:LEU:O | 5:V:558:LEU:HG | 2.18 | 0.43 |
| 2:T:193:PHE:CE1 | 5:W:548:MET:HE1 | 2.54 | 0.43 |

Continued on next page...

For Manuscript Review

Continued from previous page...

| Atom-1 | Atom-2 | Interatomic distance (Å) | Clash overlap (Å) |
|---|---|---|---|
| 2:S:234:PHE:HB2 | 2:S:244:LEU:HD21 | 2.01 | 0.43 |
| 2:S:245:GLU:HG3 | 2:S:247:ARG:NH2 | 2.34 | 0.43 |
| 2:U:71:GLU:HG3 | 2:U:77:THR:HG21 | 1.99 | 0.43 |
| 2:U:126:PRO:HD2 | 2:U:129:ILE:HG13 | 2.01 | 0.43 |
| 2:U:162:TYR:OH | 2:U:170:ILE:HA | 2.18 | 0.43 |
| 3:N:33:LEU:O | 3:N:51:ALA:N | 2.47 | 0.43 |
| 3:N:90:GLN:NE2 | 3:N:96:THR:H | 2.09 | 0.43 |
| 8:H:13:ALA:HB2 | 8:H:19:ILE:HD12 | 2.00 | 0.43 |
| 2:T:126:PRO:HB2 | 5:W:580:ARG:NH2 | 2.33 | 0.43 |
| 2:T:154:HIS:NE2 | 2:T:178:GLU:OE1 | 2.41 | 0.43 |
| 1:A:54:ALA:HB3 | 2:U:146:PRO:HG2 | 1.99 | 0.43 |
| 3:R:2:ILE:HD11 | 3:R:27:GLN:HE21 | 1.83 | 0.43 |
| 2:T:296:ASN:O | 2:T:297:LEU:HG | 2.19 | 0.43 |
| 4:F:13:VAL:O | 4:F:112:ILE:HA | 2.19 | 0.43 |
| 6:M:53:GLY:O | 6:M:72:ARG:NH2 | 2.51 | 0.43 |
| 3:P:23:CYS:HB2 | 3:P:35:TRP:CH2 | 2.54 | 0.43 |
| 3:P:55:GLN:NE2 | 3:P:56:SER:H | 2.15 | 0.43 |
| 4:D:11:LEU:O | 4:D:11:LEU:HD23 | 2.19 | 0.43 |
| 2:T:37:VAL:HG23 | 2:T:46:SER:HB2 | 2.01 | 0.43 |
| 2:T:134:ARG:HA | 2:T:134:ARG:HD3 | 1.84 | 0.43 |
| 6:Q:101:TYR:CD1 | 9:L:2:NAG:H4 | 2.54 | 0.43 |
| 6:O:59:TYR:OH | 5:X:567:GLN:HG2 | 2.18 | 0.43 |
| 1:C:12:VAL:HG11 | 1:C:85:LEU:HD12 | 2.01 | 0.43 |
| 1:E:86:ARG:H | 1:E:86:ARG:HG2 | 1.74 | 0.43 |
| 3:N:38:GLN:HE22 | 6:M:45:LEU:HD11 | 1.84 | 0.42 |
| 4:F:60:ARG:HH21 | 4:F:65:PRO:C | 2.22 | 0.42 |
| 7:I:39:GLN:C | 7:I:92:ALA:HB1 | 2.40 | 0.42 |
| 1:E:6:GLU:OE1 | 1:E:111:GLY:HA2 | 2.19 | 0.42 |
| 1:E:43:LYS:HG2 | 1:E:44:GLY:N | 2.30 | 0.42 |
| 2:U:198:PRO:HG2 | 5:X:548:MET:HA | 2.01 | 0.42 |
| 7:G:102:LEU:HB3 | 8:H:34:HIS:CD2 | 2.54 | 0.42 |
| 6:O:13:GLN:H | 6:O:13:GLN:CD | 2.22 | 0.42 |
| 2:T:142:SER:O | 2:T:221:GLN:HA | 2.19 | 0.42 |
| 2:U:234:PHE:CE2 | 2:U:236:VAL:HG12 | 2.54 | 0.42 |
| 6:M:36:TRP:CD1 | 6:M:70:ILE:HD11 | 2.46 | 0.42 |
| 5:X:562:ALA:O | 5:X:565:THR:HG22 | 2.19 | 0.42 |
| 8:J:22:THR:N | 8:J:72:THR:HA | 2.33 | 0.42 |
| 1:C:86:ARG:HG2 | 1:C:89:ASP:OD1 | 2.19 | 0.42 |
| 4:B:21:ILE:HG12 | 4:B:108:THR:HG21 | 2.01 | 0.42 |
| 1:A:58:TYR:HB3 | 2:U:117:ASP:CG | 2.39 | 0.42 |
| 2:T:63:LEU:HB3 | 5:W:585:LEU:HD22 | 2.01 | 0.42 |

Continued on next page...

For Manuscript Review

*Continued from previous page...*

| Atom-1 | Atom-2 | Interatomic distance (Å) | Clash overlap (Å) |
|---|---|---|---|
| 2:T:290:PHE:O | 2:T:295:LYS:HG2 | 2.20 | 0.42 |
| 2:S:164:ARG:HE | 2:S:164:ARG:HB2 | 1.63 | 0.42 |
| 2:S:186:LEU:HD23 | 2:S:186:LEU:HA | 1.83 | 0.42 |
| 4:F:7:SER:HB2 | 4:F:8:PRO:HD3 | 2.01 | 0.42 |
| 6:M:59:TYR:HB3 | 5:X:527:ILE:CD1 | 2.50 | 0.42 |
| 3:P:5:THR:OG1 | 3:P:24:ARG:NH1 | 2.52 | 0.42 |
| 1:C:98:THR:HB | 1:C:102:GLU:O | 2.20 | 0.42 |
| 2:T:57:LEU:HA | 2:T:62:GLN:OE1 | 2.19 | 0.42 |
| 2:U:62:GLN:HE21 | 2:U:62:GLN:HB3 | 1.73 | 0.42 |
| 4:F:45:LYS:HB2 | 4:F:48:GLN:HB2 | 2.01 | 0.42 |
| 5:W:504:ILE:HG23 | 5:W:506:ASN:H | 1.85 | 0.42 |
| 7:G:2:VAL:HB | 7:G:105:TYR:CE1 | 2.54 | 0.42 |
| 3:P:18:ARG:HH12 | 3:P:20:THR:N | 2.18 | 0.42 |
| 7:I:38:ARG:CG | 7:I:92:ALA:HB3 | 2.49 | 0.42 |
| 4:D:89:VAL:HG23 | 4:D:110:VAL:O | 2.19 | 0.42 |
| 2:T:162:TYR:OH | 2:T:170:ILE:HA | 2.19 | 0.42 |
| 5:W:515:LEU:HB2 | 5:W:555:ILE:HD12 | 2.01 | 0.42 |
| 6:Q:34:MET:CB | 6:Q:79:LEU:HD13 | 2.49 | 0.42 |
| 6:Q:39:GLN:HE21 | 6:Q:95:TYR:HE1 | 1.67 | 0.42 |
| 5:V:596:ARG:HB2 | 5:V:597:TRP:CD1 | 2.54 | 0.42 |
| 7:I:17:SER:CB | 7:I:83:MET:HG3 | 2.36 | 0.42 |
| 4:D:14:SER:HA | 4:D:113:LYS:H | 1.84 | 0.42 |
| 6:Q:88:ALA:O | 6:Q:91:THR:HG22 | 2.20 | 0.42 |
| 6:O:59:TYR:HD2 | 5:V:527:ILE:HD13 | 1.85 | 0.42 |
| 2:U:265:LYS:HB3 | 2:U:268:ASN:OD1 | 2.19 | 0.42 |
| 3:R:19:VAL:HB | 3:R:78:LEU:HD21 | 2.01 | 0.42 |
| 4:F:6:GLN:OE1 | 4:F:108:THR:HG23 | 2.19 | 0.42 |
| 8:H:47:LEU:HB2 | 8:H:48:ILE:HD12 | 2.01 | 0.42 |
| 2:U:240:THR:HG21 | 2:U:266:ARG:HG2 | 2.01 | 0.42 |
| 3:N:91:SER:O | 5:X:529:LEU:N | 2.44 | 0.42 |
| 6:M:79:LEU:HD23 | 6:M:80:TYR:N | 2.34 | 0.42 |
| 5:X:513:PRO:HB2 | 5:X:552:ASP:OD1 | 2.20 | 0.42 |
| 1:E:29:PHE:HZ | 1:E:78:LEU:HB2 | 1.85 | 0.42 |
| 4:D:55:TYR:HD2 | 4:D:56:TRP:CD1 | 2.37 | 0.42 |
| 2:T:43:LEU:HB2 | 5:W:554:LEU:HD12 | 2.02 | 0.42 |
| 3:N:61:ARG:NH2 | 3:N:79:GLN:OE1 | 2.52 | 0.42 |
| 5:W:507:ALA:HA | 6:M:32:TYR:CE1 | 2.55 | 0.42 |
| 6:O:32:TYR:HA | 5:X:507:ALA:HB2 | 2.01 | 0.42 |
| 1:C:35:HIS:HD2 | 1:C:47:TRP:NE1 | 2.16 | 0.42 |
| 2:T:300:LYS:HD2 | 2:T:300:LYS:HA | 1.86 | 0.41 |
| 6:Q:17:SER:HB2 | 6:Q:84:ASN:HA | 2.02 | 0.41 |

*Continued on next page...*

For Manuscript Review

*Continued from previous page...*

| Atom-1 | Atom-2 | Interatomic distance (Å) | Clash overlap (Å) |
|---|---|---|---|
| 3:P:6:GLN:HE22 | 3:P:87:TYR:HA | 1.85 | 0.41 |
| 6:O:71:SER:O | 6:O:80:TYR:N | 2.40 | 0.41 |
| 2:U:104:TRP:CH2 | 2:U:134:ARG:HD2 | 2.55 | 0.41 |
| 2:U:291:TRP:CD1 | 2:U:292:GLU:HG3 | 2.55 | 0.41 |
| 4:F:36:LYS:NZ | 4:F:56:TRP:CE2 | 2.88 | 0.41 |
| 6:Q:39:GLN:HG3 | 6:Q:44:GLY:O | 2.20 | 0.41 |
| 4:B:69:SER:OG | 4:B:80:THR:OG1 | 2.35 | 0.41 |
| 2:S:89:ARG:HD2 | 5:W:534:TYR:O | 2.20 | 0.41 |
| 2:S:290:PHE:CE1 | 2:S:291:TRP:HE3 | 2.38 | 0.41 |
| 7:G:73:ASP:OD2 | 7:G:76:LYS:HD3 | 2.21 | 0.41 |
| 8:J:6:GLN:OE1 | 8:J:87:TYR:HA | 2.20 | 0.41 |
| 1:C:34:MET:HB3 | 1:C:78:LEU:HD22 | 2.00 | 0.41 |
| 2:T:123:PRO:HG3 | 2:T:151:PHE:HD2 | 1.85 | 0.41 |
| 2:U:49:ASP:OD1 | 2:U:50:LYS:N | 2.53 | 0.41 |
| 2:U:89:ARG:HD2 | 5:V:534:TYR:O | 2.19 | 0.41 |
| 2:U:186:LEU:HD11 | 2:U:194:PHE:HD1 | 1.86 | 0.41 |
| 6:Q:30:SER:HB3 | 5:V:504:ILE:HG22 | 2.03 | 0.41 |
| 8:H:23:CYS:HB2 | 8:H:35:TRP:CH2 | 2.56 | 0.41 |
| 7:I:36:TRP:CD1 | 7:I:70:ILE:HD13 | 2.55 | 0.41 |
| 7:I:79:LEU:C | 7:I:79:LEU:HD23 | 2.41 | 0.41 |
| 7:I:103:PHE:HB3 | 7:I:106:TRP:NE1 | 2.35 | 0.41 |
| 2:T:48:VAL:HG23 | 2:T:49:ASP:N | 2.35 | 0.41 |
| 2:T:256:LEU:HD11 | 2:T:308:PHE:HE2 | 1.85 | 0.41 |
| 4:F:6:GLN:HA | 4:F:22:ASN:O | 2.21 | 0.41 |
| 5:X:591:ASP:O | 5:X:595:GLN:HG2 | 2.20 | 0.41 |
| 7:I:60:TYR:CE1 | 7:I:70:ILE:HG22 | 2.56 | 0.41 |
| 1:C:99:TRP:CG | 1:C:100:PHE:N | 2.89 | 0.41 |
| 1:E:43:LYS:CG | 1:E:44:GLY:H | 2.31 | 0.41 |
| 4:D:95:GLN:HE21 | 4:D:97:TYR:HB3 | 1.86 | 0.41 |
| 1:A:36:TRP:NE1 | 1:A:78:LEU:HD21 | 2.36 | 0.41 |
| 2:S:126:PRO:HD2 | 2:S:129:ILE:HG13 | 2.02 | 0.41 |
| 2:S:127:ASP:N | 2:S:127:ASP:OD1 | 2.52 | 0.41 |
| 2:U:291:TRP:NE1 | 2:U:292:GLU:HG3 | 2.36 | 0.41 |
| 3:N:92:TYR:O | 5:X:528:GLY:HA3 | 2.21 | 0.41 |
| 6:M:35:ASN:OD1 | 6:M:50:THR:OG1 | 2.36 | 0.41 |
| 5:X:569:LEU:HD12 | 5:X:569:LEU:HA | 1.84 | 0.41 |
| 4:D:13:VAL:HG11 | 4:D:19:ALA:HB2 | 2.02 | 0.41 |
| 1:A:105:PHE:N | 4:B:42:TYR:OH | 2.52 | 0.41 |
| 2:T:133:PRO:HB3 | 5:W:543:TYR:CE1 | 2.56 | 0.41 |
| 2:T:184:LEU:HD23 | 2:T:184:LEU:HA | 1.84 | 0.41 |
| 2:S:94:PRO:HB3 | 2:S:169:VAL:HG11 | 2.02 | 0.41 |

*Continued on next page...*

For Manuscript Review

*Continued from previous page...*

| Atom-1 | Atom-2 | Interatomic distance (Å) | Clash overlap (Å) |
|---|---|---|---|
| 3:N:4:MET:HE2 | 3:N:23:CYS:SG | 2.61 | 0.41 |
| 6:M:47:TRP:CH2 | 5:X:527:ILE:HG13 | 2.56 | 0.41 |
| 8:H:34:HIS:CE1 | 8:H:91:SER:HB2 | 2.54 | 0.41 |
| 5:X:610:ILE:O | 5:X:612:PRO:HD3 | 2.21 | 0.41 |
| 8:J:62:PHE:HA | 8:J:75:ILE:HD12 | 2.02 | 0.41 |
| 2:T:146:PRO:HG2 | 1:C:54:ALA:HB3 | 2.02 | 0.41 |
| 2:T:152:ALA:O | 2:T:169:VAL:HA | 2.20 | 0.41 |
| 2:T:301:ILE:O | 2:T:301:ILE:HG13 | 2.19 | 0.41 |
| 5:W:608:CYS:SG | 5:W:610:ILE:HG12 | 2.61 | 0.41 |
| 7:G:38:ARG:HG3 | 7:G:92:ALA:HB3 | 2.02 | 0.41 |
| 7:I:37:VAL:O | 7:I:95:TYR:N | 2.40 | 0.41 |
| 2:U:198:PRO:HD2 | 5:X:547:LEU:O | 2.20 | 0.41 |
| 5:W:585:LEU:HD23 | 5:W:585:LEU:HA | 1.80 | 0.41 |
| 6:M:54:MET:HA | 6:M:74:ASN:HD21 | 1.86 | 0.41 |
| 7:G:18:LEU:O | 7:G:83:MET:HB2 | 2.20 | 0.41 |
| 7:G:47:TRP:HZ2 | 7:G:50:VAL:HG12 | 1.85 | 0.41 |
| 7:I:92:ALA:O | 7:I:112:VAL:N | 2.48 | 0.41 |
| 4:D:101:PRO:O | 4:D:103:THR:HG23 | 2.21 | 0.41 |
| 2:T:96:VAL:HG12 | 2:T:167:SER:HB2 | 2.02 | 0.41 |
| 3:R:33:LEU:HD23 | 3:R:34:ASN:N | 2.36 | 0.41 |
| 4:F:95:GLN:HG2 | 4:F:96:GLN:N | 2.36 | 0.41 |
| 5:V:515:LEU:CD2 | 5:V:517:TYR:HB3 | 2.51 | 0.41 |
| 1:C:103:LEU:HD12 | 4:D:102:LEU:HD11 | 2.03 | 0.41 |
| 2:U:182:ALA:HB2 | 5:X:562:ALA:CB | 2.49 | 0.40 |
| 4:F:11:LEU:HD11 | 4:F:20:THR:O | 2.22 | 0.40 |
| 4:F:44:GLN:O | 4:F:90:ALA:HB1 | 2.21 | 0.40 |
| 8:H:94:THR:CG2 | 8:H:95:PRO:HD3 | 2.51 | 0.40 |
| 1:E:82:MET:HE3 | 1:E:85:LEU:HD21 | 2.03 | 0.40 |
| 2:S:132:PHE:HD2 | 2:S:135:CYS:SG | 2.44 | 0.40 |
| 2:U:99:TYR:HD2 | 2:U:101:ALA:O | 2.04 | 0.40 |
| 6:Q:101:TYR:CZ | 9:L:3:BMA:H3 | 2.57 | 0.40 |
| 6:O:39:GLN:HB3 | 6:O:95:TYR:HE1 | 1.86 | 0.40 |
| 7:I:45:LEU:HD11 | 8:J:44:PRO:HG2 | 2.02 | 0.40 |
| 8:J:47:LEU:HB3 | 8:J:48:ILE:HD12 | 2.02 | 0.40 |
| 8:J:61:ARG:HH22 | 8:J:79:GLN:CD | 2.25 | 0.40 |
| 1:C:12:VAL:HG11 | 1:C:85:LEU:CD1 | 2.52 | 0.40 |
| 2:U:123:PRO:HG3 | 2:U:151:PHE:CD1 | 2.52 | 0.40 |
| 4:F:1:ASP:HB2 | 4:F:101:PRO:HG3 | 2.04 | 0.40 |
| 6:Q:38:ARG:HG2 | 6:Q:48:VAL:HG22 | 2.04 | 0.40 |
| 6:Q:49:SER:OG | 6:Q:60:TYR:HD1 | 2.05 | 0.40 |
| 8:H:22:THR:HA | 8:H:72:THR:HA | 2.03 | 0.40 |

*Continued on next page...*

For Manuscript Review

*Continued from previous page...*

| Atom-1 | Atom-2 | Interatomic distance (Å) | Clash overlap (Å) |
|---|---|---|---|
| 7:I:51:ILE:HD12 | 7:I:70:ILE:HG23 | 2.02 | 0.40 |
| 1:E:36:TRP:NE1 | 1:E:80:LEU:HB2 | 2.36 | 0.40 |
| 2:S:37:VAL:HG22 | 2:S:185:ILE:HD12 | 2.02 | 0.40 |
| 2:U:309:THR:OG1 | 7:I:57:ASP:OD2 | 2.25 | 0.40 |
| 5:W:601:CYS:HB2 | 5:W:608:CYS:HB2 | 1.44 | 0.40 |
| 6:O:101:TYR:H | 9:Y:5:MAN:H62 | 1.87 | 0.40 |
| 5:V:610:ILE:O | 5:V:610:ILE:HG13 | 2.22 | 0.40 |
| 5:X:584:ILE:O | 5:X:588:LYS:HG2 | 2.22 | 0.40 |
| 2:T:112:GLU:HB2 | 2:T:142:SER:HA | 2.04 | 0.40 |
| 2:U:57:LEU:HD12 | 2:U:62:GLN:HB2 | 2.04 | 0.40 |
| 4:F:60:ARG:NH2 | 4:F:64:VAL:HG23 | 2.37 | 0.40 |
| 3:P:36:TYR:HB2 | 3:P:87:TYR:HB2 | 2.04 | 0.40 |
| 3:P:45:LYS:HE3 | 3:P:45:LYS:HB2 | 1.90 | 0.40 |
| 3:P:94:THR:HA | 5:V:527:ILE:HB | 2.03 | 0.40 |
| 6:O:32:TYR:HD1 | 6:O:98:LYS:CD | 2.35 | 0.40 |
| 5:X:562:ALA:HA | 5:X:565:THR:HG22 | 2.03 | 0.40 |
| 4:D:2:ILE:HD13 | 4:D:99:SER:HB3 | 2.04 | 0.40 |

There are no symmetry-related clashes.

#### 5.3 Torsion angles (i)

##### 5.3.1 Protein backbone (i)

In the following table, the Percentiles column shows the percent Ramachandran outliers of the chain as a percentile score with respect to all PDB entries followed by that with respect to all EM entries.

The Analysed column shows the number of residues for which the backbone conformation was analysed, and the total number of residues.

| Mol | Chain | Analysed | Favoured | Allowed | Outliers | Percentiles | |
|---|---|---|---|---|---|---|---|
| 1 | A | 114/217 (52%) | 113 (99%) | 1 (1%) | 0 | 100 | 100 |
| 1 | C | 114/217 (52%) | 111 (97%) | 3 (3%) | 0 | 100 | 100 |
| 1 | E | 114/217 (52%) | 111 (97%) | 3 (3%) | 0 | 100 | 100 |
| 2 | S | 229/282 (81%) | 225 (98%) | 4 (2%) | 0 | 100 | 100 |
| 2 | T | 265/282 (94%) | 260 (98%) | 5 (2%) | 0 | 100 | 100 |
| 2 | U | 264/282 (94%) | 257 (97%) | 7 (3%) | 0 | 100 | 100 |
| 3 | N | 101/213 (47%) | 97 (96%) | 4 (4%) | 0 | 100 | 100 |

*Continued on next page...*

For Manuscript Review

*Continued from previous page...*

| Mol | Chain | Analysed | Favoured | Allowed | Outliers | Percentiles | |
|---|---|---|---|---|---|---|---|
| 3 | P | 99/213 (46%) | 98 (99%) | 1 (1%) | 0 | 100 | 100 |
| 3 | R | 103/213 (48%) | 99 (96%) | 4 (4%) | 0 | 100 | 100 |
| 4 | B | 111/220 (50%) | 106 (96%) | 5 (4%) | 0 | 100 | 100 |
| 4 | D | 111/220 (50%) | 105 (95%) | 6 (5%) | 0 | 100 | 100 |
| 4 | F | 111/220 (50%) | 105 (95%) | 6 (5%) | 0 | 100 | 100 |
| 5 | V | 108/175 (62%) | 106 (98%) | 2 (2%) | 0 | 100 | 100 |
| 5 | W | 108/175 (62%) | 106 (98%) | 2 (2%) | 0 | 100 | 100 |
| 5 | X | 105/175 (60%) | 102 (97%) | 3 (3%) | 0 | 100 | 100 |
| 6 | M | 113/217 (52%) | 110 (97%) | 2 (2%) | 1 (1%) | 17 | 52 |
| 6 | O | 113/217 (52%) | 109 (96%) | 4 (4%) | 0 | 100 | 100 |
| 6 | Q | 114/217 (52%) | 113 (99%) | 1 (1%) | 0 | 100 | 100 |
| 7 | G | 113/215 (53%) | 110 (97%) | 3 (3%) | 0 | 100 | 100 |
| 7 | I | 111/215 (52%) | 109 (98%) | 2 (2%) | 0 | 100 | 100 |
| 8 | H | 105/215 (49%) | 99 (94%) | 6 (6%) | 0 | 100 | 100 |
| 8 | J | 91/215 (42%) | 89 (98%) | 1 (1%) | 1 (1%) | 14 | 46 |
| All | All | 2817/4832 (58%) | 2740 (97%) | 75 (3%) | 2 (0%) | 54 | 83 |

All (2) Ramachandran outliers are listed below:

| Mol | Chain | Res | Type |
|---|---|---|---|
| 6 | M | 102 | PRO |
| 8 | J | 96 | PRO |

##### 5.3.2 Protein sidechains (i)

In the following table, the Percentiles column shows the percent sidechain outliers of the chain as a percentile score with respect to all PDB entries followed by that with respect to all EM entries.

The Analysed column shows the number of residues for which the sidechain conformation was analysed, and the total number of residues.

| Mol | Chain | Analysed | Rotameric | Outliers | Percentiles | |
|---|---|---|---|---|---|---|
| 1 | A | 91/178 (51%) | 91 (100%) | 0 | 100 | 100 |
| 1 | C | 91/178 (51%) | 91 (100%) | 0 | 100 | 100 |
| 1 | E | 91/178 (51%) | 91 (100%) | 0 | 100 | 100 |

*Continued on next page...*

For Manuscript Review

*Continued from previous page...*

| Mol | Chain | Analysed | Rotameric | Outliers | Percentiles | |
|---|---|---|---|---|---|---|
| 2 | S | 201/244 (82%) | 201 (100%) | 0 | 100 | 100 |
| 2 | T | 232/244 (95%) | 232 (100%) | 0 | 100 | 100 |
| 2 | U | 232/244 (95%) | 232 (100%) | 0 | 100 | 100 |
| 3 | N | 89/188 (47%) | 89 (100%) | 0 | 100 | 100 |
| 3 | P | 87/188 (46%) | 87 (100%) | 0 | 100 | 100 |
| 3 | R | 91/188 (48%) | 91 (100%) | 0 | 100 | 100 |
| 4 | B | 98/194 (50%) | 98 (100%) | 0 | 100 | 100 |
| 4 | D | 98/194 (50%) | 98 (100%) | 0 | 100 | 100 |
| 4 | F | 98/194 (50%) | 98 (100%) | 0 | 100 | 100 |
| 5 | V | 90/144 (62%) | 90 (100%) | 0 | 100 | 100 |
| 5 | W | 90/144 (62%) | 89 (99%) | 1 (1%) | 73 | 89 |
| 5 | X | 87/144 (60%) | 87 (100%) | 0 | 100 | 100 |
| 6 | M | 92/180 (51%) | 92 (100%) | 0 | 100 | 100 |
| 6 | O | 92/180 (51%) | 92 (100%) | 0 | 100 | 100 |
| 6 | Q | 93/180 (52%) | 93 (100%) | 0 | 100 | 100 |
| 7 | G | 95/181 (52%) | 95 (100%) | 0 | 100 | 100 |
| 7 | I | 93/181 (51%) | 92 (99%) | 1 (1%) | 73 | 89 |
| 8 | H | 92/190 (48%) | 92 (100%) | 0 | 100 | 100 |
| 8 | J | 83/190 (44%) | 82 (99%) | 1 (1%) | 71 | 88 |
| All | All | 2406/4126 (58%) | 2403 (100%) | 3 (0%) | 93 | 98 |

All (3) residues with a non-rotameric sidechain are listed below:

| Mol | Chain | Res | Type |
|---|---|---|---|
| 5 | W | 601 | CYS |
| 7 | I | 74 | ASN |
| 8 | J | 104 | LYS |

Sometimes sidechains can be flipped to improve hydrogen bonding and reduce clashes. All (18) such sidechains are listed below:

| Mol | Chain | Res | Type |
|---|---|---|---|
| 1 | A | 3 | GLN |
| 2 | T | 98 | ASN |
| 2 | T | 296 | ASN |

*Continued on next page...*

For Manuscript Review

*Continued from previous page...*

| Mol | Chain | Res | Type |
|---|---|---|---|
| 2 | S | 62 | GLN |
| 2 | S | 255 | GLN |
| 3 | R | 38 | GLN |
| 5 | W | 595 | GLN |
| 6 | M | 103 | HIS |
| 8 | H | 34 | HIS |
| 3 | P | 38 | GLN |
| 6 | O | 39 | GLN |
| 5 | V | 521 | GLN |
| 5 | V | 567 | GLN |
| 5 | X | 521 | GLN |
| 8 | J | 6 | GLN |
| 8 | J | 37 | GLN |
| 1 | C | 35 | HIS |
| 1 | E | 35 | HIS |

##### 5.3.3 RNA (i)

There are no RNA molecules in this entry.

#### 5.4 Non-standard residues in protein, DNA, RNA chains (i)

There are no non-standard protein/DNA/RNA residues in this entry.

#### 5.5 Carbohydrates (i)

18 monosaccharides are modelled in this entry.

In the following table, the Counts columns list the number of bonds (or angles) for which Mogul statistics could be retrieved, the number of bonds (or angles) that are observed in the model and the number of bonds (or angles) that are defined in the Chemical Component Dictionary. The Link column lists molecule types, if any, to which the group is linked. The Z score for a bond length (or angle) is the number of standard deviations the observed value is removed from the expected value. A bond length (or angle) with $|Z| > 2$ is considered an outlier worth inspection. RMSZ is the root-mean-square of all Z scores of the bond lengths (or angles).

| Mol | Type | Chain | Res | Link | Bond lengths | | | Bond angles | | |
|---|---|---|---|---|---|---|---|---|---|---|
| | | | | | Counts | RMSZ | $\#\|Z\|>2$ | Counts | RMSZ | $\#\|Z\|>2$ |
| 9 | NAG | K | 1 | 5,9 | 14,14,15 | 0.47 | 0 | 17,19,21 | 0.68 | 0 |
| 9 | NAG | K | 2 | 9 | 14,14,15 | 0.29 | 0 | 17,19,21 | 0.54 | 0 |
| 9 | BMA | K | 3 | 9 | 11,11,12 | 0.87 | 0 | 15,15,17 | 1.26 | 2 (13%) |

For Manuscript Review

| Mol | Type | Chain | Res | Link | Bond lengths | | | Bond angles | | |
|---|---|---|---|---|---|---|---|---|---|---|
| | | | | | Counts | RMSZ | #\|Z\| > 2 | Counts | RMSZ | #\|Z\| > 2 |
| 9 | MAN | K | 4 | 9 | 11,11,12 | 1.11 | 1 (9%) | 15,15,17 | 2.64 | 6 (40%) |
| 9 | MAN | K | 5 | 9 | 11,11,12 | 0.67 | 0 | 15,15,17 | 0.94 | 1 (6%) |
| 9 | MAN | K | 6 | 9 | 11,11,12 | 0.90 | 1 (9%) | 15,15,17 | 1.01 | 1 (6%) |
| 9 | NAG | L | 1 | 5,9 | 14,14,15 | 0.40 | 0 | 17,19,21 | 0.52 | 0 |
| 9 | NAG | L | 2 | 9 | 14,14,15 | 0.18 | 0 | 17,19,21 | 0.46 | 0 |
| 9 | BMA | L | 3 | 9 | 11,11,12 | 0.69 | 0 | 15,15,17 | 1.08 | 1 (6%) |
| 9 | MAN | L | 4 | 9 | 11,11,12 | 0.63 | 0 | 15,15,17 | 1.06 | 2 (13%) |
| 9 | MAN | L | 5 | 9 | 11,11,12 | 0.77 | 0 | 15,15,17 | 0.91 | 1 (6%) |
| 9 | MAN | L | 6 | 9 | 11,11,12 | 0.82 | 1 (9%) | 15,15,17 | 0.89 | 1 (6%) |
| 9 | NAG | Y | 1 | 5,9 | 14,14,15 | 0.45 | 0 | 17,19,21 | 0.45 | 0 |
| 9 | NAG | Y | 2 | 9 | 14,14,15 | 0.46 | 0 | 17,19,21 | 0.41 | 0 |
| 9 | BMA | Y | 3 | 9 | 11,11,12 | 0.67 | 0 | 15,15,17 | 1.23 | 3 (20%) |
| 9 | MAN | Y | 4 | 9 | 11,11,12 | 0.79 | 1 (9%) | 15,15,17 | 1.14 | 2 (13%) |
| 9 | MAN | Y | 5 | 9 | 11,11,12 | 1.05 | 1 (9%) | 15,15,17 | 0.89 | 1 (6%) |
| 9 | MAN | Y | 6 | 9 | 11,11,12 | 0.81 | 1 (9%) | 15,15,17 | 0.94 | 1 (6%) |

In the following table, the Chirals column lists the number of chiral outliers, the number of chiral centers analysed, the number of these observed in the model and the number defined in the Chemical Component Dictionary. Similar counts are reported in the Torsion and Rings columns. '-' means no outliers of that kind were identified.

| Mol | Type | Chain | Res | Link | Chirals | Torsions | Rings |
|---|---|---|---|---|---|---|---|
| 9 | NAG | K | 1 | 5,9 | - | 0/6/23/26 | 0/1/1/1 |
| 9 | NAG | K | 2 | 9 | - | 2/6/23/26 | 0/1/1/1 |
| 9 | BMA | K | 3 | 9 | - | 2/2/19/22 | 0/1/1/1 |
| 9 | MAN | K | 4 | 9 | - | 2/2/19/22 | 0/1/1/1 |
| 9 | MAN | K | 5 | 9 | - | 1/2/19/22 | 0/1/1/1 |
| 9 | MAN | K | 6 | 9 | - | 0/2/19/22 | 0/1/1/1 |
| 9 | NAG | L | 1 | 5,9 | - | 0/6/23/26 | 0/1/1/1 |
| 9 | NAG | L | 2 | 9 | - | 2/6/23/26 | 0/1/1/1 |
| 9 | BMA | L | 3 | 9 | - | 2/2/19/22 | 0/1/1/1 |
| 9 | MAN | L | 4 | 9 | - | 1/2/19/22 | 1/1/1/1 |
| 9 | MAN | L | 5 | 9 | - | 2/2/19/22 | 0/1/1/1 |
| 9 | MAN | L | 6 | 9 | - | 0/2/19/22 | 0/1/1/1 |
| 9 | NAG | Y | 1 | 5,9 | - | 1/6/23/26 | 0/1/1/1 |
| 9 | NAG | Y | 2 | 9 | - | 2/6/23/26 | 0/1/1/1 |
| 9 | BMA | Y | 3 | 9 | - | 0/2/19/22 | 0/1/1/1 |
| 9 | MAN | Y | 4 | 9 | - | 0/2/19/22 | 0/1/1/1 |

*Continued on next page...*

For Manuscript Review

*Continued from previous page...*

| Mol | Type | Chain | Res | Link | Chirals | Torsions | Rings |
|---|---|---|---|---|---|---|---|
| 9 | MAN | Y | 5 | 9 | - | 1/2/19/22 | 0/1/1/1 |
| 9 | MAN | Y | 6 | 9 | - | 2/2/19/22 | 0/1/1/1 |

All (6) bond length outliers are listed below:

| Mol | Chain | Res | Type | Atoms | Z | Observed(Å) | Ideal(Å) |
|---|---|---|---|---|---|---|---|
| 9 | Y | 5 | MAN | O5-C1 | -3.28 | 1.38 | 1.43 |
| 9 | K | 4 | MAN | C1-C2 | 2.86 | 1.58 | 1.52 |
| 9 | K | 6 | MAN | O5-C1 | -2.45 | 1.39 | 1.43 |
| 9 | L | 6 | MAN | O5-C1 | -2.40 | 1.39 | 1.43 |
| 9 | Y | 4 | MAN | C1-C2 | 2.33 | 1.57 | 1.52 |
| 9 | Y | 6 | MAN | O5-C1 | -2.10 | 1.40 | 1.43 |

All (22) bond angle outliers are listed below:

| Mol | Chain | Res | Type | Atoms | Z | Observed($^o$) | Ideal($^o$) |
|---|---|---|---|---|---|---|---|
| 9 | K | 4 | MAN | C1-O5-C5 | 6.10 | 120.46 | 112.19 |
| 9 | K | 4 | MAN | C1-C2-C3 | 4.12 | 114.73 | 109.67 |
| 9 | K | 4 | MAN | O5-C5-C6 | 4.06 | 113.58 | 107.20 |
| 9 | K | 3 | BMA | C1-C2-C3 | -3.32 | 105.58 | 109.67 |
| 9 | K | 4 | MAN | O5-C5-C4 | 2.89 | 117.86 | 110.83 |
| 9 | K | 4 | MAN | O5-C1-C2 | 2.86 | 115.18 | 110.77 |
| 9 | K | 4 | MAN | O2-C2-C3 | -2.79 | 104.56 | 110.14 |
| 9 | Y | 6 | MAN | O2-C2-C3 | -2.57 | 104.99 | 110.14 |
| 9 | K | 6 | MAN | O2-C2-C3 | -2.48 | 105.17 | 110.14 |
| 9 | Y | 4 | MAN | O2-C2-C3 | -2.46 | 105.20 | 110.14 |
| 9 | L | 6 | MAN | O2-C2-C3 | -2.42 | 105.28 | 110.14 |
| 9 | K | 5 | MAN | O2-C2-C3 | -2.37 | 105.38 | 110.14 |
| 9 | L | 4 | MAN | C1-O5-C5 | 2.37 | 115.40 | 112.19 |
| 9 | L | 5 | MAN | O2-C2-C3 | -2.36 | 105.41 | 110.14 |
| 9 | L | 4 | MAN | O2-C2-C3 | -2.34 | 105.45 | 110.14 |
| 9 | Y | 3 | BMA | O5-C5-C6 | 2.32 | 110.84 | 107.20 |
| 9 | Y | 3 | BMA | C1-O5-C5 | 2.31 | 115.33 | 112.19 |
| 9 | Y | 5 | MAN | O2-C2-C3 | -2.29 | 105.55 | 110.14 |
| 9 | Y | 3 | BMA | O5-C5-C4 | -2.25 | 105.35 | 110.83 |
| 9 | Y | 4 | MAN | C1-O5-C5 | 2.15 | 115.10 | 112.19 |
| 9 | K | 3 | BMA | O3-C3-C4 | 2.08 | 115.16 | 110.35 |
| 9 | L | 3 | BMA | C1-O5-C5 | 2.06 | 114.98 | 112.19 |

There are no chirality outliers.

All (20) torsion outliers are listed below:

For Manuscript Review

| Mol | Chain | Res | Type | Atoms |
|---|---|---|---|---|
| 9 | L | 2 | NAG | O5-C5-C6-O6 |
| 9 | K | 2 | NAG | O5-C5-C6-O6 |
| 9 | Y | 2 | NAG | O5-C5-C6-O6 |
| 9 | K | 2 | NAG | C4-C5-C6-O6 |
| 9 | K | 4 | MAN | O5-C5-C6-O6 |
| 9 | L | 2 | NAG | C4-C5-C6-O6 |
| 9 | K | 3 | BMA | C4-C5-C6-O6 |
| 9 | Y | 2 | NAG | C4-C5-C6-O6 |
| 9 | L | 5 | MAN | O5-C5-C6-O6 |
| 9 | L | 4 | MAN | O5-C5-C6-O6 |
| 9 | L | 5 | MAN | C4-C5-C6-O6 |
| 9 | K | 3 | BMA | O5-C5-C6-O6 |
| 9 | K | 4 | MAN | C4-C5-C6-O6 |
| 9 | L | 3 | BMA | C4-C5-C6-O6 |
| 9 | Y | 5 | MAN | O5-C5-C6-O6 |
| 9 | Y | 1 | NAG | O5-C5-C6-O6 |
| 9 | Y | 6 | MAN | O5-C5-C6-O6 |
| 9 | K | 5 | MAN | C4-C5-C6-O6 |
| 9 | L | 3 | BMA | O5-C5-C6-O6 |
| 9 | Y | 6 | MAN | C4-C5-C6-O6 |

All (1) ring outliers are listed below:

| Mol | Chain | Res | Type | Atoms |
|---|---|---|---|---|
| 9 | L | 4 | MAN | C1-C2-C3-C4-C5-O5 |

10 monomers are involved in 15 short contacts:

| Mol | Chain | Res | Type | Clashes | Symm-Clashes |
|---|---|---|---|---|---|
| 9 | K | 3 | BMA | 1 | 0 |
| 9 | L | 3 | BMA | 1 | 0 |
| 9 | Y | 5 | MAN | 1 | 0 |
| 9 | K | 2 | NAG | 3 | 0 |
| 9 | Y | 2 | NAG | 2 | 0 |
| 9 | L | 2 | NAG | 2 | 0 |
| 9 | L | 1 | NAG | 1 | 0 |
| 9 | Y | 1 | NAG | 1 | 0 |
| 9 | K | 1 | NAG | 1 | 0 |
| 9 | K | 5 | MAN | 2 | 0 |

The following is a two-dimensional graphical depiction of Mogul quality analysis of bond lengths, bond angles, torsion angles, and ring geometry for oligosaccharide.

For Manuscript Review

**Oligosaccharide Chain K**

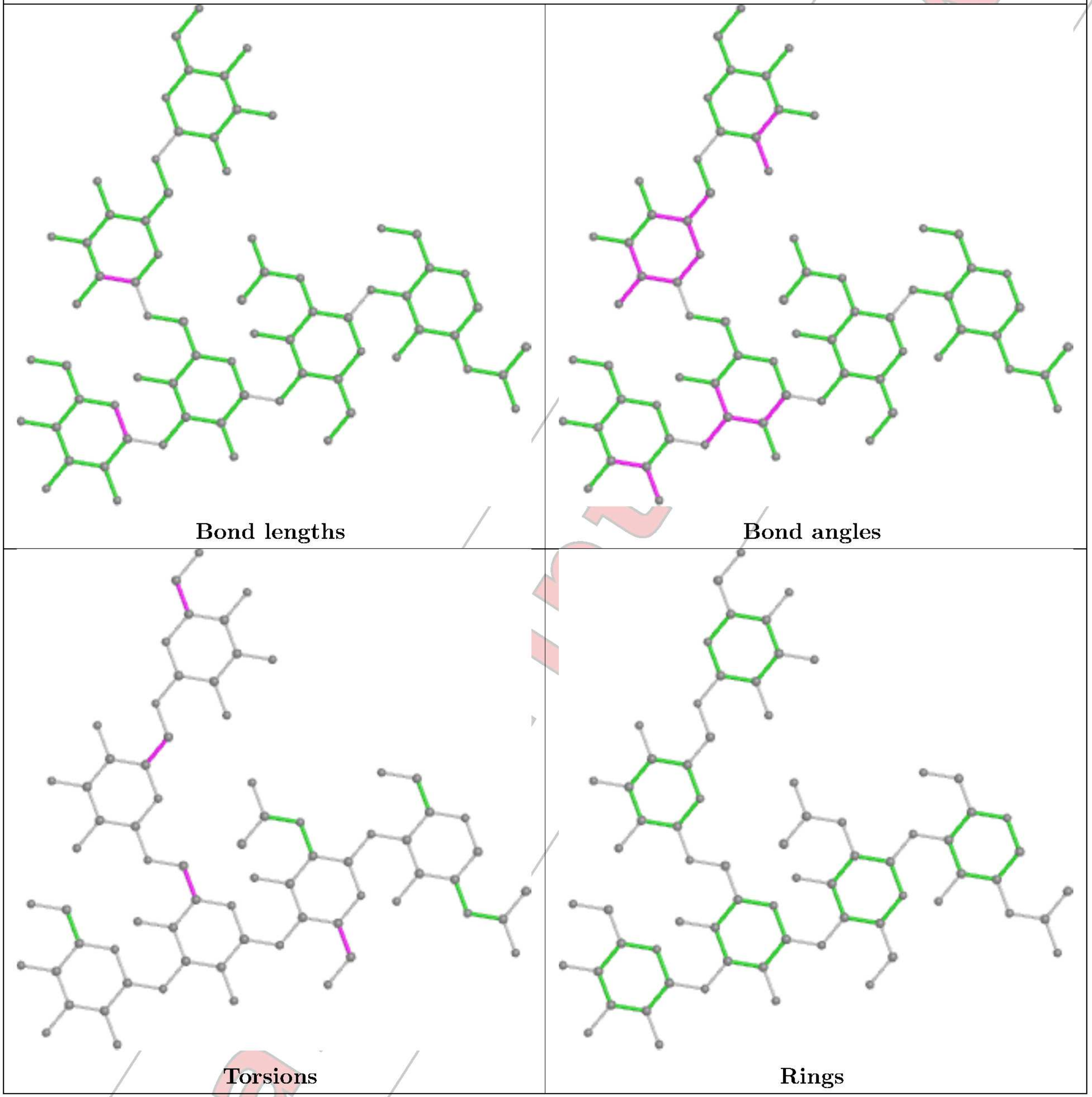

**Oligosaccharide Chain L**

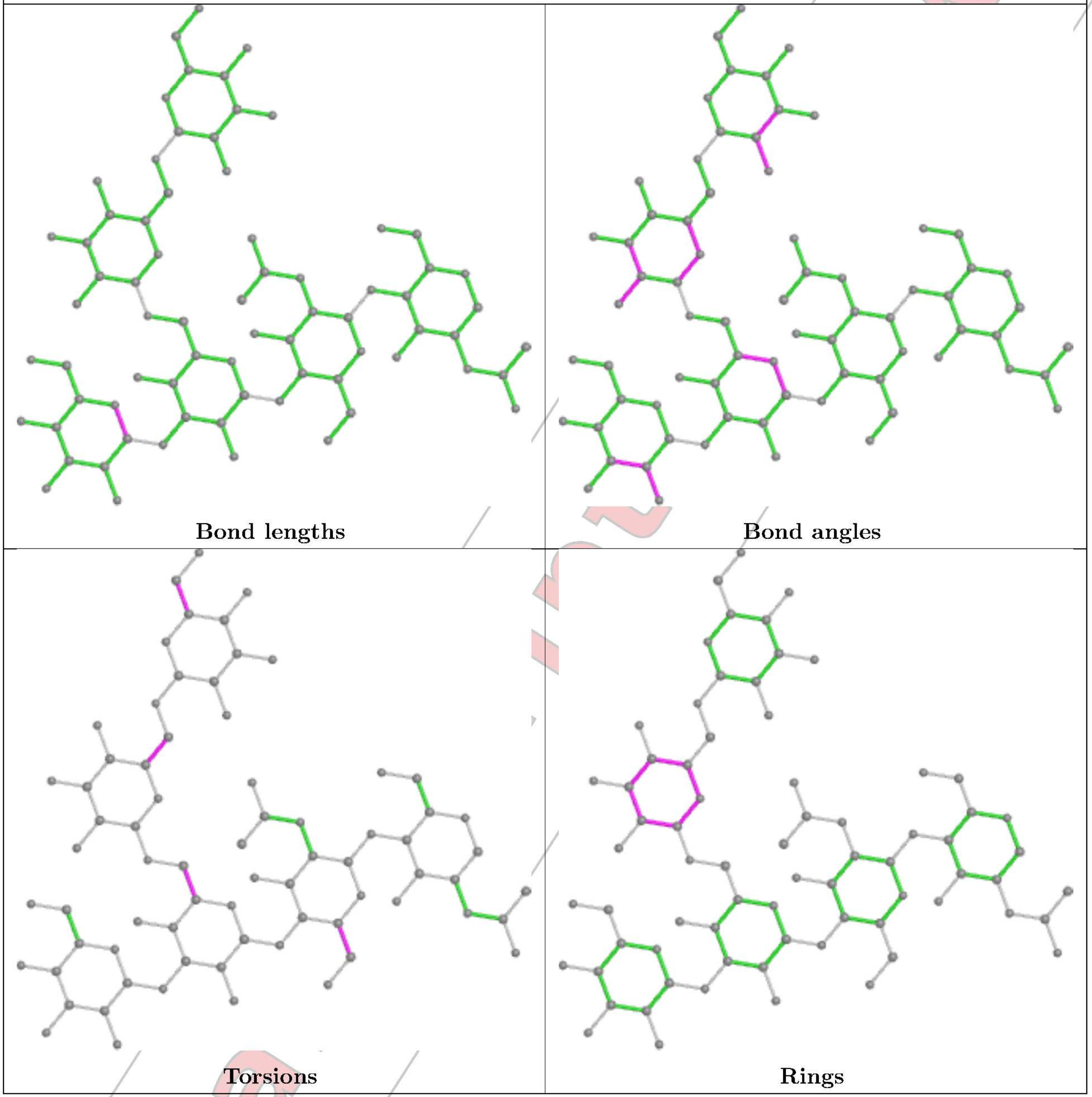


For Manuscript Review

**Oligosaccharide Chain Y**

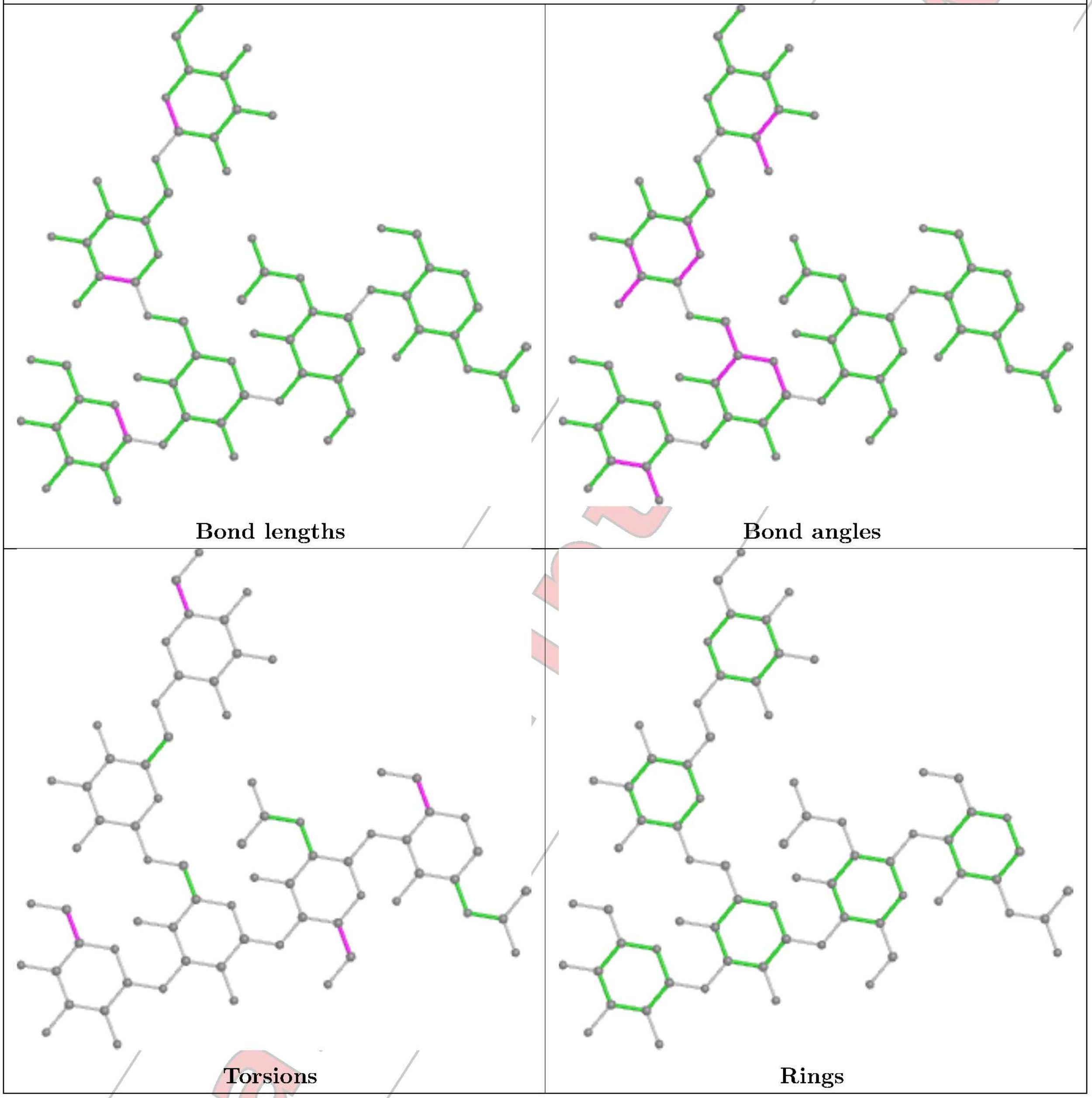


#### 5.6 Ligand geometry (i)

8 ligands are modelled in this entry.

In the following table, the Counts columns list the number of bonds (or angles) for which Mogul statistics could be retrieved, the number of bonds (or angles) that are observed in the model and the number of bonds (or angles) that are defined in the Chemical Component Dictionary. The Link column lists molecule types, if any, to which the group is linked. The Z score for a bond length (or angle) is the number of standard deviations the observed value is removed from the expected value. A bond length (or angle) with $|Z| > 2$ is considered an outlier worth inspection. RMSZ is the root-mean-square of all Z scores of the bond lengths (or angles).

For Manuscript Review

| Mol | Type | Chain | Res | Link | Bond lengths | | | Bond angles | | |
|---|---|---|---|---|---|---|---|---|---|---|
| | | | | | Counts | RMSZ | #\|Z\| > 2 | Counts | RMSZ | #\|Z\| > 2 |
| 10 | NAG | U | 503 | 2 | 14,14,15 | 0.26 | 0 | 17,19,21 | 0.51 | 0 |
| 10 | NAG | T | 502 | 2 | 14,14,15 | 0.34 | 0 | 17,19,21 | 0.45 | 0 |
| 10 | NAG | S | 401 | 2 | 14,14,15 | 0.21 | 0 | 17,19,21 | 0.38 | 0 |
| 10 | NAG | S | 402 | 2 | 14,14,15 | 0.29 | 0 | 17,19,21 | 0.40 | 0 |
| 10 | NAG | T | 501 | 2 | 14,14,15 | 0.20 | 0 | 17,19,21 | 0.53 | 0 |
| 10 | NAG | U | 502 | 2 | 14,14,15 | 0.22 | 0 | 17,19,21 | 0.50 | 0 |
| 10 | NAG | U | 501 | 2 | 14,14,15 | 0.25 | 0 | 17,19,21 | 0.38 | 0 |
| 10 | NAG | T | 503 | 2 | 14,14,15 | 0.58 | 0 | 17,19,21 | 0.47 | 0 |

In the following table, the Chirals column lists the number of chiral outliers, the number of chiral centers analysed, the number of these observed in the model and the number defined in the Chemical Component Dictionary. Similar counts are reported in the Torsion and Rings columns. '-' means no outliers of that kind were identified.

| Mol | Type | Chain | Res | Link | Chirals | Torsions | Rings |
|---|---|---|---|---|---|---|---|
| 10 | NAG | U | 503 | 2 | - | 2/6/23/26 | 0/1/1/1 |
| 10 | NAG | T | 502 | 2 | - | 0/6/23/26 | 0/1/1/1 |
| 10 | NAG | S | 401 | 2 | - | 0/6/23/26 | 0/1/1/1 |
| 10 | NAG | S | 402 | 2 | - | 0/6/23/26 | 0/1/1/1 |
| 10 | NAG | T | 501 | 2 | - | 2/6/23/26 | 0/1/1/1 |
| 10 | NAG | U | 502 | 2 | - | 0/6/23/26 | 0/1/1/1 |
| 10 | NAG | U | 501 | 2 | - | 2/6/23/26 | 0/1/1/1 |
| 10 | NAG | T | 503 | 2 | - | 2/6/23/26 | 0/1/1/1 |

There are no bond length outliers.

There are no bond angle outliers.

There are no chirality outliers.

All (8) torsion outliers are listed below:

| Mol | Chain | Res | Type | Atoms |
|---|---|---|---|---|
| 10 | T | 501 | NAG | C4-C5-C6-O6 |
| 10 | T | 501 | NAG | O5-C5-C6-O6 |
| 10 | T | 503 | NAG | C1-C2-N2-C7 |
| 10 | U | 501 | NAG | C4-C5-C6-O6 |
| 10 | U | 503 | NAG | O5-C5-C6-O6 |
| 10 | U | 501 | NAG | O5-C5-C6-O6 |
| 10 | T | 503 | NAG | C3-C2-N2-C7 |
| 10 | U | 503 | NAG | C3-C2-N2-C7 |

There are no ring outliers.

For Manuscript Review

1 monomer is involved in 1 short contact:

| Mol | Chain | Res | Type | Clashes | Symm-Clashes |
|---|---|---|---|---|---|
| 10 | T | 503 | NAG | 1 | 0 |

The following is a two-dimensional graphical depiction of Mogul quality analysis of bond lengths, bond angles, torsion angles, and ring geometry for all instances of the Ligand of Interest. In addition, ligands with molecular weight > 250 and outliers as shown on the validation Tables will also be included. For torsion angles, if less then 5% of the Mogul distribution of torsion angles is within 10 degrees of the torsion angle in question, then that torsion angle is considered an outlier. Any bond that is central to one or more torsion angles identified as an outlier by Mogul will be highlighted in the graph. For rings, the root-mean-square deviation (RMSD) between the ring in question and similar rings identified by Mogul is calculated over all ring torsion angles. If the average RMSD is greater than 60 degrees and the minimal RMSD between the ring in question and any Mogul-identified rings is also greater than 60 degrees, then that ring is considered an outlier. The outliers are highlighted in purple. The color gray indicates Mogul did not find sufficient equivalents in the CSD to analyse the geometry.

**Ligand NAG U 503**

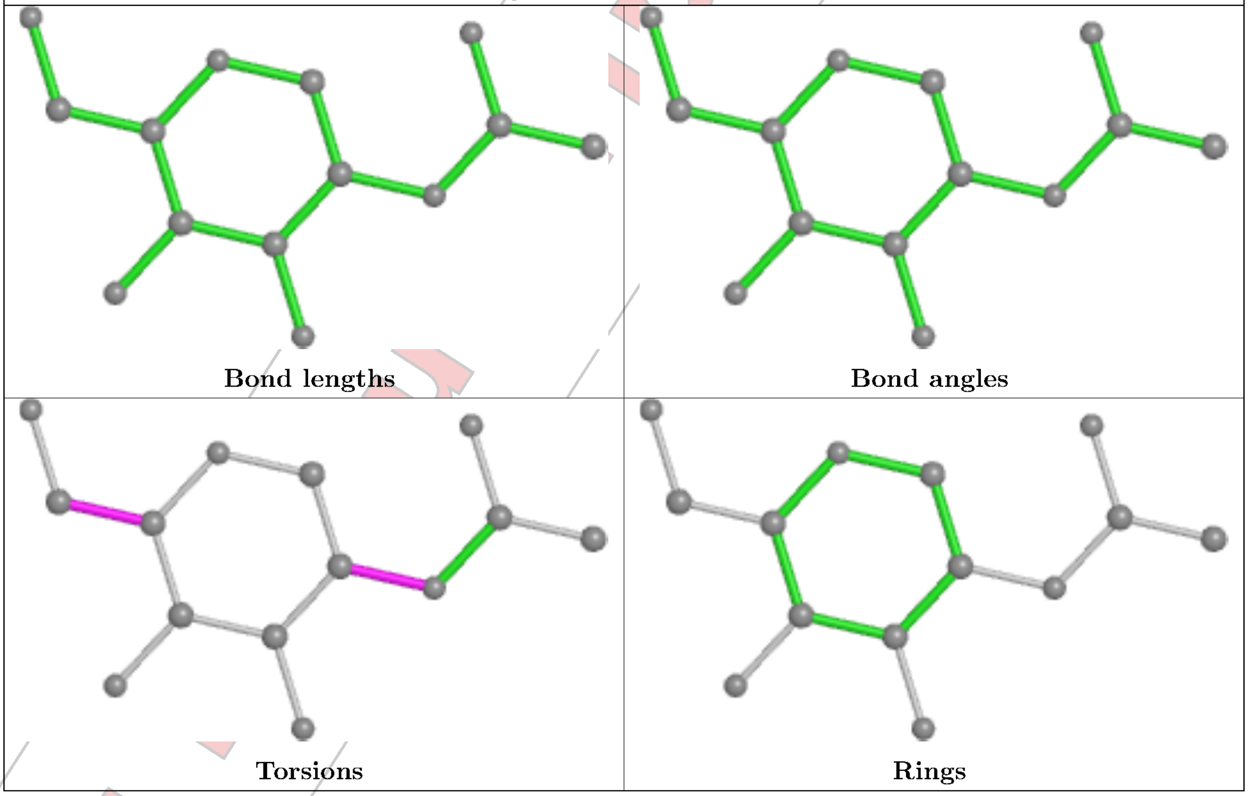


For Review

**Ligand NAG T 502**

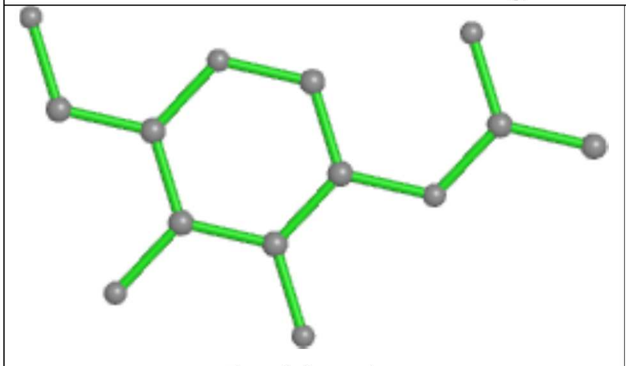
Bond lengths

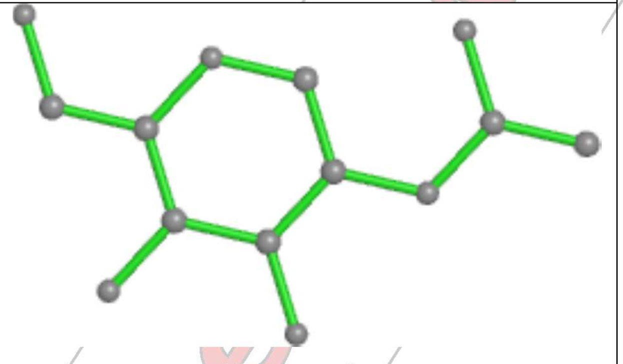
Bond angles

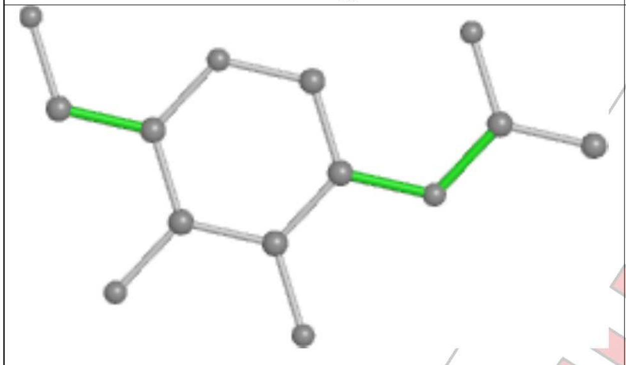
Torsions

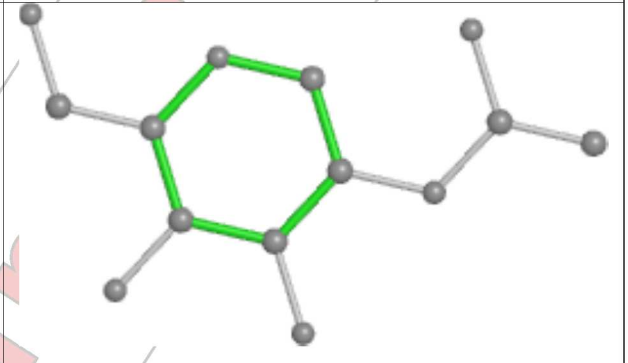
Rings

**Ligand NAG S 401**

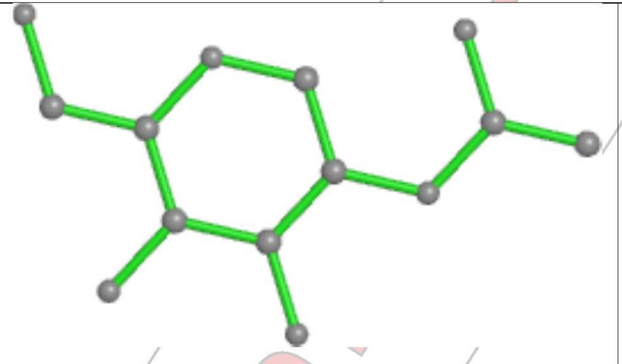
Bond lengths

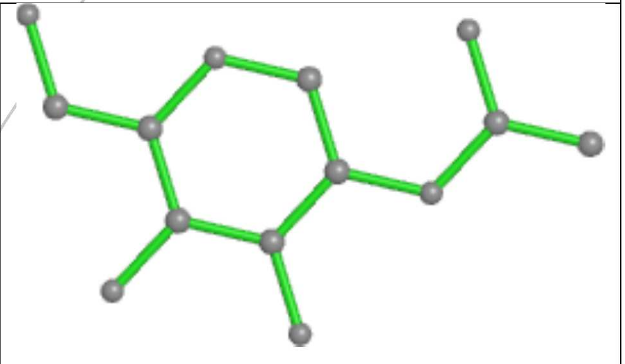
Bond angles

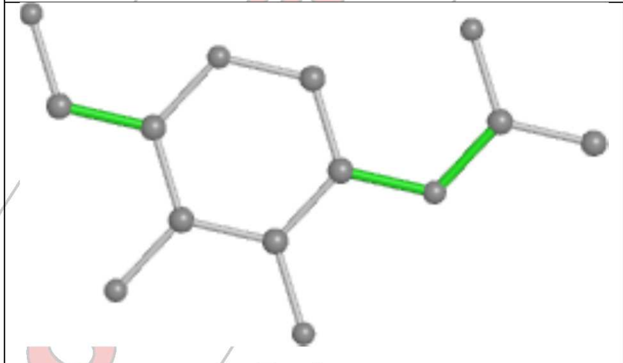
Torsions

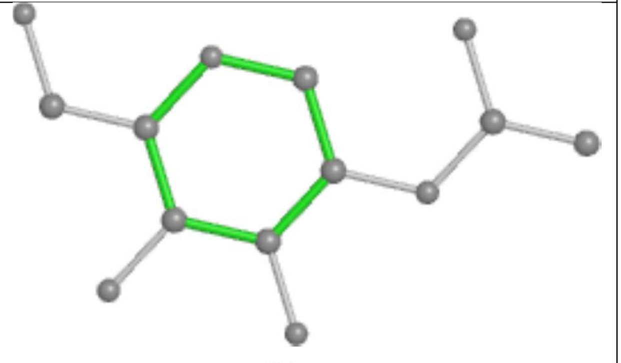
Rings

**Ligand NAG S 402**

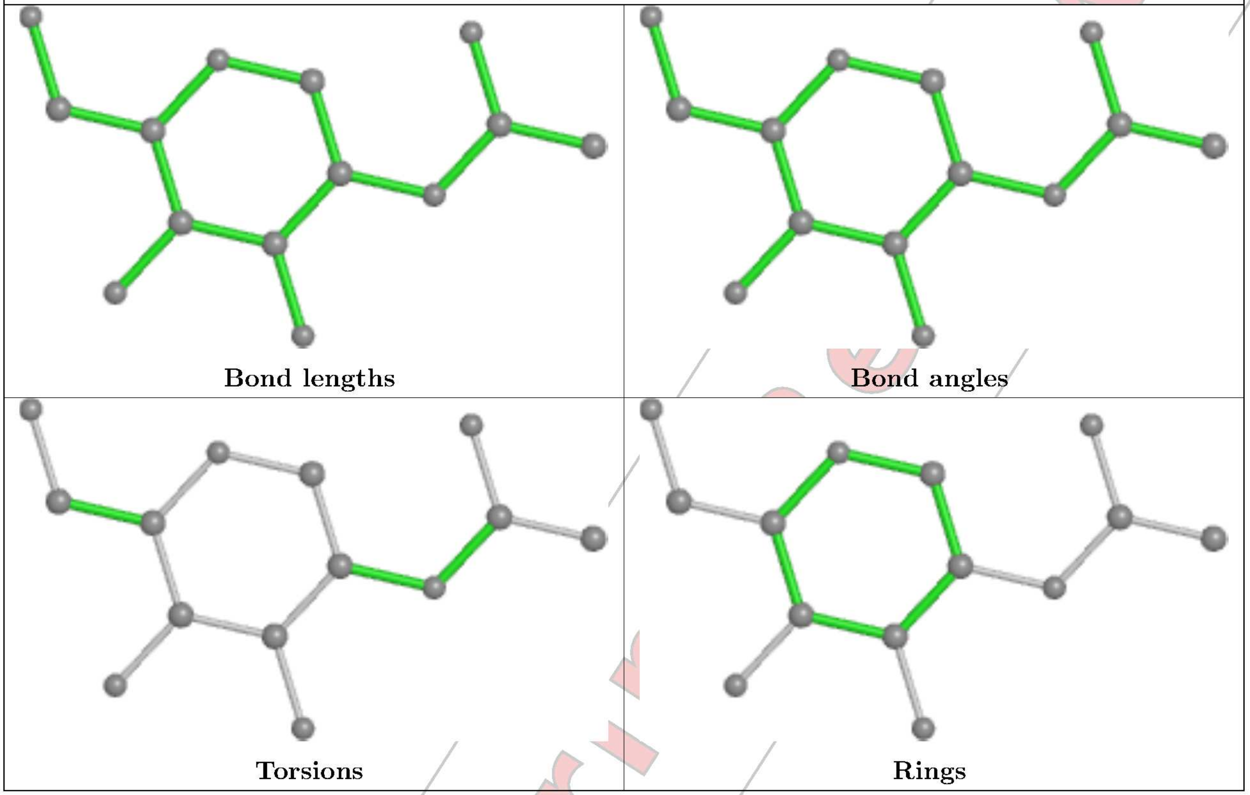


**Ligand NAG T 501**

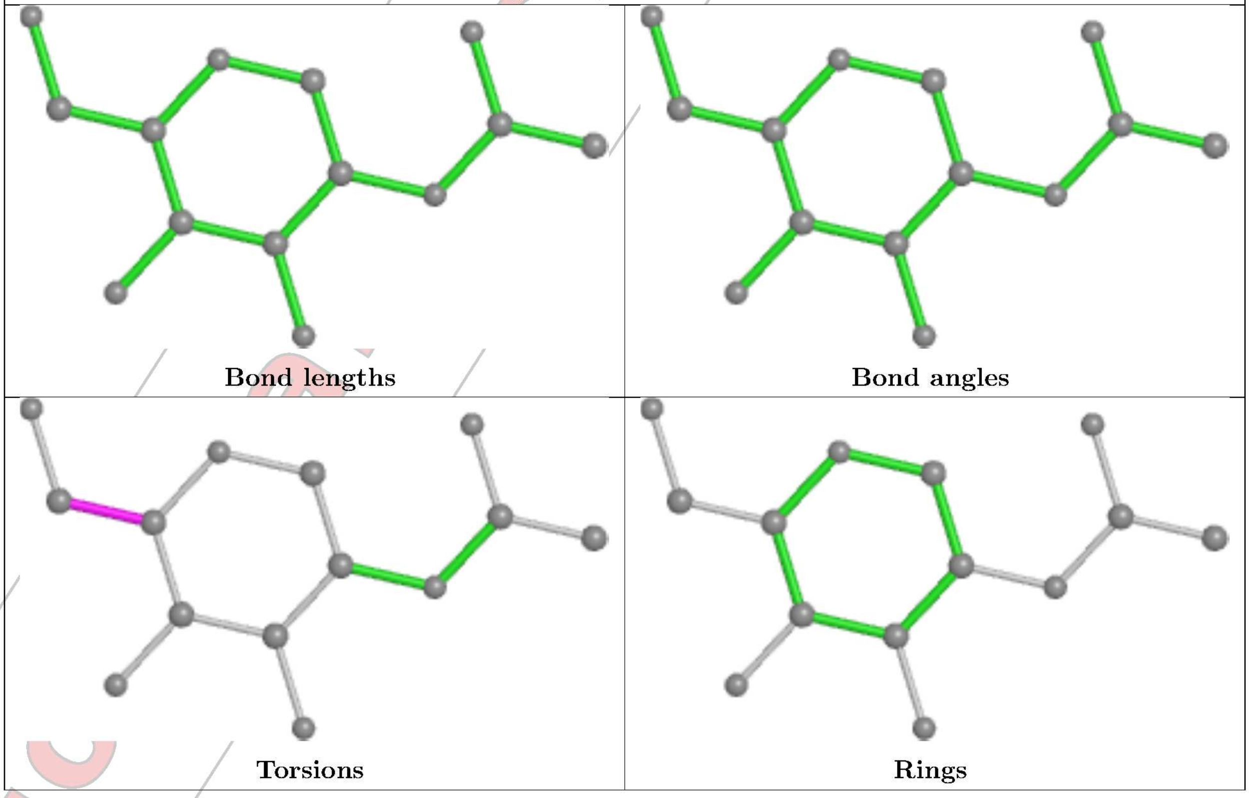

#### Ligand NAG U 502

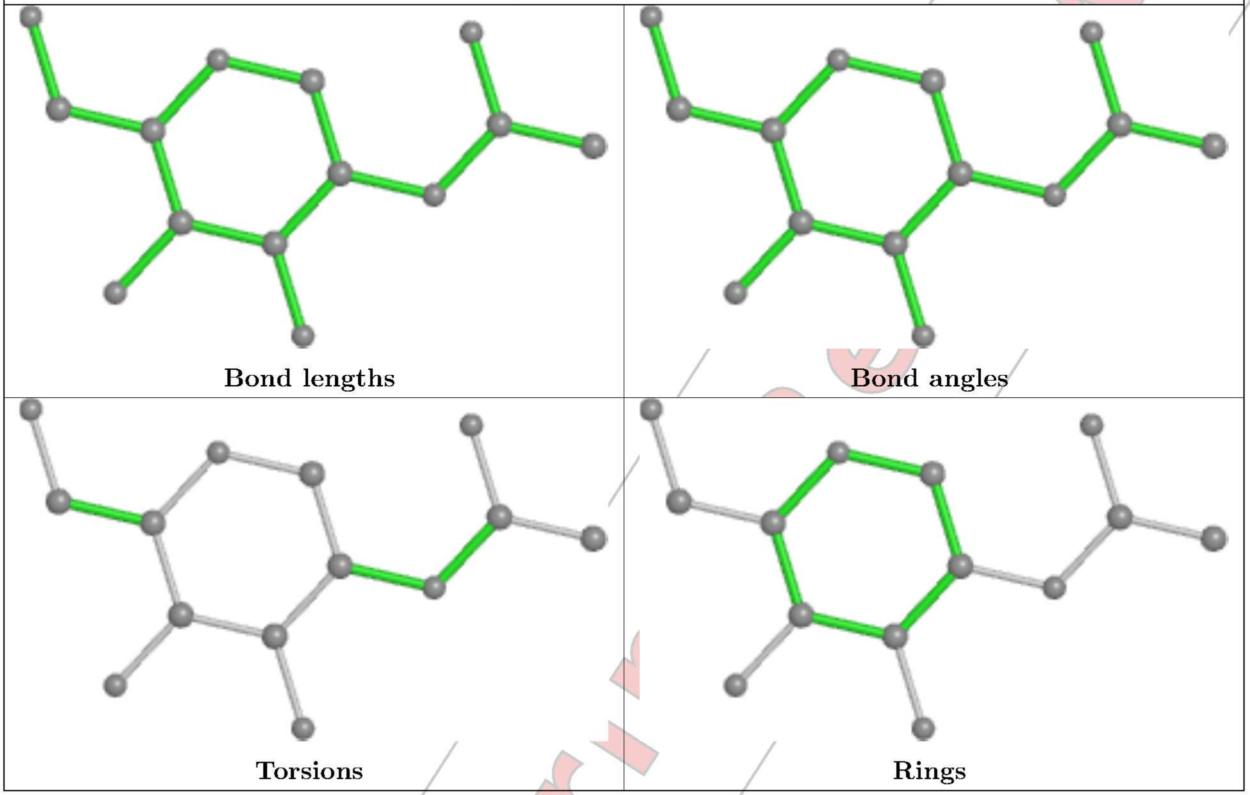


#### Ligand NAG U 501

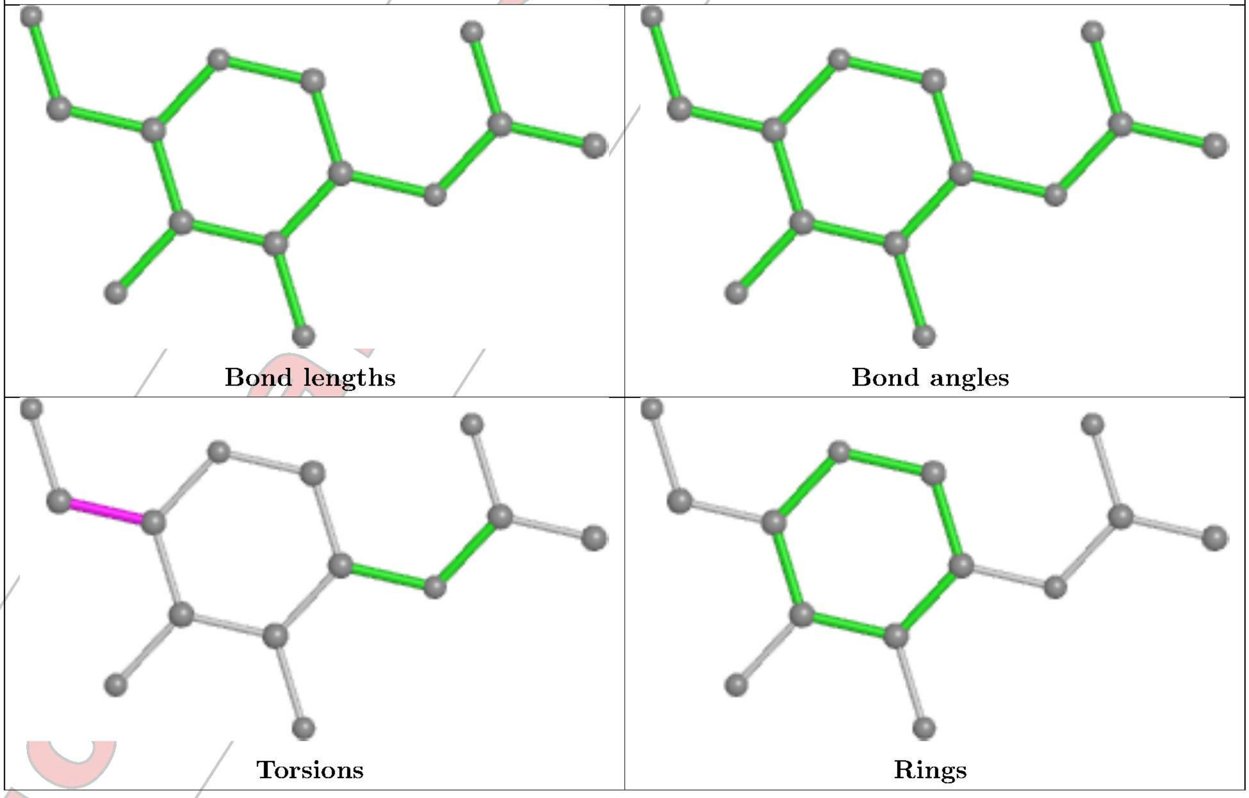

**Ligand NAG T 503**

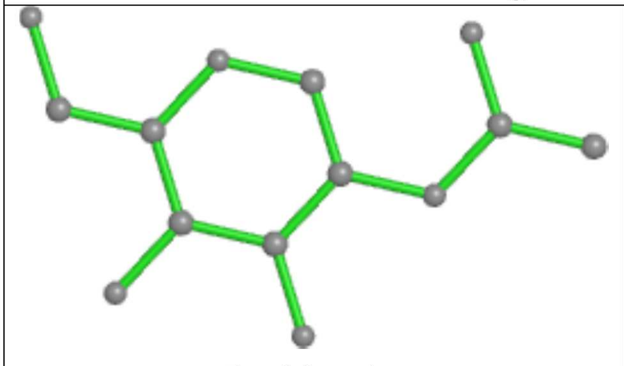

**Bond lengths**

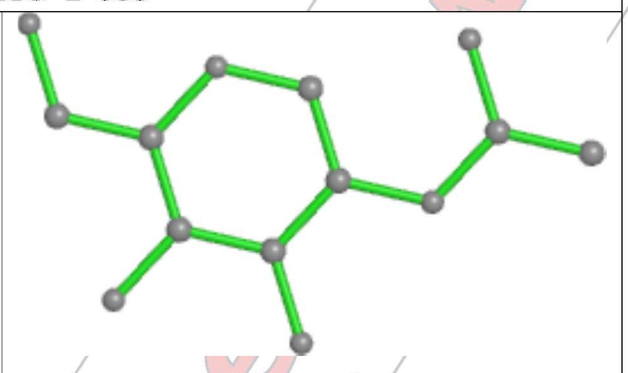

**Bond angles**

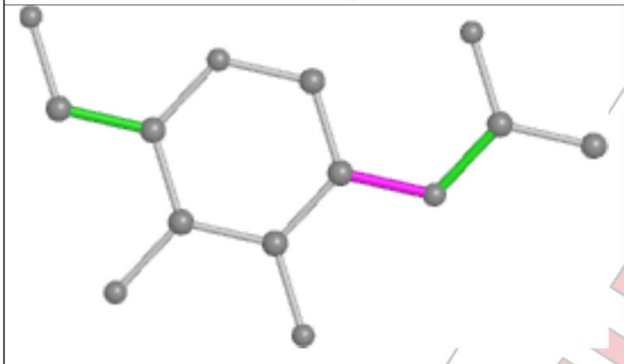

**Torsions**

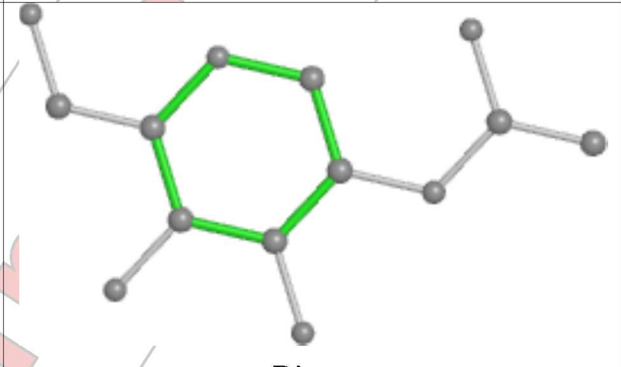

**Rings**

#### 5.7 Other polymers ⓘ

There are no such residues in this entry.

#### 5.8 Polymer linkage issues ⓘ

There are no chain breaks in this entry.

For Manuscript Review

### 6 Map visualisation (i)

This section contains visualisations of the EMDB entry EMD-26005. These allow visual inspection of the internal detail of the map and identification of artifacts.

Images derived from a raw map, generated by summing the deposited half-maps, are presented below the corresponding image components of the primary map to allow further visual inspection and comparison with those of the primary map.

#### 6.1 Orthogonal projections (i)

##### 6.1.1 Primary map

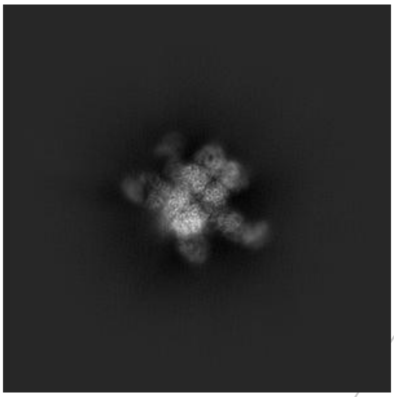

X

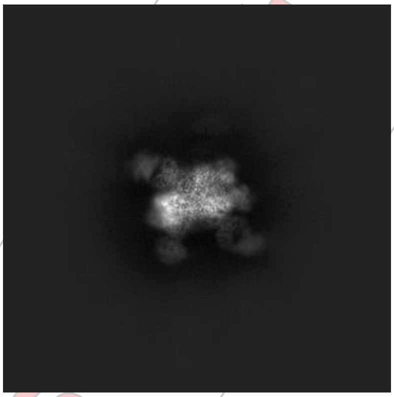

Y

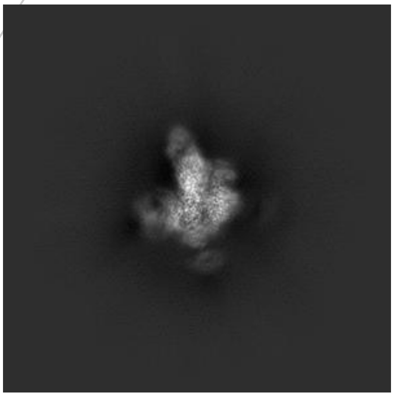

Z

##### 6.1.2 Raw map

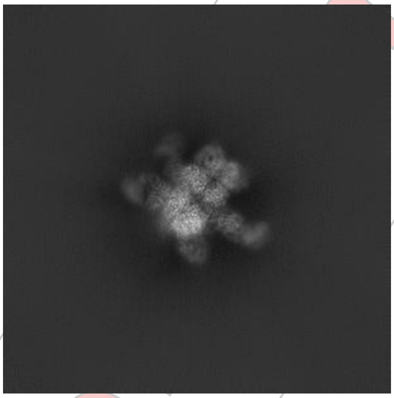

X

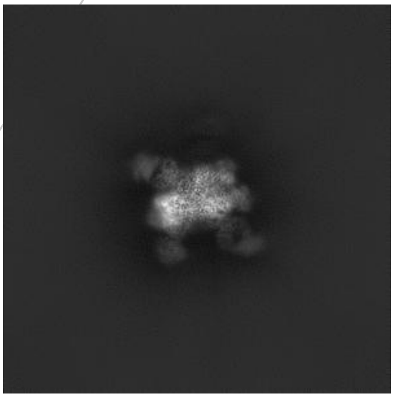

Y

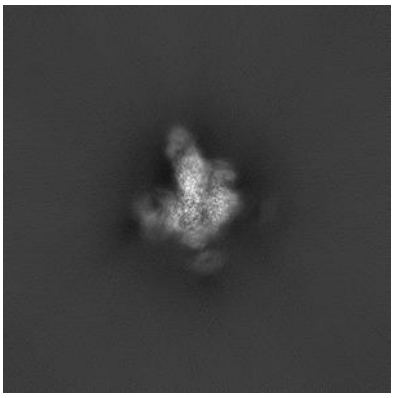

Z

The images above show the map projected in three orthogonal directions.

For Manuscript Review

#### 6.2 Central slices (i)

##### 6.2.1 Primary map

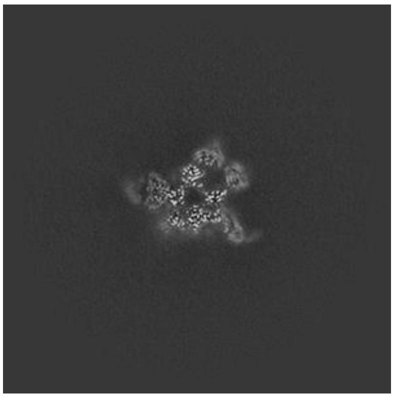
X Index: 256

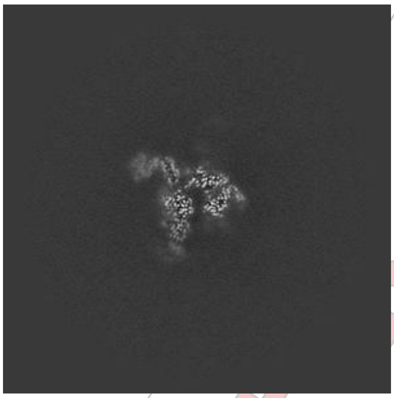
Y Index: 256

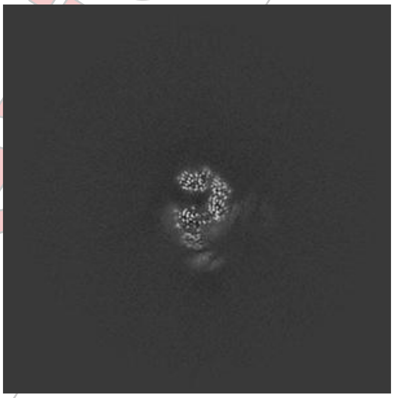
Z Index: 256

##### 6.2.2 Raw map

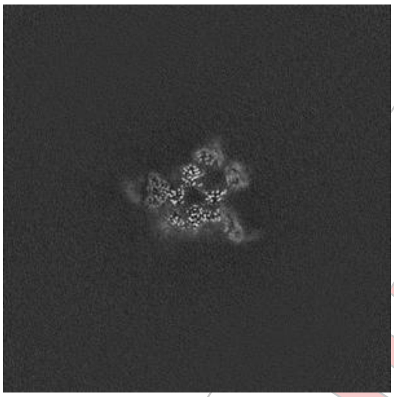
X Index: 256

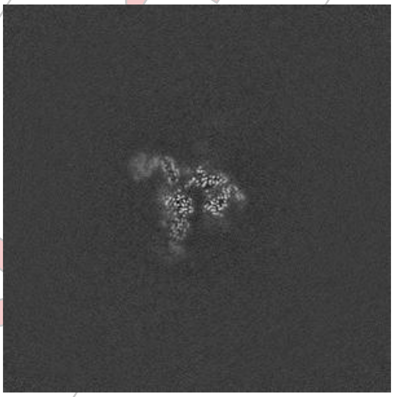
Y Index: 256

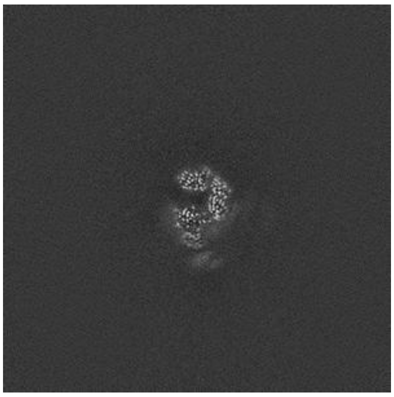
Z Index: 256

The images above show central slices of the map in three orthogonal directions.

For Manuscript Review

#### 6.3 Largest variance slices (i)

##### 6.3.1 Primary map

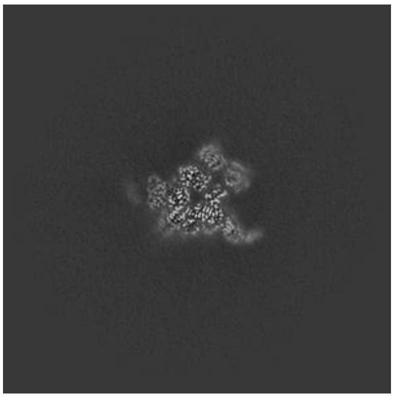

X Index: 251

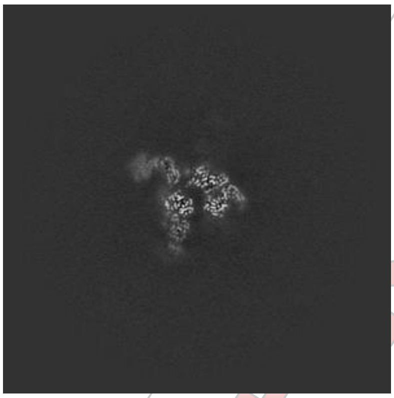

Y Index: 257

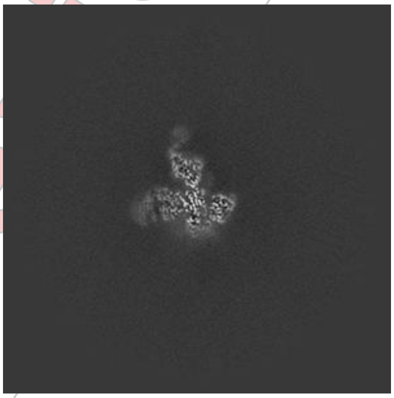

Z Index: 226

##### 6.3.2 Raw map

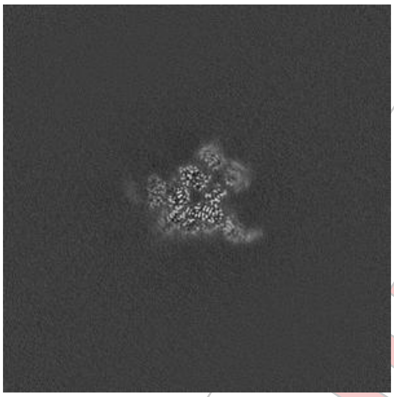

X Index: 251

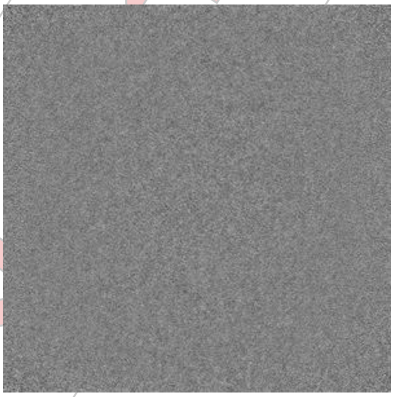

Y Index: 0

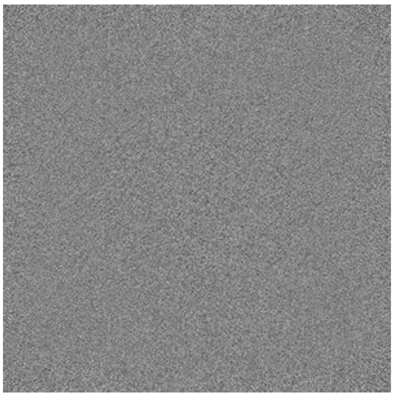

Z Index: 0

The images above show the largest variance slices of the map in three orthogonal directions.

For Manuscript Review

#### 6.4 Orthogonal surface views (i)

##### 6.4.1 Primary map

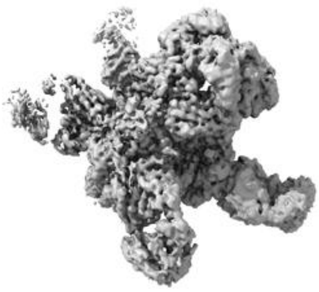
X

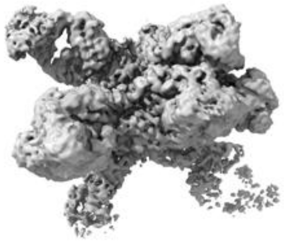
Y

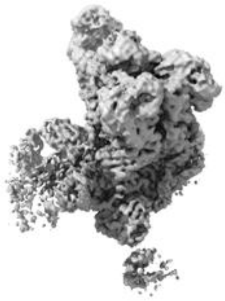
Z

The images above show the 3D surface view of the map at the recommended contour level 0.25. These images, in conjunction with the slice images, may facilitate assessment of whether an appropriate contour level has been provided.

##### 6.4.2 Raw map

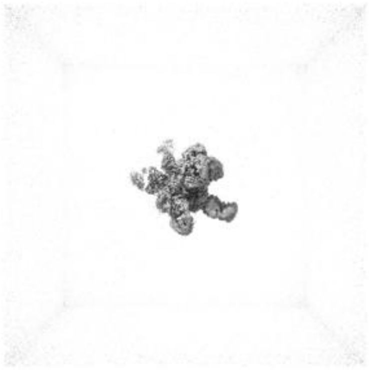
X

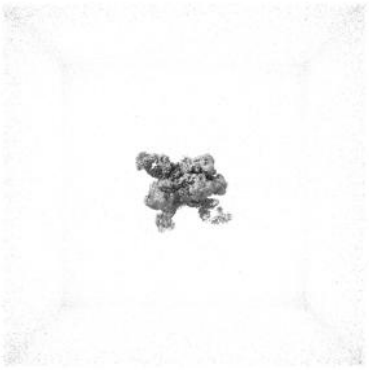
Y

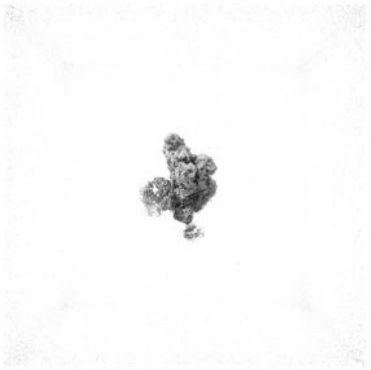
Z

These images show the 3D surface of the raw map. The raw map's contour level was selected so that its surface encloses the same volume as the primary map does at its recommended contour level.

#### 6.5 Mask visualisation (i)

This section was not generated. No masks/segmentation were deposited.

For Manuscript Review

### 7 Map analysis (i)

This section contains the results of statistical analysis of the map.

#### 7.1 Map-value distribution (i)

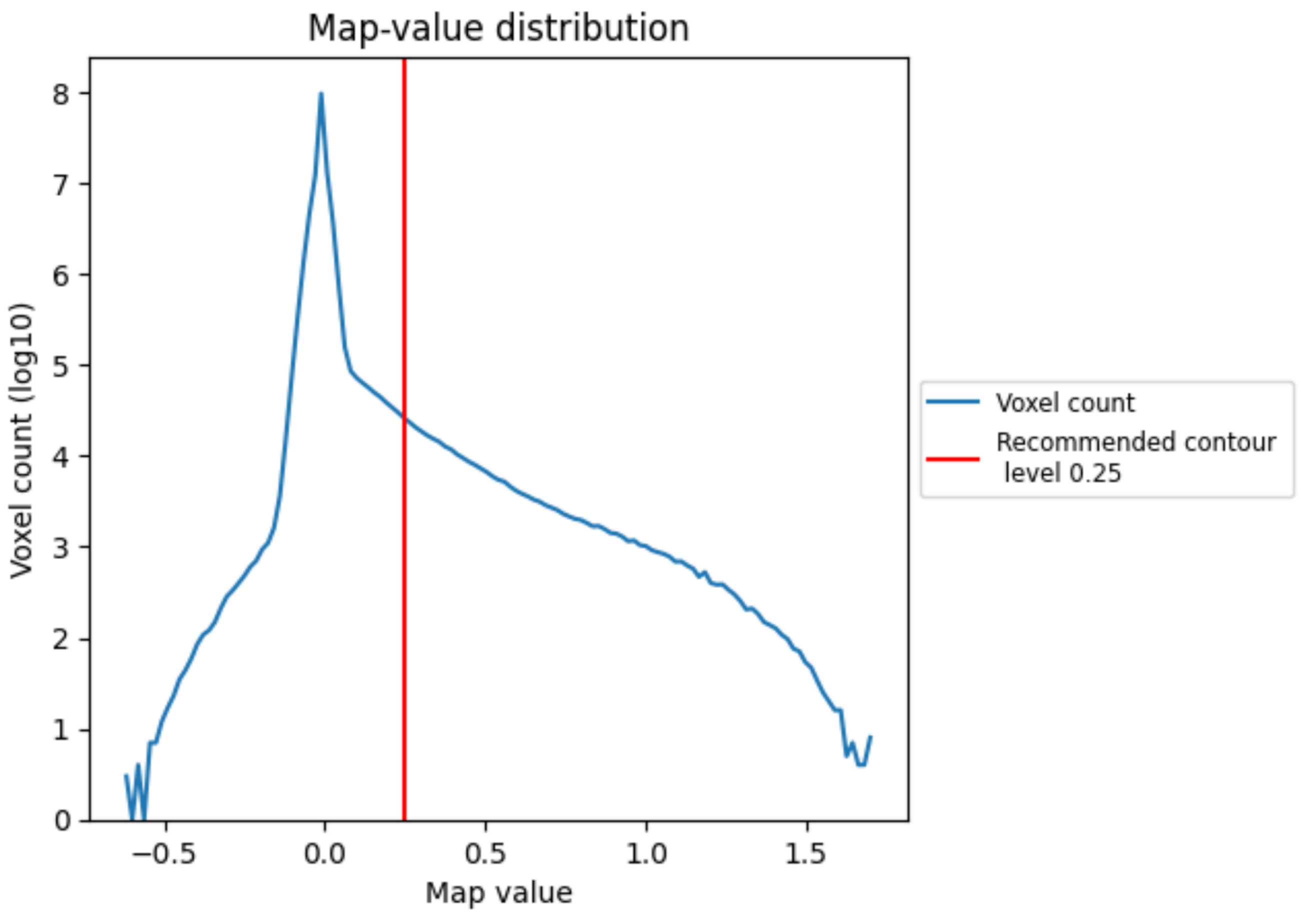


The map-value distribution is plotted in 128 intervals along the x-axis. The y-axis is logarithmic. A spike in this graph at zero usually indicates that the volume has been masked.

For Manuscript Review

#### 7.2 Volume estimate (i)

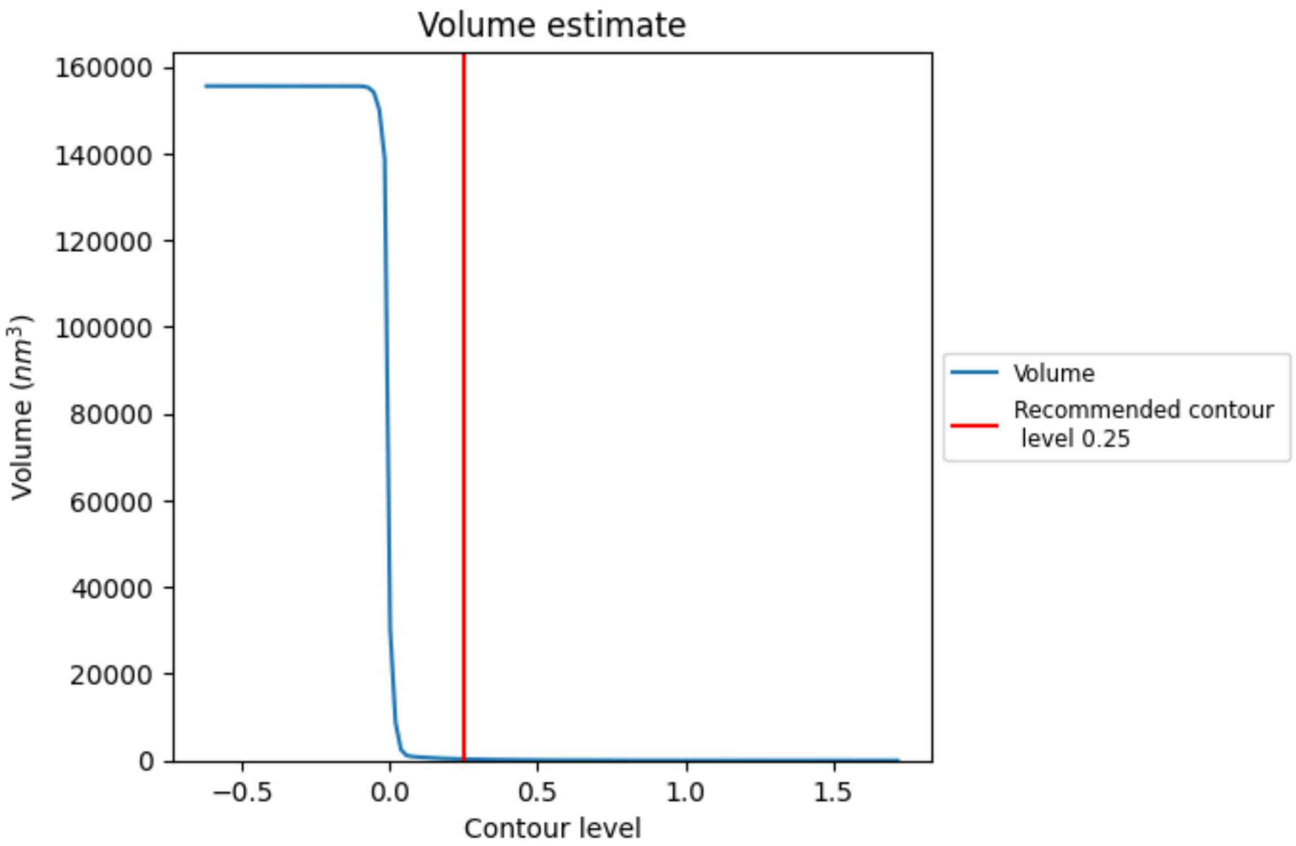


The volume at the recommended contour level is 332 $nm^3$; this corresponds to an approximate mass of 300 kDa.

The volume estimate graph shows how the enclosed volume varies with the contour level. The recommended contour level is shown as a vertical line and the intersection between the line and the curve gives the volume of the enclosed surface at the given level.

For Manuscript Review

#### 7.3 Rotationally averaged power spectrum ⓘ

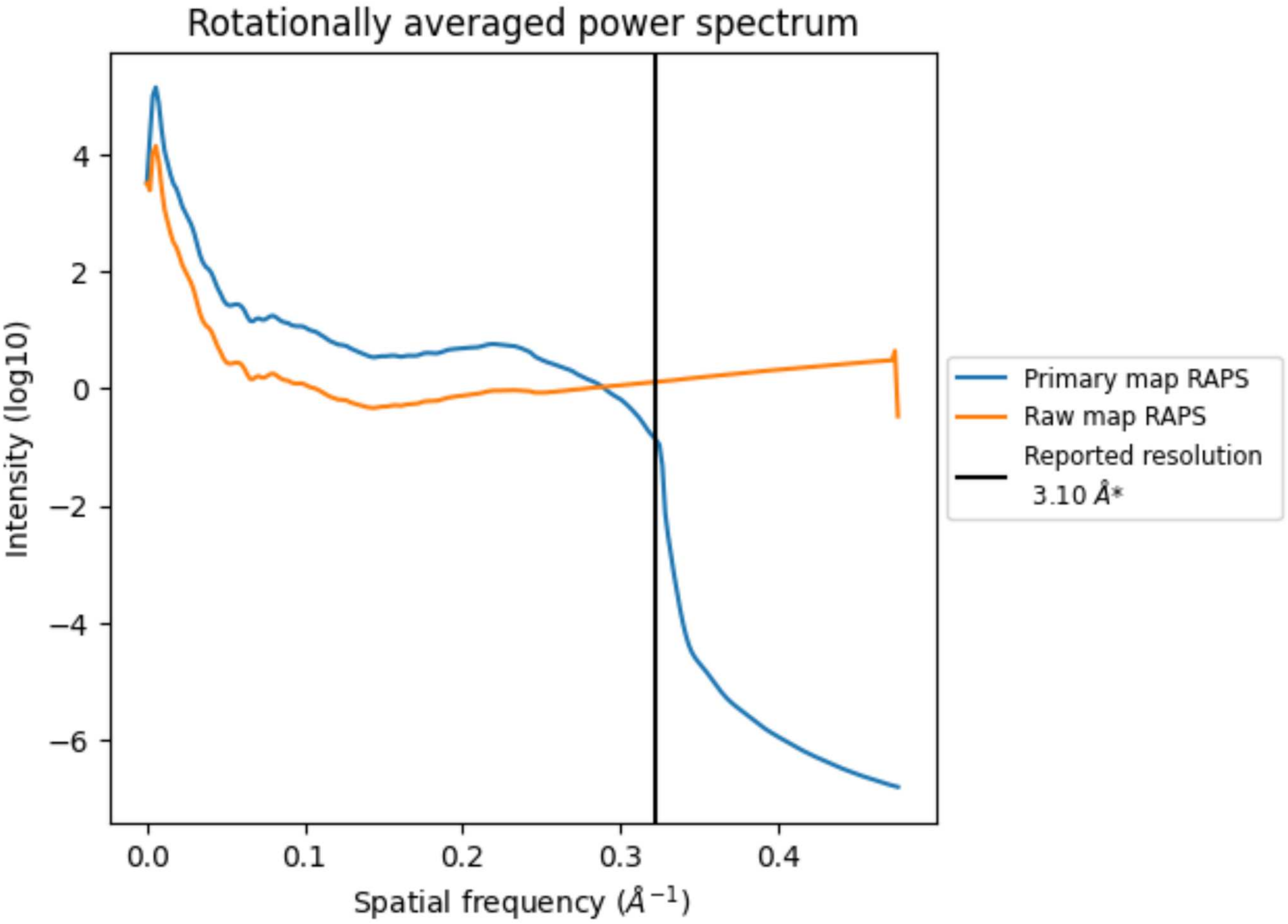


*Reported resolution corresponds to spatial frequency of 0.323 Å$^{-1}$

For Manuscript Review

### 8 Fourier-Shell correlation (i)

Fourier-Shell Correlation (FSC) is the most commonly used method to estimate the resolution of single-particle and subtomogram-averaged maps. The shape of the curve depends on the imposed symmetry, mask and whether or not the two 3D reconstructions used were processed from a common reference. The reported resolution is shown as a black line. A curve is displayed for the half-bit criterion in addition to lines showing the 0.143 gold standard cut-off and 0.5 cut-off.

#### 8.1 FSC (i)

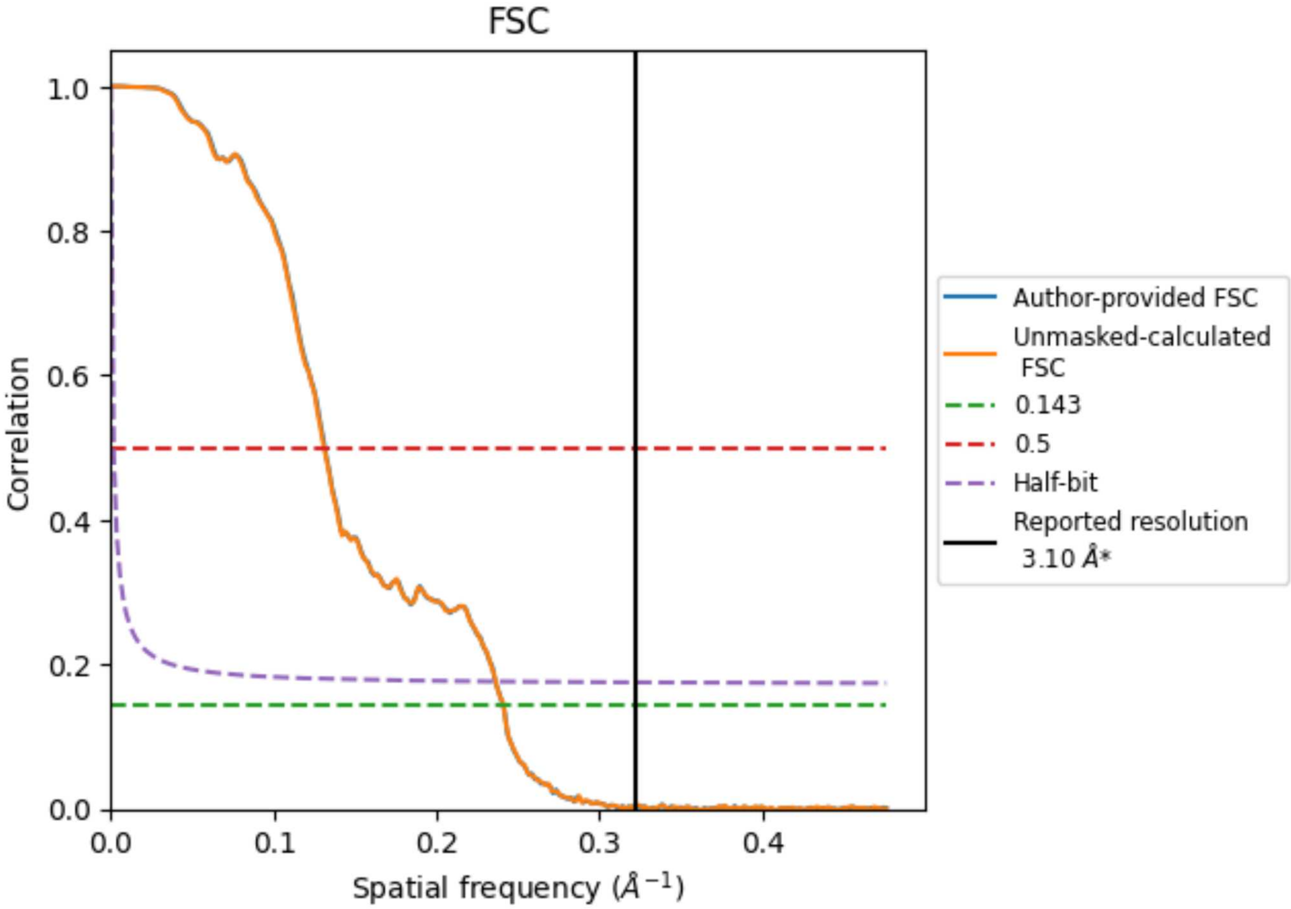


*Reported resolution corresponds to spatial frequency of 0.323 Å$^{-1}$

For Manuscript Review

#### 8.2 Resolution estimates (i)

| Resolution estimate (Å) | Estimation criterion (FSC cut-off) | | |
|---|---|---|---|
| | 0.143 | 0.5 | Half-bit |
| Reported by author | 3.10 | - | - |
| Author-provided FSC curve | 4.15 | 7.61 | 4.24 |
| Unmasked-calculated* | 4.15 | 7.65 | 4.23 |

*Resolution estimate based on FSC curve calculated by comparison of deposited half-maps. The value from author-provided FSC intersecting FSC 0.143 CUT-OFF 4.15 differs from the reported value 3.1 by more than 10 %

The value from deposited half-maps intersecting FSC 0.143 CUT-OFF 4.15 differs from the reported value 3.1 by more than 10 %

For Manuscript Review

### 9 Map-model fit (i)

This section contains information regarding the fit between EMDB map EMD-26005 and PDB model 7TN9. Per-residue inclusion information can be found in section 3 on page 8.

#### 9.1 Map-model overlay (i)

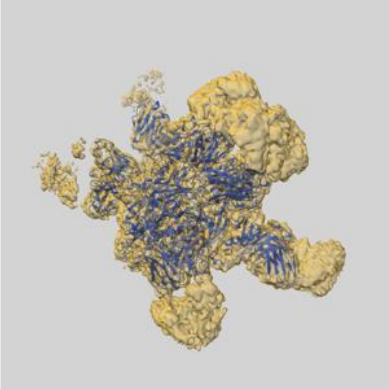

X

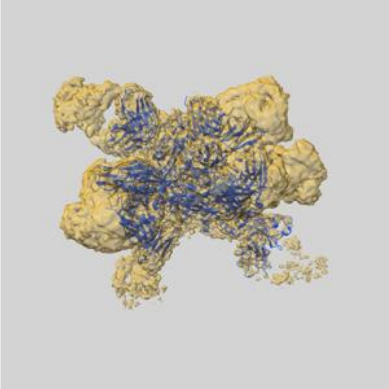

Y

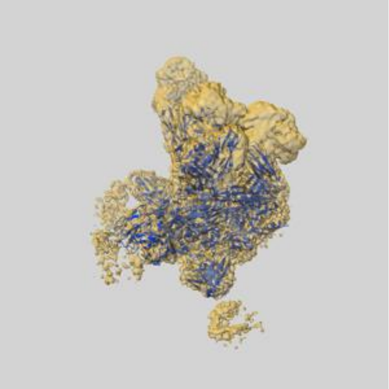

Z

The images above show the 3D surface view of the map at the recommended contour level 0.25 at 50% transparency in yellow overlaid with a ribbon representation of the model coloured in blue. These images allow for the visual assessment of the quality of fit between the atomic model and the map.

For Manuscript Review

#### 9.2 Atom inclusion (i)

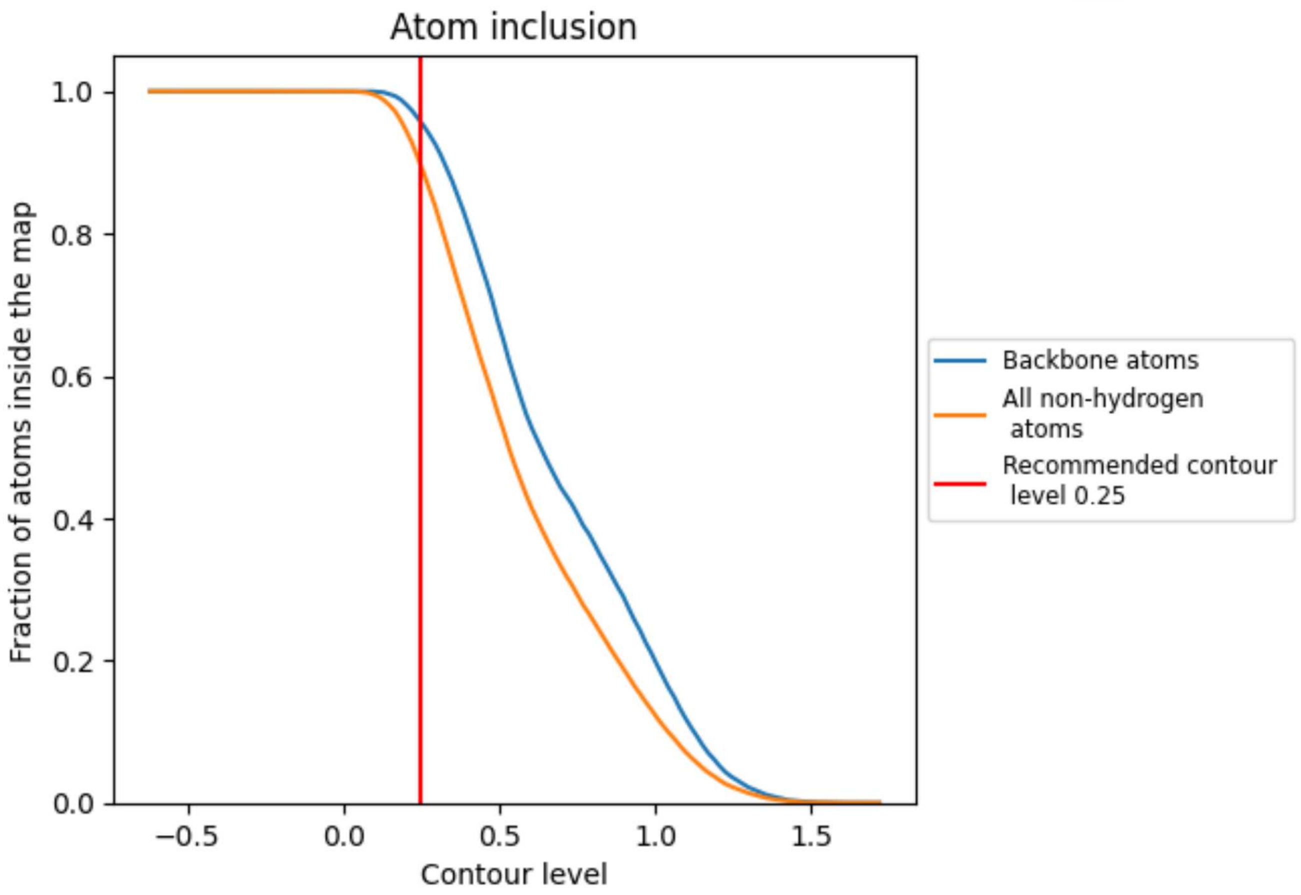


At the recommended contour level, 96% of all backbone atoms, 89% of all non-hydrogen atoms, are inside the map.

For Manuscript Review
